# Coordinated regulation of trichome morphogenesis and flavonoid pathway by a MYB–HDZIP–JAZ module in banana (*Musa* sp.)

**DOI:** 10.1101/2025.07.03.662943

**Authors:** Samar Singh, Shivi Tyagi, Prashant Misra, Ashutosh Pandey

## Abstract

- Trichomes are epidermal structures that function in defence against biotic and abiotic stresses. The developmental and metabolic aspects of trichomes have not been studied in banana (*Musa acuminata*). In the present study, we report that banana cultivars with AAA genome contain trichomes on inflorescence stalk.
- Transcriptome analysis of epidermal tissues of trichome-rich and trichome-free banana cultivars suggested that genes concerning defence response and specialized metabolism are differentially regulated. Based on the gene expression and phylogenetic analysis, two transcription factors, MaTFR (trichome and flavonol related R2R3-MYB) and MaHDZIV6 (HDZIP-IV), were identified as candidate regulators of trichome development.
- Both genes partially complemented the glabrous phenotype of trichome-related mutants of Arabidopsis. Transient overexpression and silencing in banana embryogenic cells (ECS) suggested that both MaTFR and MaHDZIV6 regulate the transcription of flavonol biosynthesis genes and modulate flavonoids content. Furthermore, MaTFR was found to directly activate the expression of *MaHDZIV6*.
- Trichome-enriched cultivars have enhanced active JA content compared to trichome-free cultivar suggesting the involvement of JA signalling in regulating trichome development. Two JAZ repressors, MaJAZ5 and MaJAZ8 interacted with MaTFR and attenuated the flavonoid biosynthesis. Our study concludes that the MYB, HDZIP-IV, and JAZ module fine-tunes the flavonoid biosynthesis and the trichome development in banana.

## Introduction

Trichomes are hair-like structures that emerge from epidermal cells and cover the aerial parts of plants. They are typically categorized into glandular trichomes (GTs) and non-glandular trichomes (NGTs), based on their structure and function and can be unicellular or multicellular (Watts & Kariyat, 2021; Feng *et al*., 2023). Trichomes serve as the first line of defense for plants, protecting them from insects, pathogens, and herbivores, and minimizing damage from UV radiation, low temperatures, and excessive water loss (Yan *et al*., 2012; Oksanen, 2018; Amada *et al*., 2023; Kabir *et al*., 2024). GTs are of particular interest as they play a central role in the biosynthesis and storage of a wide range of specialized metabolites, including alkaloids, flavonoids, phenylpropanoids, terpenoids, methyl ketones, and acyl sugars (Schilmiller *et al*., 2012; Li *et al*., 2022; Sugimoto *et al*., 2022; Feng *et al*., 2023). These metabolites are highly valued in pharmaceuticals, flavors, fragrances, and pesticides. For obvious reasons, there have been growing interest in understanding gene expression in GTs and molecular basis of their development in diverse plant species (Chalvin *et al*., 2020; Schuurink & Tissier, 2020).

The development of unicellular trichomes of *Arabidopsis thaliana* is well-studied, and several molecular regulators, governing their initiation, and patterning have been identified (Pattanaik *et al*., 2014; Arteaga *et al*., 2021; Méndez-Vigo *et al*., 2025). To this end, studies have suggested that an MYB–bHLH–WDR (MBW) complex of three regulatory proteins, namely GL1 (GLABRA 1), GL3 (GLABRA 3), EGL3 (ENHANCER OF GLABRA 3), and TTG1 (TRANSPARENT TESTA GLABRA 1) function as a core regulator of trichome development in Arabidopsis (Yang & Ye, 2013). Besides, transcription factors (TFs) belonging to other families that modify the activity of the core MYB–bHLH–WDR (MBW), have also been identified. Taken together, the complex interplay of these regulators fine-tunes the trichome development under spatial and temporal cues (Han *et al*., 2022; Dong *et al*., 2023). As compared to the unicellular trichome development, systemic information about the molecular basis of GTs development remains scarce. Recent studies on plants such as *Solanum lycopersicum*, *Nicotiana tabacum*, *Cucumis sativus*, and *Artemisia annua*, however, have been adding to the knowledge base of GT development. The TFs belonging to MYB, HD-ZIP, WRKY, MADS and Zn-Finger families have been identified to be involved in the development of GTs (Li *et al*., 2009; Matías Hernández *et al*., 2017; Xie *et al*., 2021; Yuan *et al*., 2024). Further, based on these studies, it is apparent that the molecular mechanism of GT development differs from that of unicellular trichomes.

Certain R2R3-MYB subfamily members, like MIXTA and MIXTA-like genes, regulate specialized metabolism and epidermal cell fate, including GT development (Chezem & Clay, 2016). MIXTA TFs are positive regulators of development of multicellular trichomes and GT in several plant species, such as *Antirrhinum majus*, *S. lycopersicum* and *A. annua* (Brockington *et al*., 2013). Other types of MYB TFs have also been implicated in the GT development. For example, in *A. annua*, TLR1, an R2R3 MYB TF, negatively impacts trichome growth. *A. annua* overexpressing TLR1 had lower trichome density and reduced artemisinin content, while RNAi suppression of TLR1 increased both trichome density and artemisinin content, compared to WT plants (Lv *et al*., 2022).

The plant-specific HD-ZIP IV subfamily of TFs are involved in the regulation of plant epidermal cell differentiation, cuticle biosynthesis, trichome and stomatal development (Yang *et al*., 2011; Nadakuduti *et al*., 2012; Schrick *et al*., 2023). In Arabidopsis, for example, GL2, an HDZIP IV family of TF positively regulates unicellular trichome development (Khosla *et al*., 2014). GaHOX1, a GL2-like gene, is predominantly expressed in cotton fiber cells, which functionally complement the *gl2* Arabidopsis mutant suggesting its role in cotton fiber development (Guan *et al*., 2008). In tomato, Wooly, another HDZIP IV family TF is involved in the positive regulation of GT development (Yang *et al*., 2011). Likewise, two HDZIP IV subfamily members, AaHD1 and AaHD8 function as positive regulators of GT development in *A. annua* (Yan *et al*., 2018). In *C. sativus*, another HDZIP IV subfamily gene, namely, Tril (Trichome-less) promotes the initiation of trichomes and fruit spine (Zhang *et al*., 2021).

Trichome-bearing tissues are known to experience reduced photodamage, highlighting their protective role against environmental stress. This protective effect is largely attributed to the accumulation of anthocyanins and other polyphenols in trichomes, which enhance defense against UV radiation (Saxena *et al*., 2023). In our previous study, we reported that GTs in *Cicer arietinum* accumulate flavonols and their derivatives, including kaempferol and quercetin glycosides, suggesting a metabolic specialization toward flavonoid biosynthesis (Saxena *et al*., 2023). Similarly, the metabolic activity of type-VI GTs (tVI-GTs) in *S. lycopersicum* is dominated by the synthesis and accumulation of flavonoids (Sugimoto *et al*., 2022). A recent study also revealed that *C. sativus* GTs accumulate flavonoids, underscoring the conserved role of trichomes in flavonoid-mediated stress tolerance (Feng *et al*., 2023). Flavonoids, classified into flavonols, flavones, flavanones, isoflavones, anthocyanins, and proanthocyanidins, play key roles in plant growth and stress responses (Halbwirth, 2010; Lago *et al*., 2014; Maloney *et al*., 2014; Chen *et al*., 2019; Yang *et al*., 2021; Saxena *et al*., 2023). Flavonols, with a 3-hydroxyflavone backbone, contribute to UV protection, ROS scavenging, and auxin transport regulation, influencing root and pollen development (Ko *et al*., 2014; Kitamura *et al*., 2016; Muhlemann *et al*., 2018; Liang *et al*., 2020; Chapman & Muday, 2021). Flavonols are produced through the flavonoid biosynthesis pathway, which involves several key enzymes such as chalcone synthase (CHS), chalcone isomerase (CHI), flavanone 3-hydroxylase (F3H), flavonoid 3′-hydroxylase (F3′H), flavonoid 3′5′-hydroxylase (F3′5′H), and flavonol synthase (FLS). These enzymes have been well studied in various plant species (Nagamatsu *et al*., 2007; Jiang *et al*., 2020).

Phytohormones play vital role in plant development and modulation of specialized metabolites (Pal *et al*., 2023; Gasperini & Howe, 2024). For example, Jasmonic acid (JA) plays diverse roles in vegetative growth, plant development, senescence, and defense mechanisms (Huang *et al*., 2017). It promotes the initiation of outgrowths on leaves and works synergistically with other hormones to regulate trichome formation and determine their density on various plant organs (Hua *et al*., 2022). The canonical JA signalling pathway relies on Jasmonate ZIM (JAZ) proteins, which function as repressors of downstream JA signaling TFs. JAZ proteins are targeted for degradation via ubiquitination mediated by CORONATINE INSENSITIVE1 (COI1) in response to JA (Chini *et al*., 2007; Pauwels & Goossens, 2011). For instance, AtJAZ1, AtJAZ8, and AtJAZ11 interact with AtMYB21 and AtMYB24 to regulate stamen development in Arabidopsis (Song *et al*., 2011). NbJAZ3 reduces trichome genesis in tobacco by targeting the NbWo, which was further shown to suppress the transcriptional activation of NbCycB2 (Yan *et al*., 2022). In cotton (*Gossypium sp*.), GhJAZ2 negatively impacts fiber initiation and elongation by interacting with GhMYB25-like (Hu *et al*., 2016). Treatment of Methyl JA has been shown to promote fiber elongation in cotton ovules (Xia *et al*., 2018) . In tomato, overexpression (OE) of *SlJAZ2* led to reduced plant height, shortened internode length, and decreased trichome density (Yu *et al*., 2018). These findings indicate that JAZ proteins often serve as negative regulators in organ morphogenesis including trichome development.

Trichomes have also been reported to be present in monocot plant species. For example, in rice, three distinct types of trichomes, namely micro, macro, and glandular hairs have been reported (Sturaro *et al*., 2005; Zheng *et al*., 2016). A few transcriptional regulators, such as OsWOX3B, HL6, and OsSPL10 have been implicated in the developmental regulation of trichomes in rice (Li *et al*., 2012; Zhang *et al*., 2012; Lan *et al*., 2019; Liao *et al*., 2023). Like rice, trichomes in maize are also diversified and represented by macro hairs, prickle hairs and bicellular micro hairs (Moose *et al*., 2004). An HDZIP IV subfamily TF OCL4 and a SPL family TF, have been shown to play a key role in trichome development in maize (Vernoud *et al*., 2009; Kong *et al*., 2021). Despite the knowledge about these TFs, molecular basis of trichome development remains poorly understood in monocots.

Banana, a monocot plant species is one of the most important fruit crops. The global production of banana is significantly impacted by various stresses, such as drought, cold, salinity, and pathogen attack (Xu *et al*., 2020). Previous studies have shown that B-genome containing cultivars like Plantain (PLT) are tolerant against *Fusarium oxysporum* f. sp. *cubense* 4 (Foc4) causing severe wilt disease, *Xanthomonas* infection, and drought stress (Vanhove *et al*., 2012; Davey *et al*., 2013). However, Grand Naine (GN) and Red Banana (RB) having AAA-genome are susceptible to various abiotic and biotic stresses (Tripathi *et al*., 2008). Trichomes have been implicated in conferring tolerance against biotic and abiotic stress. However, to the best of our knowledge, till date, no systematic study has been conducted to study the molecular aspects of trichome biology in banana.

Against this backdrop, in the present study, we firstly reported the presence of trichomes on the fruit stalks of GN and RB cultivars and performed a detailed microscopic and histological characterization of banana trichomes. The presence of trichomes on the fruit stalks of these cultivars may be an evolutionary adaptation in AAA genome cultivars to protect its inflorescence against stresses. Transcriptome analysis of trichome-free epidermis of PLT and trichome-rich (TR) epidermis of GN and RB led to the identification of MaTFR and MaHDZIV6 as key regulators of trichome development and flavonol biosynthesis. MaTFR and MaHDZIV6 were found to interact at both protein–protein and protein–DNA levels, reinforcing their regulatory relationship. JA quantification and methyl JA treatment confirmed the involvement of JA signalling in trichome development. Notably, MaJAZ5 and MaJAZ8 were shown to interact with MaTFR, reducing its transcriptional activity and thereby modulating the expression of flavonoid biosynthetic genes. Our study suggests that the MaTFR–MaHDZIV6 module is tightly regulated by JA signalling, with JAZ proteins acting as negative regulators under stress or developmental cues. Overall, the MaTFR–MaHDZIV6–JAZ regulatory module emerges as a central player in coordinating trichome development and flavonoid biosynthesis in banana.

## Materials and methods

### Plant materials and growth conditions

Banana cultivars GN, RB and PLT were grown at the field of the National Institute of Plant Genome Research (NIPGR), New Delhi. The planting materials were collected from the NIPGR field. To study the trichomes, the epidermal layers were carefully peeled from the inflorescence stalk of GN, RB and PLT using fine forceps under a stereomicroscope. The isolated epidermis was then cut into small segments and mounted directly onto clean glass slides without any staining or fixation. The samples were subsequently observed under a stereo microscope to document trichome morphology and distribution.

*Arabidopsis thaliana* Columbia-0 (Col-0) wild-type, the *gl1* and *gl2* mutants (TAIR stock; AT3G27920.1 and AT1G79840.1, respectively both in the Ler background), promoter-reporter lines (*proMaTFR::GUS* and *proMaHDZIV6::GUS*), and transgenic Arabidopsis lines overexpressing *MaTFR* and *MaHDZIV6* were grown in a controlled growth chamber (Percival AR-41L3, Perry, IA, USA) under a 16 h light / 8 h dark photoperiod, with a light intensity of 100 µmol m ² s ¹ at a constant temperature of 22°C *N. benthamiana* plants were grown under the same controlled conditions as described for *A. thaliana*. Embryogenic cell suspension (ECS) cultures derived from immature male flower buds were used for transient overexpression and RNA interference (RNAi) experiments. Six-week-old banana seedlings (GN cultivar) were used for methyl JA treatment. The seedlings were grown under controlled conditions in a plant growth chamber (Model: AR-41L3, Percival) set at 26 °C with a 16-hour light/8-hour dark photoperiod, light intensity of 250 μmol m ² s ¹, and 60% relative humidity. Plants were treated either with a mock solution or with 100 µM methyl JA (Sigma Aldrich). Leaf samples were harvested at six time points (0, 6, 12, 24, 48, and 72 h) post-treatment, immediately frozen in liquid nitrogen, and stored at –80 °C until further analysis.

### Morphological characterization of trichomes through stereo microscopy and scanning-electron microscopy

The epidermal layers of inflorescence stalks from GN, RB, and PLT were examined for trichome morphology using scanning electron microscopy (SEM) (EVO LS10; Carl Zeiss). For *Arabidopsis thaliana* (Col-0), promoter-GUS lines and overexpressing lines, were grown on half-strength Murashige and Skoog (½ MS) medium. T generation overexpression lines of *MaTFR* and *MaHDZIV6* were also subjected to SEM (EVO LS10; Carl Zeiss) to assess trichome morphology and to quantify trichome density.

### Metabolite Localization in Trichomes

To identify flavanols in trichomes, a 0.25% (w/v) DPBA staining solution (Sigma Aldrich) was prepared in Milli-Q water, supplemented with 0.00375% (v/v) Triton X-100 (Naik *et al*., 2021). Approximately 2 mL of the freshly prepared DPBA solution was transferred to a sterile Petri plate, and small segments of epidermal tissue were immersed for 10 minutes. After staining, the tissues were rinsed twice with Milli-Q water and mounted on glass slides (Nguyen, 2020). Samples were then examined using a confocal laser scanning microscope to visualize flavanol-specific fluorescence signals.

For lipid content analysis, the epidermal sections were immersed in a saturated solution of freshly prepared Sudan IV dye (Sigma Aldrich) for 15 minutes. The stained tissues were subsequently rinsed twice with 70% ethanol to remove excess dye. Lipid accumulation in trichomes was visualized by observing red or orange coloration under a stereomicroscope using a brightfield filter (Nikon SMZ25 with DS-Fi3 camera, Tokyo, Japan). Terpenoids were detected using the NADI staining reagent, a mixture of α-naphthol (Sigma Aldrich) and N, N-dimethyl-p-phenylenediamine hydrochloride (Sigma Aldrich). A 10 mL aliquot of the freshly prepared NADI reagent was poured into a Petri plate containing thin slices of epidermal tissue. After adequate staining, the samples were examined under a stereomicroscope using a brightfield filter to detect terpenoid-associated coloration (Serrato-Valenti *et al*., 1997).

### Ultra-high-performance liquid chromatography and liquid chromatography–mass spectrometry for detection of targeted flavonoids and endogenous defense phytohormones

Targeted flavonoids metabolite profiling was performed following previously established protocols (Pandey *et al*., 2016; Saxena *et al*., 2023). Individual flavonoids were determined either as glycosylated forms or as aglycones by preparing non-hydrolyzed and acid-hydrolyzed extracts, respectively, as described earlier (Misra *et al*., 2010). For non-hydrolyzed extraction, plant material was finely shredded and homogenized in 80% methanol at room temperature. The homogenate was then heated at 70 °C for 15 minutes, followed by centrifugation at 12,000 rpm for 10 minutes. To prepare acid-hydrolyzed extracts, the resulting supernatant was mixed with three volumes of 1 N HCl and incubated at 94 °C for two hours. After hydrolysis, an equal volume of ethyl acetate was added, and the upper organic phase was collected for purification. The ethyl acetate was subsequently evaporated under reduced pressure (90– 100 mbar) at 37 °C using a Rotavapor R-300 (BUCHI, Flawil, Switzerland). The dried residue was reconstituted in 80% methanol prior to LC–MS analysis. Prior to UHPLC/LC–MS analysis, samples were filtered through a 0.22 µm Millipore syringe filter (Millipore, USA) to remove particulates.

Quantitative analysis of targeted flavonoids was performed using an Exion LC UPLC system (Sciex) coupled with a QTRAP 6500+ triple quadrupole mass spectrometer (AB Sciex), equipped with an electrospray ionization (ESI) source, as described earlier (Naik *et al*., 2021). The ionization voltage was set at 5500 V in positive mode. Source parameters included Gas 1 and Gas 2 at 70 psi, curtain gas at 40 psi, collision-assisted dissociation (CAD) set to medium, and a source temperature of 650 °C. The instrument was operated in multiple reaction monitoring (MRM) mode for both identification and quantification. Analytical standards were obtained from Merck, and data processing was performed using Analyst software (version 1.5.2). Defense-related phytohormones were quantified from TF (PLT) and TR (GN and RB) following the protocol by (Vadassery *et al*., 2012). 30 mg of lyophilized tissue was extracted in methanol containing internal standards. After centrifugation at 12,000 × g for 15 minutes at 4 °C, the supernatants were collected and dried using a speed vacuum concentrator. The dried residue was reconstituted in 500 µL methanol, filtered through a 0.22 µm Millipore syringe filter, and subjected to LC–MS analysis.

Quantification was performed on the same LC–MS system described above, using a UPLC C18 column (Exion LC, Sciex, Framingham, MA, USA) coupled to a QTRAP 6500+ mass spectrometer (AB Sciex) with ESI. The analysis was conducted in MRM mode for accurate quantification of hormone levels.

### RNA isolation and transcriptome profiling of TF and TR cultivars

The TF epidermis of PLT and TR epidermis of GN and RB cultivars were collected and immediately frozen in liquid nitrogen to preserve RNA integrity. Total RNA was extracted using the Spectrum™ Plant Total RNA Kit, following the manufacturer’s protocol. The quality and quantity of the extracted RNA were evaluated using agarose gel electrophoresis, a NanoDrop spectrophotometer, and an Agilent Bioanalyzer. To eliminate potential genomic DNA contamination, RNA samples were treated with RNase-free DNase using the TURBO DNA-free™ Kit (Invitrogen, USA). Only RNA samples with an RNA integrity number (RIN) of ≥6 were used for RNA-seq library preparation. Libraries were constructed for paired-end sequencing according to the Illumina protocol and subsequently sequenced using the Illumina NOVASEQ 6000 sequencing platform to generate transcriptomic data for downstream analyses.

High-throughput RNA sequencing data were processed using a standardized pipeline. Initially, raw sequencing reads in FASTQ format were assessed with FastQC at default parameters for quality assessment. Reads with low-quality bases and adapter sequences were trimmed using Fastp v0.20.1, followed by a second round of quality checks with FastQC to verify the integrity of the cleaned data. The processed reads were aligned to the reference genome using STAR (Spliced Transcripts Alignment to a Reference) v2.7 in paired-end mode. The reference genome and its corresponding annotation file in GTF format were obtained from Banana Genome Hub (https://banana-genome-hub.southgreen.fr/). Following alignment, gene counts were quantified using featureCounts v2.0.3 from the Subread package and gene expression levels were calculated as FPKM (Fragments Per Kilobase of transcript per Million mapped reads). The resulting count matrix was analyzed for differential expression using DESeq2 in R. Differentially expressed genes (DEGs) were identified in different combinations (PLT vs GN, PLT vs RB and GN vs RB) based on statistical significance (adjusted p-value < 0.05) and fold change thresholds. The volcano plots were generated for visualization of upregulated and downregulated DEGs. The venn diagram was made to identify unique and common DEGs in different combinations (PLT vs GN, PLT vs RB and GN vs RB) using Venny 2.1 (https://bioinfogp.cnb.csic.es/tools/venny/index.html). Heatmaps were generated using FPKM values by TBtools (Chen *et al*., 2020) to visualize the gene expression profiles. The upregulated and downregulated DEGs (PLT vs GN, PLT vs RB and GN vs RB) at padj value <0.05 and log2(fold change) ≥2 or ≤-2 were subjected to gene ontology (GO) enrichment analyses at Banana Genome Hub (https://banana-genome-hub.southgreen.fr/content/go-enrichment). The GO enrichment results were visualized by bubble plots generated using ggplot2 R package.

### Gene expression analysis

Total RNA was isolated from the epidermal layers of selected *Musa* cultivars (PLT, GN and RB) and *Arabidopsis thaliana* using the Spectrum™ Plant Total RNA Kit (Sigma-Aldrich), following the manufacturer’s instructions. To eliminate any residual genomic DNA contamination, the extracted RNA was treated with RNase-free DNase using the TURBO DNA-free™ Kit (Invitrogen, USA). First-strand cDNA synthesis was carried out using 1 µg of DNase-treated total RNA with the RevertAid H Minus First Strand cDNA Synthesis Kit (Thermo Fisher Scientific), employing oligo(dT) primers. Quantitative real-time PCR (RT-qPCR) was performed using a 7500 Fast Real-Time PCR System (Applied Biosystems) with SYBR Green detection. Each reaction (10 µL total volume) contained 2× SYBR Green PCR Master Mix (Applied Biosystems) and diluted cDNA corresponding to 10 ng of input total RNA. The *MaACTIN1* gene (GenBank accession AF246288) was used as an internal reference for normalization of transcript levels. Relative gene expression was calculated using the 2^−ΔΔCT^ method (Livak & Schmittgen, 2001). Data are presented as fold changes relative to the tissue with the lowest expression level or to the control condition, as appropriate. Each RT-qPCR analysis included three biological and three technical replicates.

### Identification of trichome-specific regulators

Phylogenetic analysis was conducted for trichome-related HD-ZIV, JAZ and MYB proteins, separately, to determine the evolutionary relationships among the protein sequences. The sequences were aligned using the MUSCLE algorithm in MEGA 11 software (Tamura *et al*., 2021). The phylogenetic tree was constructed using the Maximum Likelihood (ML) method. Bootstrap analysis with 1000 replicates was performed to assess the robustness of the tree topology. The tree was visualized and annotated using the iTOL v7 software (Letunic and Bork, 2024). The co-expression analysis was performed using RStudio with R version 4.4.2 and interaction network was generated using Cytoscape3.10.3 (Shannon *et al*., 2003).

### Cloning of transcriptional regulators

Full-length coding sequences (CDSs) of MaMYB-encoding gene (*MaTFR), MaHDZIV6*, and selected MaJAZ excluding the stop codons, were amplified using first-strand cDNA synthesized from the TR epidermal tissue of banana. Gene-specific primers containing Gateway™ attB recombination sites were designed based on sequence information from the Banana Genome Hub (see Table S1). The resulting PCR amplicons were recombined into the Gateway® entry vector pDONR™Zeo (Invitrogen) via BP recombination reaction and transformed into *Escherichia coli* DH5α (Invitrogen) competent cells. The integrity of the entry clones was confirmed by Sanger sequencing, performed at the NIPGR sequencing core facility (New Delhi, India).

### Subcellular localization and co-localization assay

Entry clones harboring CDSs of MaTFR, MaHDZIV6, were cloned into the pSITE-3CA binary vector and MaTFR in pSITE4NB vector via LR recombination to create yellow fluorescent protein (YFP) fusion constructs (Chakrabarty *et al*., 2007). Empty vectors served as negative controls. The resulting binary plasmids were transformed into Agrobacterium tumefaciens strain GV3101-pMP90 (Koncz & Schell, 1986). Equal volumes of *A. tumefaciens* cultures containing the target gene and marker were mixed for agro-infiltration. The abaxial sides of *N. benthamiana* leaves were infiltrated using a syringe. The infiltrated plants were kept in the dark at 25°C for 2 days before analysis. Leaf discs were excised from the infiltrated regions and examined using an argon laser confocal scanning microscope (TCS SP5, Leica Microsystems, Wetzlar, Germany) equipped with YFP and RFP filters. YFP fluorescence was detected at 514 nm excitation and 527 nm emission, while RFP fluorescence was detected at 558 nm excitation and 583 nm emission.

### Promoter isolation and reporter construct preparation

Putative promoter sequences of target genes were retrieved from the Banana Genome Hub (https://banana-genome-hub.southgreen.fr) based on *M. acuminata* genome annotations (D’hont *et al*., 2012). These sequences were analyzed for the presence of *cis*-regulatory motifs using the New PLACE database (https://www.dna.affrc.go.jp), with additional manual curation to validate key regulatory elements. Total genomic DNA was extracted from banana leaves using the DNeasy Plant Mini Kit (Qiagen). Promoter fragments were amplified from genomic DNA using gene-specific primers (listed in Table S1) and Phusion™ High-Fidelity DNA Polymerase (Thermo Scientific). The resulting amplicons were cloned into the pENTR™/D-TOPO™ entry vector (Thermo Fisher Scientific) and confirmed by Sanger sequencing. For the generation of *proMaTFR::GUS* and *proMaHDZIV6::GUS*, the confirmed promoter entry clones were recombined into the destination vector pKGWFS7 (Karimi *et al*., 2002). Verified promoter entry clones—such as *proMaHDZIV6-*1994, p*roMaFLS1*–2508, *proMaF3*′*5*′*H1*–1335, and *proMaF3*′*5*′*H2*–1335 were subsequently recombined into the destination vector p635nRRF containing the *pro35S::REN* reporter cassette (Kumar *et al*., 2018) via Gateway™ LR recombination reaction to generate dual-luciferase reporter constructs.

### Heterologous expression in Arabidopsis mutants and histochemical GUS staining

Entry clones containing the coding sequences of *MaTFR* and *MaHDZIV6* were recombined into the Gateway-compatible binary destination vectors pSITE-3CA (Chakrabarty *et al*., 2007). and pMDC32 (Curtis & Grossniklaus, 2003), respectively. The pSITE-3CA vector carries a kanamycin resistance gene, while pMDC32 harbors a hygromycin resistance gene. Both vectors feature a double *CaMV 35S* promoter to drive constitutive overexpression of the transgenes. The resulting expression plasmids were introduced into *A. tumefaciens* strain GV3101 (pMP90) via chemical transformation. Transformed Agrobacterium cultures were used to genetically transform *A. thaliana* trichome-less mutants *gl1-2 (AT3G27920.1)* and *gl2-1 (AT1G79840.1)* using the floral dip method (Clough & Bent, 1998). T transgenic seedlings were selected on Murashige and Skoog (MS) agar medium containing 50 mg/L kanamycin or 15 mg/L hygromycin, depending on the selectable marker used. Resistant seedlings were transferred to soil and grown under controlled growth conditions for seed production. Homozygous T lines were screened and subsequently evaluated for trichome phenotypes, and flavonol content.T transgenic Arabidopsis plants harboring *proMaTFR::GUS* or *proMaHDZIV6::GUS* constructs were subjected to histochemical GUS staining. Whole seedlings were vacuum-infiltrated with GUS staining buffer containing 50 mM sodium phosphate buffer (pH 7.0), 1 mM 5-bromo-4-chloro-3-indolyl-β-D-glucuronide (X-Gluc; cyclohexylammonium salt), 0.4 mM potassium ferricyanide, 0.4 mM potassium ferrocyanide, 2 mM EDTA, and 0.12% (v/v) Triton X-100. Infiltration was carried out for 20–30 minutes under vacuum. After infiltration, the seedlings were incubated at 37 °C in the dark for 2–3 days or until the GUS signal developed. Following staining, tissues were cleared with chloral hydrate solution, and residual chlorophyll was removed by repeated washing with 70% ethanol.

### Yeast two-hybrid (Y2H) assay

For yeast two-hybrid (Y2H) assays, the coding sequences of MaTFR, MaHDZIV6, MaJAZs were cloned into the prey vector pGADT7g or the bait vector pGBKT7g (Clontech Laboratories Inc., San Jose, CA, USA). The resulting plasmids were co-transformed into *Saccharomyces cerevisiae* strain Y2HGold (Clontech) and cultured at 30°C on a synthetic defined (SD) medium lacking tryptophan (Trp) and leucine (Leu). For interaction tests, yeast colonies carrying the respective bait and prey constructs were spotted onto SD medium lacking Trp, Leu, histidine (His), and adenine (Ade) (QDO), supplemented with 40 mg/ml X-Gal and 5 mM 3-amino-1,2,4-triazole (3-AT). The interaction between p53 and large T antigen served as a positive control, while empty pGADT7g and pGBKT7g vectors were used as negative controls (Pipas & Levine, 2001).

### Bimolecular fluorescence complementation assay

The entry clones carrying the coding sequences of MaTFR, MaHDZIV6, MaJAZ5 and MaJAZ8 were recombined into the vectors 35S-pSITE-nYFP-N1 or 35S-pSITE-cYFP-N1 (Martin *et al*., 2009). The resulting plasmids were individually transformed into *A. tumefaciens* strain GV3101 (pMP90) (Koncz & Schell, 1986). Agrobacterium cells harboring constructs encoding the nYFP and cYFP fusion proteins were resuspended in a freshly prepared infiltration medium containing 10 mM MgCl, 10 mM MES-KOH (pH 5.7), and 200 μM acetosyringone. These bacterial suspensions were co-infiltrated into *N. benthamiana* leaves at a 1:1 ratio. Empty vectors (nYFP and cYFP) were included as negative controls. After 48-h incubation at 22°C, YFP fluorescence was detected using a confocal laser-scanning microscope (TCS SP5, Leica) with excitation at 514 nm and emission at 527 nm.

### Split luciferase assay

The full-length coding sequences MaHDZIV6, MaJAZ5 and MaJAZ8 were cloned into the pCAMBIA1300-nLUC vector, while the entry clones of MaTFR, were inserted into the pCAMBIA1300-cLUC vector (Chen *et al*., 2008). Agrobacterium strain GV3101, carrying the respective nLUC and cLUC constructs, was individually resuspended in freshly prepared infiltration medium containing 10 mM MgCl, 10 mM MES-KOH (pH 5.7), and 150 μMacetosyringone. The bacterial suspensions were co-infiltrated into *N. benthamiana* leaves in a 1:1 ratio. After infiltration, the leaves were treated with 100 μM D-luciferin, and luciferase (LUC) activity was quantified 48 hours later using a low-light charge-coupled device (CCD) imaging system (Bio-Rad).

### Dual-luciferase reporter assay

To evaluate the transactivation potential of the transcriptional regulators *MaTFR* and *MaHDZIV6*, their coding sequences (CDSs) were cloned into the destination vector pSITE-3CA (Chakrabarty *et al*., 2007) generating effector constructs driven by the 2xCaMV 35S promoter. Reporter constructs (*proMaHDZIV6, proMaFLS1, proMaFLS2, proMaF3’5’H1, proMaF3’5’H2)* in the p635nRRF vector contained each promoter fragment upstream of the *LUC* gene (encoding firefly luciferase), while REN (Renilla luciferase) expression was driven by the constitutive *CaMV35S* promoter, serving as an internal control for dual-luciferase reporter assays, the effector and reporter constructs were transformed into *A. tumefaciens* strain GV3101 (pMP90) (GoldBio). The constructs were then infiltrated into the abaxial side of *N. benthamiana* leaves using a syringe, following the protocol outlined by Walter *et al*. (2004). After 48 h, luminescence was measured in extracts from the infiltrated leaf discs using the Dual-Luciferase Reporter Assay System (Promega), as per the manufacturer’s instructions. LUC (firefly luciferase) and REN (Renilla luciferase) activities were quantified with a POLARstar Omega multimode plate reader (BMG Labtech), and LUC activity was normalized to REN activity. The results are presented as the mean ± SD of the LUC/REN ratio from four independent biological replicates.

### Yeast oneDhybrid (Y1H) assay

The Y1H assay was performed using the Y1H gold yeast strain (Takara Bio Inc., Shiga, Japan). The promoter fragments of *MaFLS1, MaFLS2, MaF3’5’H1, MaF3’5’H2, and MaHDZIV6* were cloned into the pABAi vector (Cat. No. 630491, Takara Bio Inc.). The pABAi vector harbouring promoter fragments was linearized with Bsp119I (BstBI) enzyme and transformed into Y1H Gold strain according to the Matchmaker™ manual. The MaTFR and MaHDZIV6 CDS fragments were also cloned into the pGADT7-gateway vector. The construct plasmids having the desired genes and empty pGADT7-gateway vector as control were transformed into pre-transformed Y1H strains containing target promoter sequence. Interactions were assessed using Aureobasidin A (AbA)-supplemented media. To determine the minimum inhibitory concentration (MIC) values, the growth of various promoter-integrated Y1H strains was evaluated on SD/-Ura media containing different concentrations of AbA. For the MBS-containing promoter fragments, the MIC values were 1 μM for *MaFLS1,* 900 nM for *MaF3’5’H1,* and 850 nM for *MaF3’5’H2*. In the case of L1-box-containing promoter fragments, the MIC was 900 nM for *MaFLS2*, and 100 nM for *MaF3’5’H1*.

### ChIP-qPCR assay

ChIP assays were performed following the protocol as described by (Saleh *et al*., 2008) with minor modifications. GFP-tagged MaTFR, MaHDZIV6 fusion proteins vacuum infiltered into banana ECS. 4 days post-infiltration, 3-4 g of tissue per sample was cross-linked with 1% (v/v) formaldehyde to stabilize protein–DNA complexes. The samples were flash-frozen in liquid nitrogen, ground, and homogenized in nuclei isolation and lysis buffers before sonication. Chromatin was sonicated using a Diagenode Bioruptor Plus at 4°C. Sonicated chromatin samples were pre-cleared with Protein A agarose beads (Millipore #16-157), and chromatin immunoprecipitation was performed using 2 µL of anti-GFP antibody (Abcam, Catalog #ab290) for specific binding in two biological replicates. As a negative control, 2 µL of anti-IgG antibody (Abcam, Catalog #ab48386) was used. RT-qPCR analysis was carried out using 5 µL of 2x SYBR Green PCR Master Mix (Applied Biosystems), 1 µL of ChIP DNA, and 0.5 µL each of forward and reverse primers. The reactions underwent 40 amplification cycles as previously described. The relative fold enrichment of target promoter fragments was determined using the 2^(-ΔCT) method (Livak & Schmittgen, 2001). Primer sequences used for ChIP-qPCR are provided in the Table S1.

### Transient overexpression and gene silencing in embryogenic cells of banana

For transient overexpression studies, CDSs of *MaTFR*, *MaHDZIV6*, and *MaJAZs* were cloned into the Gateway-compatible destination vector pANIC6b (Mann *et al*., 2012). This vector enables strong expression of transgenes in monocot species via *A. tumefaciens*-mediated transformation (Rajput *et al*., 2022; Naik *et al*., 2025). pANIC6b carries a hygromycin resistance gene driven by the *O. sativa* actin1 promoter (*proOsACT1, LOC4338914*), a *Panicum virgatum* ubiquitin1+3 promoter (proPvUBI) driving a GUSPlus reporter, and a Gateway cassette under the control of the *Zea mays* ubiquitin1 promoter and intron for efficient overexpression. The overexpression constructs, along with an EV control, were introduced into *A. tumefaciens* strain EHA105 (Mehra *et al*., 2019) and used to transform banana embryogenic cells by vacuum infiltration, following a previously established protocol (Kaur *et al*., 2020). After five days of co-cultivation on modified medium (MS basal salts, HiMedia), successful transformation was confirmed via GUS staining. For gene silencing experiments, 250–350 bp CDS fragments of *MaTFR* and *MaHDZIV6* were cloned into the RNA interference vector pANIC8b, which is also Gateway-compatible and designed for gene silencing in monocots. These constructs were similarly transformed into *A. tumefaciens* strain EHA105 and introduced into banana embryogenic cells using the same method as for overexpression. GUS staining was used to verify transformation. The tissues were then harvested and frozen in liquid nitrogen for subsequent expression analysis (RT-qPCR) and targeted metabolite profiling (LC-MS).

### Statistical test

Statistical analyses were conducted using one-way and two-way analysis of variance (ANOVA), followed by Tukey’s multiple comparison test. For specific pairwise comparisons, unpaired Student’s t-tests were applied. Statistical significance is indicated as follows: P ≤ 0.05 (*), P ≤ 0.01 (**), P ≤ 0.001 (***), P ≤ 0.0001 (****); ns denotes non-significant differences.

## Results

### Structural diversity and histochemical assays of banana trichomes

In banana, the Plantain (PLT) cultivar (AAB or ABB) is trichome-free (TF), while Grand Naine (GN) and Red Banana (RB) cultivars (AAA genome) are trichome-rich (TR). Trichomes are generally found on all aerial parts of most plant species. Interestingly, trichomes are exclusively found on the inflorescence stalk of the GN and RB cultivars (Fig. 1a). Other tissues like pseudo stem, leaves, bract, flower, and fruit tissues, are devoid of trichomes in all the banana cultivars. Stereomicroscopy and Scanning Electron Microscopy (SEM) imaging showed three types of semi-glandular trichomes in banana: (1) unicellular, unbranched type (2) multicellular, unbranched type, and (3) multicellular, branched type trichomes (Fig. 1b-d). Propidium iodide staining confirmed that banana trichomes are both unicellular and multicellular type, with later being composed of two to eight cells (Fig. 1d). Histochemical staining was used to study the presence of types of metabolites in the trichomes of in both GN and RB. The DPBA staining showed intense green fluorescence signals signifying the presence of flavonols, specifically kaempferol derivatives, with varied accumulation patterns found across trichomes. A few trichomes also accumulate flavonols in the glandular head of trichomes. The presence of a bluish color following NADI staining in the trichomes represents the accumulation of terpenoids in overall trichomes as well as in a few trichome glands. Sudan IV staining, giving a reddish hue in whole trichomes, suggests the lipid content (Fig. 1e).

**Fig. 1.**
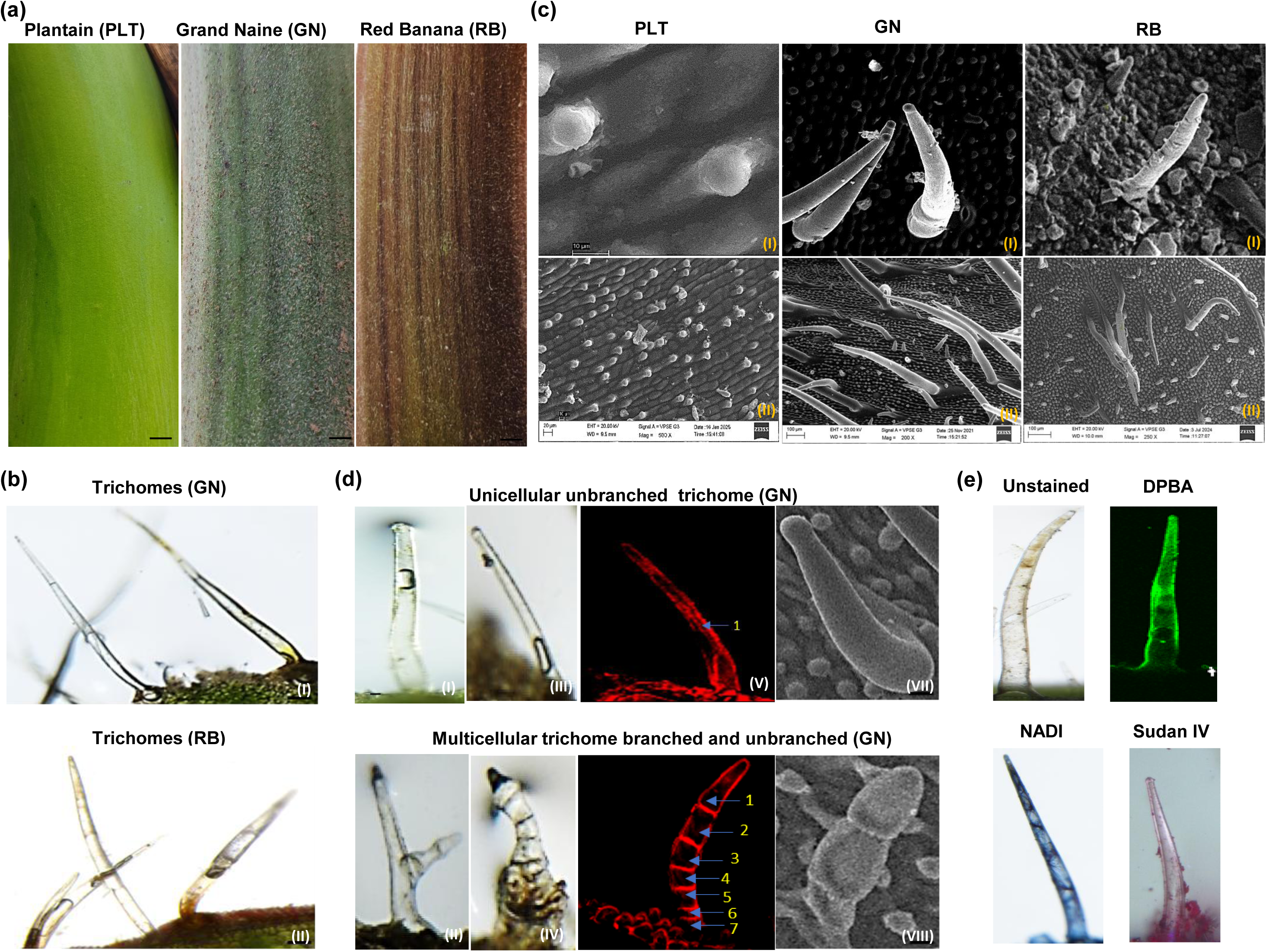
Morphological features of trichomes in *Musa* species. **(a)** A pictorial representation comparing the epidermis of TF (PLT) with the TR (GN and RB) cultivars. **(b)** Stereomicroscopy images showing the diversity of trichomes in GN and RB. **(c)** Scanning electron microscopy (SEM) analysis of the epidermis reveals only initiating cells, with no evidence of trichome morphogenesis in the inflorescence stalk of PLT. In contrast, well-developed trichomes are observed on the inflorescence stalks of GN and RB. **(d)** Banana trichomes are predominantly of the unicellular, unbranched type (I, III), although a few branched, multicellular trichomes are also observed (II, IV). Propidium iodide staining (V, VI) and SEM (VII, VIII) confirm the presence of both unicellular and multicellular trichomes in banana. In multicellular trichomes, the number of cells vary among different trichome types. **(e)** Unstained and stained trichomes: DPBA staining produces intense green fluorescence, indicating the presence of flavonols. The bluish color, due to NADI reagent, highlights terpenoid accumulation in trichomes and some trichome glands. Additionally, Sudan IV staining imparts a reddish hue to the trichomes, marking the presence of lipids.

### Transcriptome analysis of TF and TR banana cultivars

In order to gain insights into the differentially expressed genes (DEGs) between the TF and TR epidermis, transcriptome sequencing was carried out. Transcriptomic profiling across the three cultivars revealed significant transcriptional variations between TF (PLT) and TR epidermis (GN and RB) as visualized by volcano plots (Fig. S1a). The MapMan analysis revealed distinct patterns of secondary metabolism gene expression among PLT, GN and RB. Notably, genes associated with flavonoid biosynthesis were majorly expressed in RB (Fig. S1b). The comparison of up-regulated and down-regulated DEGs across the PLT, GN, and RB cultivars revealed both unique and shared expression patterns (Fig. 2a-b; Dataset S1). A total of 688 (up-regulated) and 570 (down-regulated) DEGs were identified in GN as compared to PLT; total 3906 (up-regulated) and 3624 (down-regulated) DEGs in RB as compared to PLT; and total of 3216 (up-regulated) and 1920 (down-regulated) DEGs in RB as compare to GN at adjusted p-value < 0.05 and fold change ≥ 2 (Fig. 2a-b; Dataset S1). GO enrichment analysis was performed to identify the significant biological processes, pathways, and functions associated with these DEGs. Interestingly, the L-phenylalanine biosynthetic process and UDP-glycosyltransferase activity were enriched terms in RB compared to PLT and GN, while the flavonoid biosynthetic process was more enriched in PLT than in GN. The sesquiterpene biosynthesis process was enriched in GN cultivar. Response to hormone, specifically response to jasmonic acid (JA), was majorly enriched in RB cultivar (Fig. 2c-h, Dataset S2).

**Fig. 2.**
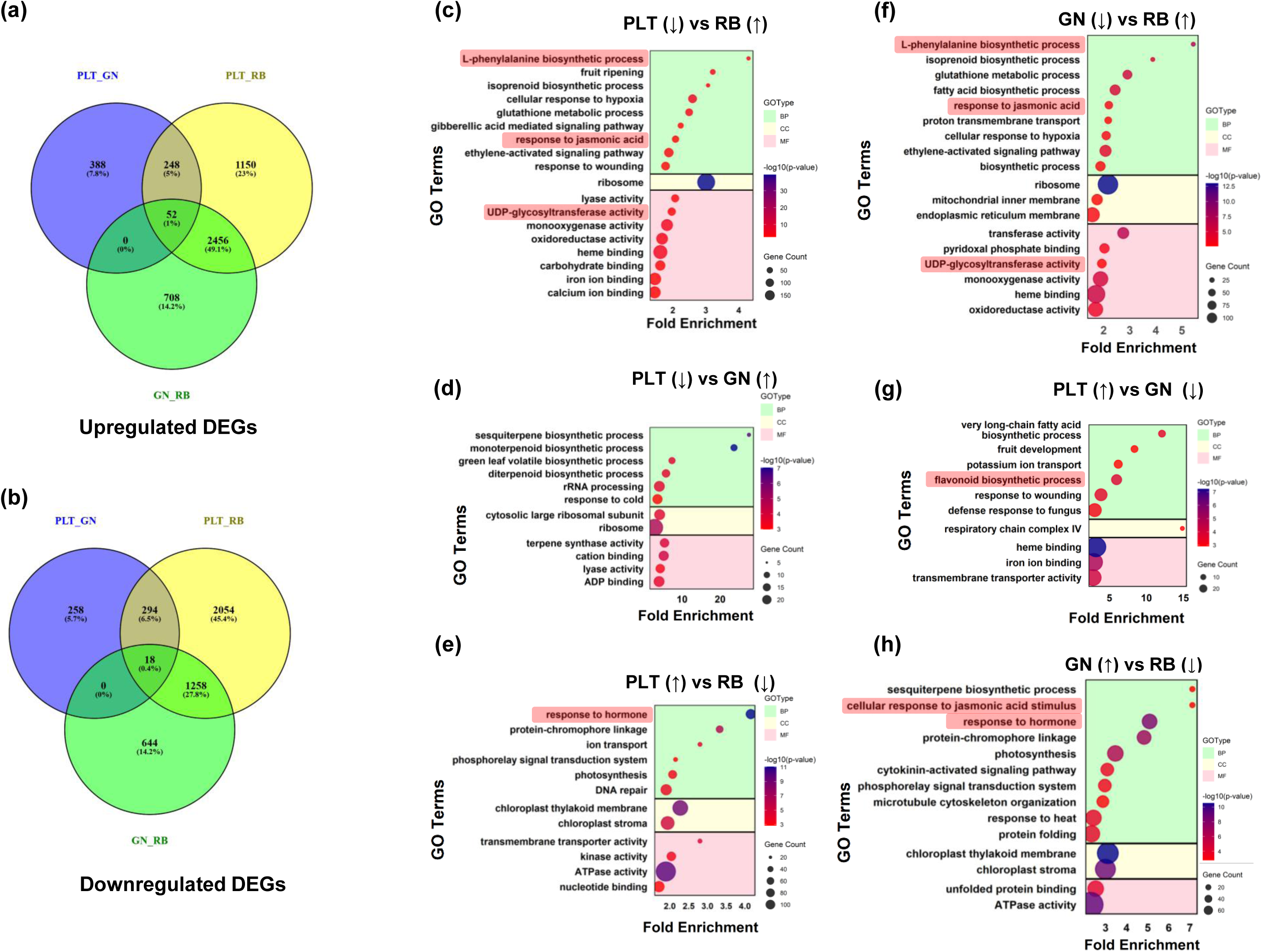
DEGs count & GO enrichment analysis of TF and TR tissues of banana. **(a)** Venn diagram showing overlapping and unique upregulated and **(b)** downregulated differentially expressed genes (DEGs) in PLT (TF) vs GN (TR), PLT vs RB (TR) and GN vs RB comparisons. Bubble plots showing the GO enrichment analysis of DEGs in different comparisons: **(c)** PLT (↓) vs RB (↑), **(d)** PLT (↓) vs GN (↑), **(e)** PLT (↑) vs RB (↓), **(f)** GN (↓) vs RB (↑), **(g)** PLT (↑) vs GN (↓), **(h)** GN (↑) vs RB (↓). The up (↑) and down (↓) arrow represents upregulated and downregulated gene expression, respectively. The x-axis and y-axis represent the fold enrichment and different GO terms, respectively. The GO terms for biological process (BP), cellular component (CC) and molecular function (MF) are highlighted with different colours. Bubble size corresponds to the gene count and colour bar scale represents −log10(p-value).

We explored the expression of defense hormone-related genes including abscisic acid (ABA), ethylene (ET), salicylic acid (SA) and JA in epidermis of TF (PLT) and TR (GN and RB) cultivars. Differential expression of various hormone-related genes across these cultivars highlights their potential roles in regulating defense responses and trichome development (Fig. S2a-d). In order to identify potential transcription factors concerning the regulation of trichome morphogenesis, the expression of genes belonging to different transcription factors families was analyzed. To this end, bHLH, WRKY, and C2H2 TF encoding genes with average FPKM ≥10 in PLT, GN and RB were selected for expression profiling (Fig. S3). The MYB and HDZIP sub family IV TFs have been reported to play pivotal role in the development of trichomes (Sen & Das, 2016; Sharif *et al*., 2021; Xie *et al*., 2021). Therefore, the trichome-related MaMYB TFs were identified from our previous study (Pucker et., 2020) and their expression were analyzed in PLT, GN and RB (Fig. S3). The expression of *Macma4_10_g17890* was significantly higher in TR cultivars (GN and RB). Further, we have analyzed the expression of previously reported 21 HDZIP sub family IV TF genes in banana (Pandey *et al*., 2016). The expression of certain TFs identified by homology with functionally known trichome-related TFs in Arabidopsis was analyzed in these tissues. The expression of *Macma4_09_g01400* was found to be higher in both GN and RB as compared to PLT (Fig. S3).

### Flavonoid biosynthesis genes expression and metabolite content are differentially modulated in TF and TR banana cultivars

A comprehensive pictorial representation of the flavonoid biosynthesis pathway illustrating the sequential enzymatic steps involved in the production of various flavonoid classes is presented. We used transcriptome and qRT-PCR data to analyze the expression patterns of flavonoid biosynthetic genes. Most of the structural genes of the flavonoid pathway displayed differential expression among the three samples. Out of the six homologs of *MaCHS* genes*, MaCHS1* was not expressed in any of the cultivar. The expression of most of the *MaCHS* genes (*MaCHS2, MaCHS3, MaCHS4 MaCHS5* and *MaCHS6*) was higher in TR cultivars as compared to TF cultivar. In contrast, *MaCHS6* expression was highest in GN. Overall, the expression of *MaCHS* paralogs was higher in the TR epidermis of both GN and RB cultivars, except *MaCHS4*, which was highly expressed in the TF cultivar of banana (Fig. S4a, b). The expressions of *MaCHI2* and *MaF3H1-2* were differentially varied in GN cultivar (Fig. S4b). The expression of *MaF3’H* and *MaFLS* paralogs was consistently high in TR cultivars compared to TF cultivar. Similarly, *MaDFR1* and *MaANS* expression levels were elevated in TR cultivars, while the opposite trend was observed for *MaDFR2* (Fig. S4a, b). The differential expression patterns of structural genes of flavonoid biosynthesis suggest the modulated accumulation of flavonoids in different cultivars.

We carried out targeted analysis of flavonoids and other phenolics in the TF and TR banana cultivars using LC-MS to study the accumulation of specialized metabolites in trichomes. Our analysis suggested that phenylpropanoids such as *p*-coumaric acid, cinnamic acid, and syringic acid were more abundant in TR as compared to TF cultivar. However, the contents of caffeic and chlorogenic acids were found to be higher in the TF compared to TR cultivars.

Among flavonoids, the content of eriodyctiol, was higher in the TR as compared to TF cultivar. The contents of other flavonoids (flavonols and dihydro-flavonol) viz. kaempferol, myricetin, dihydro-myricetin, rutin hydrate, and kaempferol derivatives were higher TR in comparison to TF cultivar. However, the dihydro-quercetin was present at higher level in TF compared to TR cultivars (Fig. S5a). In general, the content of proanthocyanidins was reported to be higher in the TR as compared to TF cultivar (Fig. S5b). Total anthocyanin content was significantly higher in the RB in comparison to PLT and GN cultivars (Fig. S5c). Thus, the metabolite analysis suggests contrasting variations in the TF and TR cultivars. The differential accumulation of these metabolites might be helpful as protective molecules against biotic and abiotic stresses.

### Identification and characterization of a trichome-related R2R3 MYB TFs

Certain R2-R3 MYB transcription factors plays a crucial role in trichome development across various plant species (Su *et al*., 2020; Li *et al*., 2022a). Therefore, we performed the phylogenetic analysis of putative trichome-related R2-R3 MYB TFs reported in our earlier study in banana (Pucker *et al*., 2020) along with functionally characterized trichome-related MYB transcription factors in other species (Fig. 3a). Further, the qRT-PCR analysis confirmed that six trichome-related R2R3-MYB genes (*Macma4_06_g16640.1*, *Macma4_02_g22860.1*, *Macma4_09_g11600.1*, *Macma4_05_g06590.1*, *Macma4_09_g27080.1*, and *Macma4_10_g17890.1*) exhibited significantly higher expression in TR (GN and RB) cultivars compared to the TF cultivar (Fig. 3b, S6). In contrast, two genes (*Macma4_06_g40680.1* and *Macma4_07_g20460.1*) displayed higher expression in the TF cultivar in comparison to TR cultivars (Fig. S6). Based on phylogenetic clustering of Macma4_10_g17890 with AtMYB5 from Arabidopsis and VvMYBA1 from *Vitis vinifera* and its higher expression in TR cultivars, we selected it as a potential trichome and flavonol regulator R2-R3 MYB (MaTFR). The *MaTFR* emerged as a strong candidate for further functional characterization to elucidate its role in trichome development in banana.

**Fig. 3.**
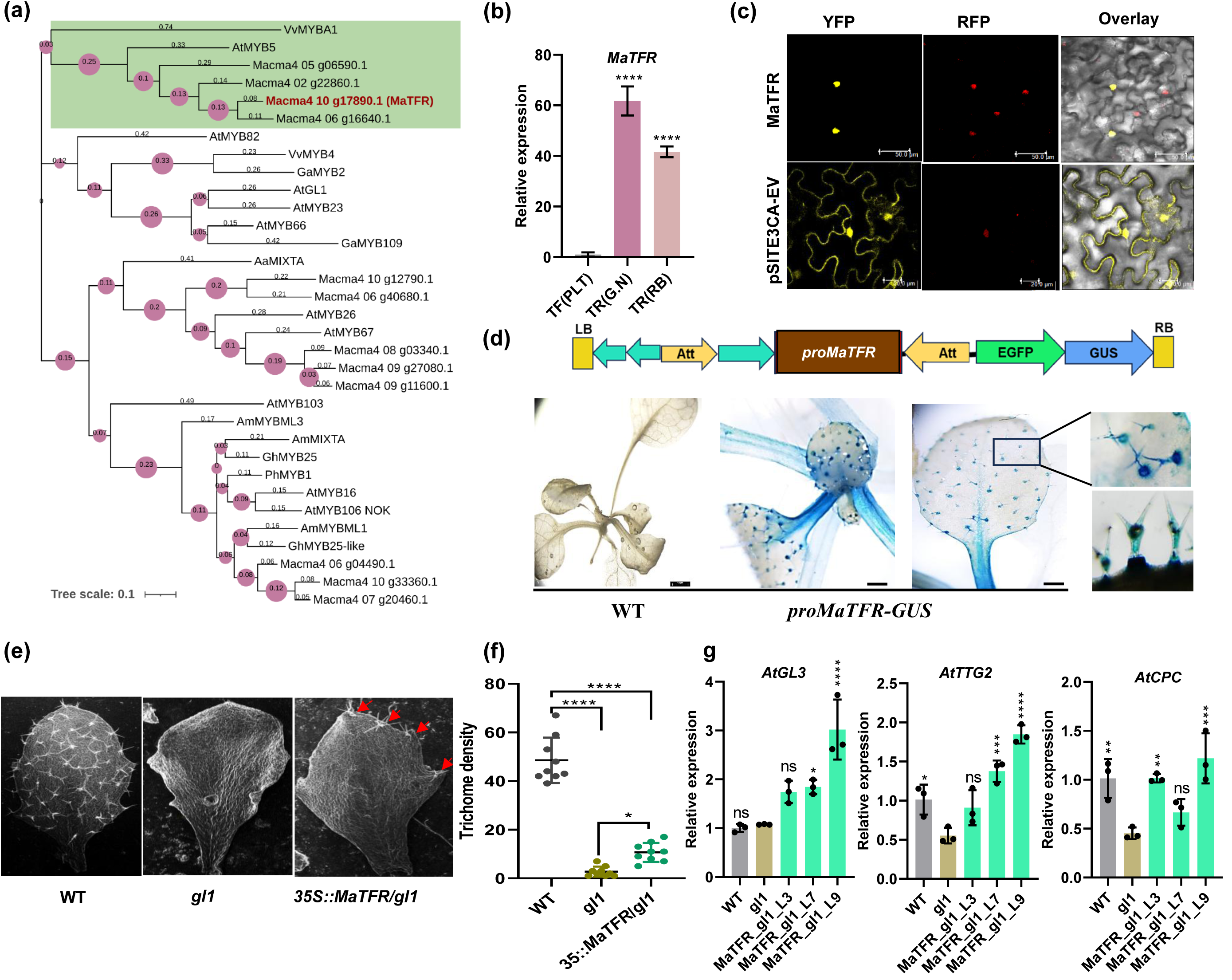
*MaTFR* is highly expressed in trichomes and partially restores function in Arabidopsis. **(a)** Phylogenetic tree of trichome-related MYBs from banana and other plant species. MYBs are grouped into distinct clades, with the clade containing MaTFR highlighted in green. Branch lengths are indicated along each branch, and bootstrap support values are visualized by the size of pink bubbles at each node. **(b)** *MaTFR* expression high in TR (GN and RB) as compared to TF(PLT) cultivar of banana as determined by RT-qPCR. Values shown are means ± SD of three biological replicates. **(c)** Subcellular localization of MaTFR in *N. benthamiana* leaves. The YFP-MaTFR fusion protein localizes to the nucleus and coincides with the nuclear localization signal red fluorescent protein (NLS-RFP) as observed in confocal microscopy. EV pSITE-3CA as used as negative control to visualize free YFP. Scale bar, 50µm. **(d)** Representative image of the GUS-GFP cassette used for promoter-GUS lines driven by 2 kb *MaTFR* promoter. Wildtype seedling used as a control. T3 homozygous *MaTFR* promoter-GUS lines predominantly show GUS on the trichomes. **(e)** For the complementation assay, the WT leaf used as a control, shows uniform trichomes, *gl1* leaf is trichome free, however the *MaTFR* complemented lines leaf shows trichomes on the leaf margins. **(f)** Similar area used for calculation for trichome density on Arabidopsis leaf for WT, *gl1,* and complemented lines in (n=9). **(g)** RT-qPCR analysis demonstrated that trichome-specific regulatory genes (*AtGL3*, *AtTTG2*, and *AtCPC*) were significantly upregulated in *MaTFR*-complemented lines within the *gl1* mutant background. Asterisks represent the level of significant change at *P* ≤ 0.05 (*)*, P* ≤ *0.01* (**), *P* ≤ *0.001* (***), P ≤ 0.0001 (****); ns = not significant.

We determined the subcellular localization of MaTFR by transiently co-infiltrating constructs encoding YFP fusion proteins into *Nicotiana benthamiana* epidermal cells, along with the nuclear marker NLS-RFP (red fluorescent protein fused to a nuclear localization signal). Fluorescence from the YFP fusions was observed exclusively in the nucleus, precisely co-localizing with NLS-RFP confirming that MaTFR are nuclear-localized proteins (Fig. 3c).

A promoter-reporter construct containing the 2 kb upstream regulatory region of the *MaTFR* gene fused to the GUS reporter was used to develop transgenic *Arabidopsis thaliana* lines. GUS staining of these lines demonstrated that *MaTFR* promoter activity was predominantly confined to trichomes (Fig. 3b). Notably, the highest intensity of GUS staining was observed in the initial cells during the early stages of trichome development in true leaves of 10-day-old Arabidopsis seedlings under normal growth conditions, suggesting that the *MaTFR* promoter is most active during these initial phases (Fig. 3d). Although the staining was most dominant in the developing trichome cells, the entire trichome structure exhibited GUS activity, indicating sustained expression throughout trichome maturation.

To verify the role of *MaTFR* in trichome formation, a genetic complementation assay was conducted using the *Arabidopsis gl1 mutant* (Bloomer *et al*., 2012). Unlike wild type (WT), the *gl1 mutant* shows defective trichome development on its leaves and stems. However, expression of *35Spro::MaTFR* transgene in the *gl1 mutant* partially restored trichome formation in leaves (Fig. 3e-f). Gene expression analysis in *35Spro::MaTFR* transgenic lines showed that MaTFR activates downstream transcription factor genes, including *AtGL3*, *AtTTG2*, and *AtCPC* (Fig. 3g), which are the key regulators of trichome formation in Arabidopsis (Johnson *et al*., 2002). Additionally, MaTFR enhanced the expression of *AtF3’H* and *AtFLS1*, which are involved in flavonoid biosynthesis. Consistently, flavonol quantification in *35Spro::MaTFR*-transgenic lines revealed elevated levels of myricetin and quercetin, (Fig. S7a-b), suggesting that MaTFR positively regulates both flavonol biosynthesis and trichome development.

### Transient overexpression and RNAi silencing of *MaTFR* in banana embryogenic cells (ECS) substantially modulate the expression of structural genes of flavonol biosynthesis and *MaHDZIV6*

Since MaTFR display preferential expression in TR samples, which also accumulate higher amounts of flavonoids, we studied whether MaTFR is also involved in the regulation of flavonoid biosynthesis. To this end, MaTFR was transiently overexpressed and silenced in banana ECS. The GUS staining confirmed the successful transformation of the expression cassette in ECS (Fig. 4a). The qRT-PCR analysis revealed that *MaTFR* expression was significantly up regulated (23-fold, P ≤ 0.0001) in *proZmUBI::MaTFR-OE* ECS, while its expression was notably reduced (0.53-fold, P = 0.0013) in *MaTFR*-silenced ECS compared to their respective empty vector (EV) transformed ECS. Flavonol biosynthesis genes showed differential expression patterns in response to *MaTFR* regulation. *MaFLS1* was up regulated (2.2-fold) in OE ECS but downregulated (0.46-fold) in *MaTFR-*silenced ECS, whereas *MaFLS2* expression remained largely unchanged in both conditions. Genes encoding enzymes involved in flavonoid biosynthesis-related CYP450 proteins also exhibited varying expression patterns. In *MaTFR*-OE ECS, *MaF3’5’H1* (1.8-fold), *MaF3’5’H2* (1.9-fold), *MaF3’5’H4,* and *MaF3’5’H6* were up regulated, while *MaF3’5’H3, MaF3’5’H5,* and *MaF3’5’H7* showed no significant change. Conversely, in *MaTFR-*silenced ECS, *MaF3’5’H1* (0.3-fold), *MaF3’5’H2* (0.59-fold), and *MaF3’5’H7* were downregulated, while *MaF3’5’H3, MaF3’5’H4, MaF3’5’H5,* and *MaF3’5’H6* remained unchanged (Fig. 4b).

**Fig. 4.**
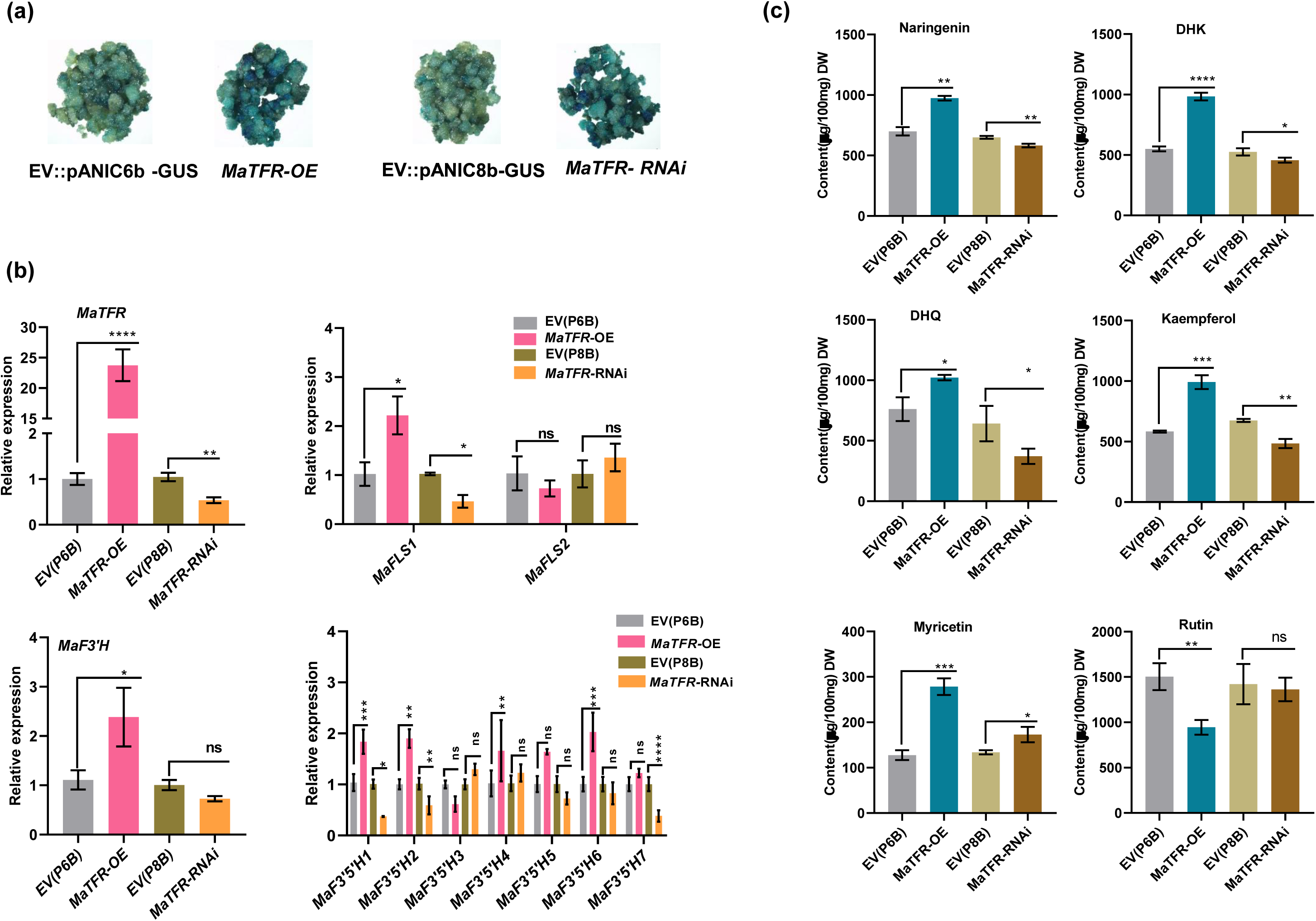
Transient overexpression and silencing of *MaTFR* and its target gene expression in banana ECS. **(a)** Representative images of GUS-stained banana ECS showing successful transient transformation of OE and silencing constructs of *MaTFR* and their respective EV constructs. Images were captured four days after transformation. **(b)** Relative expression levels of *MaTFR*, *MaFLS1/2*, and *MaF3’5’H1-7* were analysed in banana ECS subjected to transient *MaTFR* overexpression and silencing using RT-qPCR. Expression was compared to EV controls: pANIC6b for OE and pANIC8B for silencing. Values are shown as fold changes from three biological and three technical replicates. **(c)** The accumulation of kaempferol, and myricetin aglycones and glycan forms (DHK, DHQ and rutin) was measured in *MaTFR* OE and silenced banana ECS. Flavonol levels are represent in µg per 100 g of dry weight (DW) from three replicates. Statistical analysis was performed using one-way ANOVA followed by Tukey’s multiple comparison test. Significance levels: *P* ≤ 0.05 (*)*, P* ≤ *0.01* (**), *P* ≤ *0.001* (***), P ≤ 0.0001 (****); ns = not significant.

Flavonol quantification further supported the regulatory role of MaTFR. In *MaTFR*-OE ECS, levels of naringenin, kaempferol, myricetin, and dihydro-flavonols (DHK, DHQ) were increased, while in MaTFR silenced ECS, naringenin, kaempferol, DHK, and DHQ levels were reduced. Myricetin levels showed a slight increase in silenced ECS, though significantly lower compared to *MaTFR*-OE. Notably, rutin content decreased in *MaTFR*-OE ECS but increased in MaTFR-silenced ECS (Fig. 4c).

We also identified candidate trichome-related HDZIP sub family IV members in banana through phylogenetic analysis (Fig. S8a). MaHDZIV6 (Macma4_04_g21150.1), MaHDZIV8 (Macma4_05_g14400.1) and MaHDZIV16 (Macma4_10_g21800.1) were grouped with trichome-development related HDZIPs from monocots plant species. Among these, only *MaHDZIV6* was found to express at significantly higher level in the TR (GN and RB) cultivars, as compared to TF cultivars (Fig. S8b). Interestingly, in *MaTFR*-OE ECS, the expression of *MaHDZIV6* and *MaHDZIV8* was significantly upregulated (6-fold) and 1.9-fold, respectively, while *MaHDZIV16* expression remained unchanged. In contrast, in *MaTFR*-silenced ECS, *MaHDZIV6* and *MaHDZIV8* showed no significant changes in expression, but *MaHDZIV16* was notably upregulated by about 1.7-fold. (Fig. S8c). These findings indicate that MaTFR acts as a preferential upstream regulator of MaHDZIV6, contributing to trichome development either directly or indirectly, possibly through the modulation of *MaHDZIV6* expression.

### MaTFR is a transcriptional activator of structural genes of flavonol biosynthesis and *MaHDZIV6*

Overexpression of *MaTFR* in banana ECS led to a significant upregulation of structural genes involved in flavonol biosynthesis, as well as increased expression of *MaHDZIV6*. In contrast, silencing of *MaTFR* resulted in reduced expression of these genes, suggesting that they may function as downstream targets of the *MaTFR* transcription factor. Structural analysis of a 2.5 kb *proMaFLS1*, 1.3 kb *proMaF3’5’H1*, 1kb *proMaF3’5’H2* and 2 kb *proMaHDZIV6* promoter fragment revealed several MYB core elements (Fig. 5a). To assess the transactivation potential of MaTFR on*, MaFLS1, MaF3’5’H1, MaF3’5’H2,* and *MaHDZIV6* promoters, we conducted dual-luciferase assays in *N. benthamiana* leaves. The analysis suggested positive transactivation activity of *MaFLS1, MaF3’5’H1*, *MaF3’5’H2,* and *MaHDZIV6* promoter fragments by MaTFR. (Fig. 5b-c). Further, we performed Y1H assays to validate the binding of MaTFR to *proMaFLS1-989, MaF3’5’H1-574, MaF3’5’H2-414,* and *proMaHDZIV6-893,* promoters. For this, we utilized a promoter fragment containing the MBS motif to assess its interaction with the MaTFR-AD fusion protein. The Y1H growth assay results demonstrated a strong binding affinity between MaTFR to *proMaFLS1, proMaF3’5’H1, proMaF3’5’H2,* and *proMaHDZIV6* fragments, confirming its potential role in regulating flavonol biosynthesis in banana (Fig. 5d). Further to test the direct binding of MaTFR via the MBS, we overexpressed pro*ZmUBI*::*MaTFR* construct into banana ECS, followed by a ChIP assay using an anti-GFP antibody. The ChIP-qPCR assay showed specific binding of MaTFR to MBS-containing promoter regions of target genes. In the *MaFLS1* promoter, the F2 fragment (215 bp) showed (6.2-fold) enrichment, while the F1 fragment (166 bp) exhibited no significant enrichment. For the *MaF3’5’H1* promoter, the F1 fragment (183 bp) demonstrated (3.2-fold) enrichment. In the *MaF3’5’H2* promoter, the F2 (129 bp) fragment showed (2.4-fold) enrichment, whereas the F1 (133 bp) fragment did not. Additionally, in the *MaHDZIV6* promoter, both F1 (151 bp) and F2 (138) fragments 2.9-fold and 3.9-fold enrichment respectively, while the F3 (147 bp) fragment showed no significant enrichment. These findings indicate that MaTFR directly regulates the expression of *MaFLS1, MaF3’5’H1, MaF3’5’H2*, and *MaHDZIV6* by binding to their respective MBS elements (Fig. 5e).

**Fig. 5.**
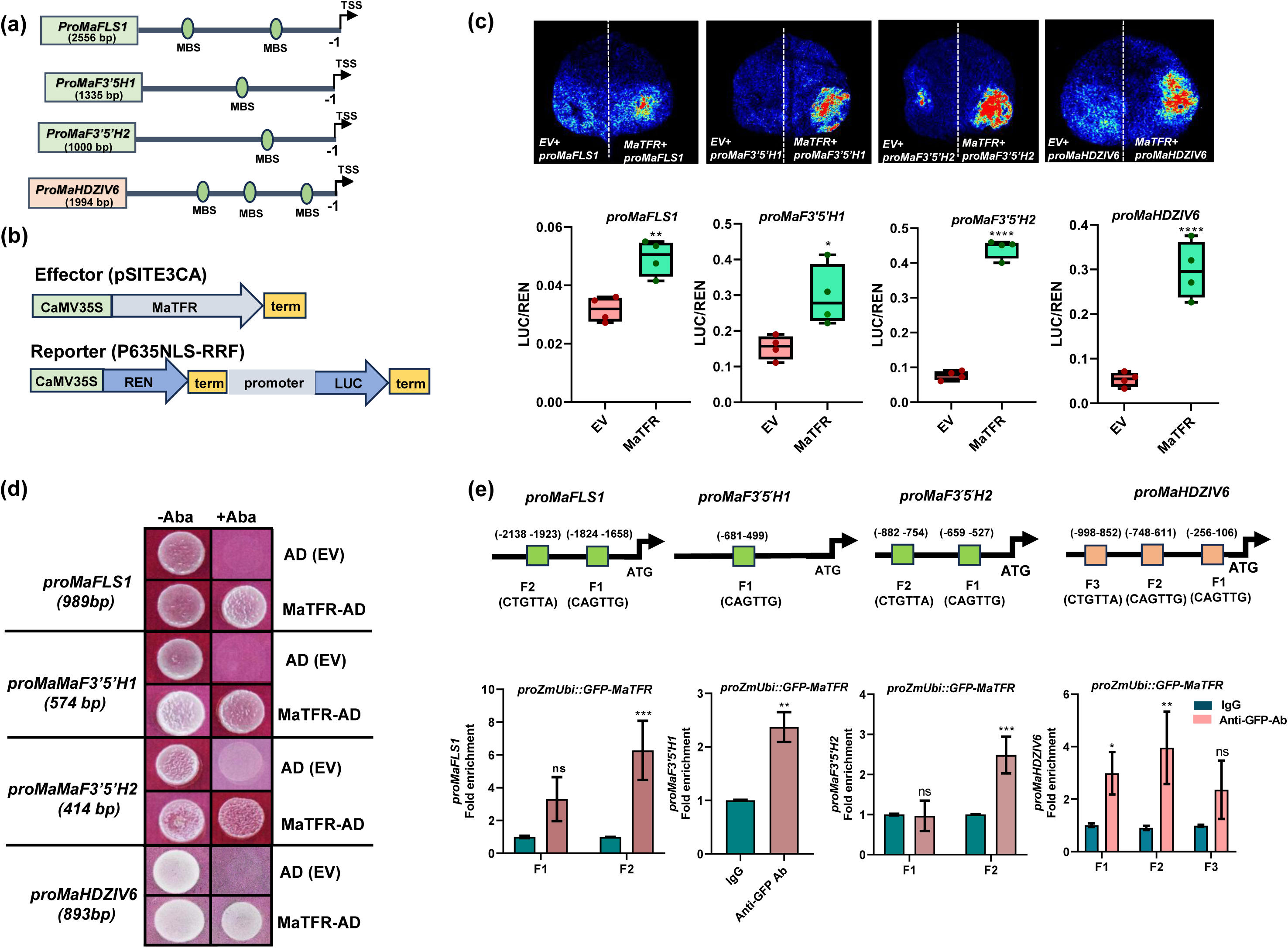
Dual luciferase assay and binding analysis of MaTFR to *MaFLS1, MaF3’5’H1, MaF3’5’H2 and MaHDZIV6* promoters. **(a)** Schematic representation of the promoters used in dual-luciferase assays. **(b)** A schematic diagram of the effector and reporter constructs used in dual-luciferase assays. **(c)** The transactivation activity of *MaFLS1, MaF3’5’H1, MaF3’5’H2* and *MaHDZIV6* promoters by MaTFR TF was analysed using a dual luciferase assay. Results are presented as the mean ± SD of LUC/REN ratios from four independent biological replicates (*n* = 4). **(d)** Yeast One-Hybrid (Y1H) assay confirming the interaction of *MaTFR* with *proMaFLS1, proMaF3’5’H1, proMaF3’5’H2* and *proMaHDZIV6* containing MBS motifs. The empty *pGADT7* vector (EV) was used as a negative control. Interaction was indicated by yeast growth on synthetic dropout media lacking uracil (Ura–) and supplemented with aureobasidin A (AbA). AD = activation domain. **(e)** ChIP-qPCR was performed to examine the binding of MaTFR to various promoter fragments. In *proMaFLS1*, enrichment was observed only in F1 fragment, with no enrichment in F2 fragment. In *proMaF3’5’H1*, enrichment was detected in F1 fragment. For *proMaF3’5’H2*, enrichment occurred only in F2 fragment, with no enrichment in F1. In the case of *proMaHDZIV6*, MaTFR was enriched in both F1 and F2 fragments, while no enrichment was observed in F3 fragment. Each fragments contain the MBS consensus sequence. Fold enrichment values are represented as means ± SD from two biological replicates. Ab (antibody).

### Functional characterization of trichome-related MaHDZIV6 transcription factor

We observed that the *MaHDZIV6* expression is significantly higher in TR cultivars compared to the TF epidermis, suggesting its potential role in cultivar-specific traits (Fig. 6a). Further we determined the subcellular localization of MaHDZIV6 by expressing it as a YFP fusion protein along nuclear NLS-RFP marker in *N. benthamiana* epidermal cells. YFP fluorescence was observed in the nucleus, overlapping with NLS-RFP marker indicating, that MaHDZIV6 is a nuclear-localized protein (Fig. 6b). A promoter-reporter construct comprising the 1994 bp upstream regulatory region of the *MaHDZIV6* gene fused to the GUS reporter was used to generate transgenic *A. thaliana* lines. GUS staining revealed that the *MaHDZIV6* promoter was predominantly active in trichomes of true leaves of 10-day old seedlings. (Fig. 6c). The highest *GUS* expression was observed in early trichome precursor cells, indicating peak promoter activity during the initial stages of trichome development, with less activity in mature trichomes.

**Fig. 6.**
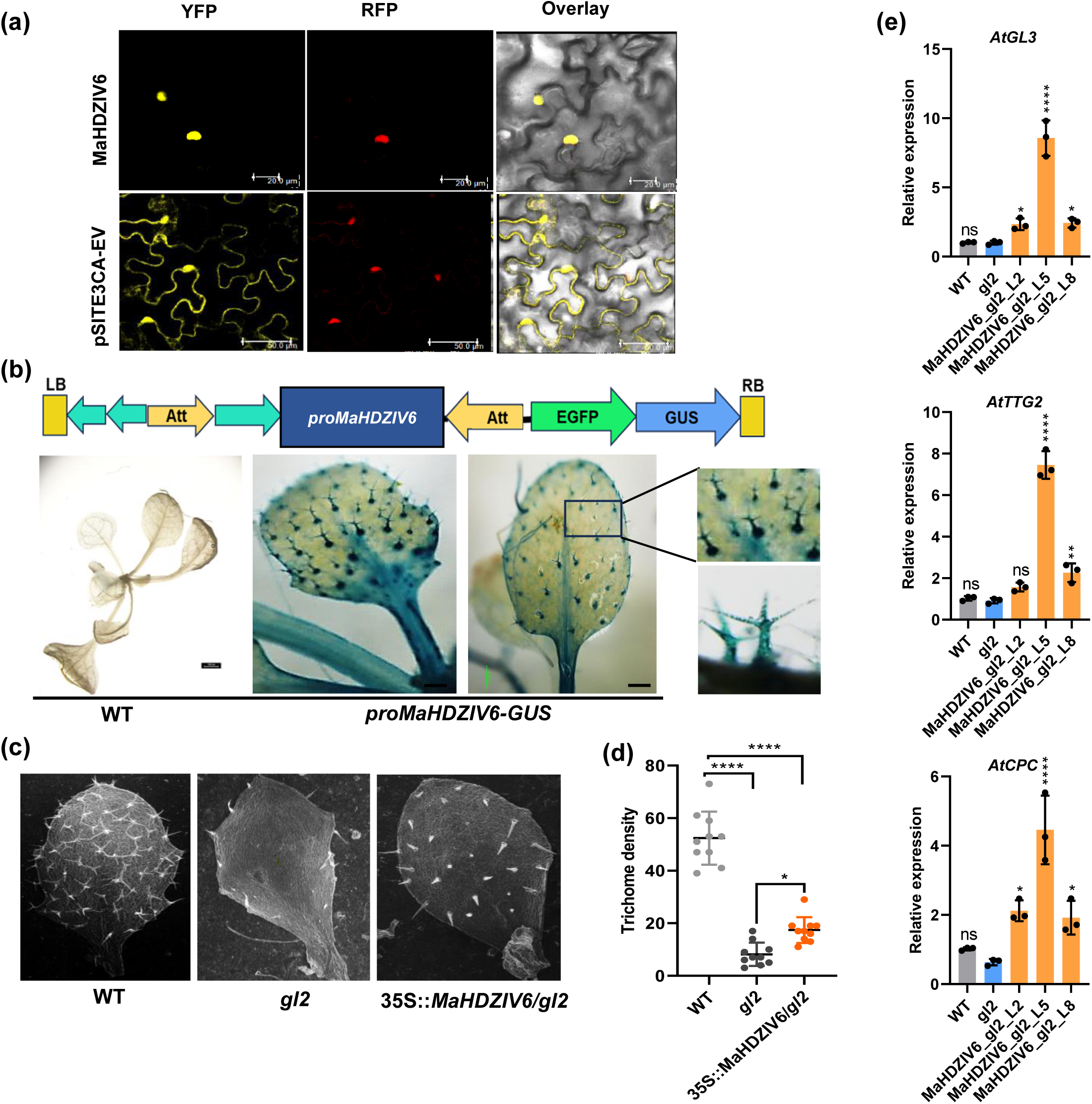
MaHDZIV6 is predominantly expressed in trichomes and partially complements *gl2* Arabidopsis mutant. **(a)** Subcellular localization of MaHDZIV6 was examined in *N. benthamiana* leaves. The YFP-MaHDZIV6 fusion protein was observed in the nucleus, where it co-localized with the nuclear localization signal red fluorescent protein (NLS-RFP) in confocal microscopy. An EV (pSITE-3CA) was used as a negative control to visualize free YFP. Scale bar: 50 μm. **(b)** A schematic representation of the GUS-GFP cassette used in promoter-GUS lines driven by the *MaHDZIV6* promoter. Col-0 seedlings served as a control. Positive T3 *MaHDZIV6* promoter-GUS lines predominantly exhibited GUS activity in trichomes. **(c)** The leaves of wild type displayed uniform trichome distribution, while *gl2* mutant leaves were devoid of trichomes. In contrast, MaHDZIV6-complemented lines exhibited trichomes along the leaf margins. **(d)** Trichome density was quantified from equal leaf areas in WT, *gl2,* and complemented lines (n = 6), indicating partial restoration of trichome formation by MaHDZIV6. **(e)** The RT-qPCR analysis revealed modulation of trichome-specific regulatory genes (*AtGL3*, *AtTTG2*, and *AtCPC*) in *MaHDZIV6*-complemented lines within the *gl2* mutant background.

The genetic complementation assay was performed in the *Arabidopsis gl2* mutant to verify the role of *MaHDZIV6* in trichome formation. The *gl2* mutant, unlike the WT, displays unbranched trichomes only on the margins of its leaves (Khosla *et al*., 2014). However, *35Spro::MaHDZIV6* transgenic expression in the *gl2* mutant partially restored trichome formation on leaves of complemented lines. However, only unbranched trichome phenotype was observed on entire leaves (Fig. 6d-e). Gene expression analysis revealed that MaHDZIV6 can activate downstream TF genes, including *AtGL3*, *AtTTG2*, and *AtCPC* (Fig. 6f), which are well-known regulators of trichome formation. Interestingly, MaHDZIV6 also upregulated the expression of *AtF3’H* and *AtFLS1*, which are involved in the flavonol biosynthesis pathway in heterologous system. Moreover, flavonol quantification in *MaHDZIV6*-complemented lines showed increased levels of myricetin, kaempferol, and quercetin, indicating that MaHDZIV6 positively regulates flavonol biosynthesis alongside trichome development (Supplemental Fig. 9a-b).

### MaTFR physically interacts with MaHDZIV6 protein

We demonstrated that MaTFR and MaHDZIV6 TFs may be potential regulators of semi-glandular trichome development in banana. We carried out the co-expression analysis of these transcription factors and flavonoid biosynthesis genes with other differentially expressed transcription factors to gain a broader view of putative regulatory partners. We found that *MaHDZIV6* displayed co-expression with certain transcription factor family members such as MYB, bHLH, bZIP, JAZ, etc. (Fig. 7a). We further confirmed the direct interaction between MaHDZIV6 and MaTFR proteins by the yeast two hybrid (Y2H) assay. The strong physical interaction between MaTFR and the MaHDZIV6 proteins was demonstrated by the appearance of a blue color on a QDO selective medium supplemented with X-α-Gal and 3-AT. (Fig. 7b). To further validate these findings *in planta*, we conducted bimolecular fluorescence complementation (BiFC) and luciferase complementation imaging (LCI) assays in *N. benthamiana* leaves. In the BiFC assays, YFP reconstitution—indicative of protein-protein interaction was detected when constructs encoding MaTFR-YFP^N^ and MaHDZIV6-YFP^C^ were co-infiltrated into *N. benthamiana* leaves (Fig. 7c). Similarly, LCI confirmed the interaction between MaTFR MaHDZIV6 proteins (Fig. 7d). We also tested the localization of both the interacting proteins into *N. benthamiana* leaves, and observed that both proteins colocalized in the nucleus, indicating their interaction (Fig. 7e). These findings strongly suggest that MaTFR and MaHDZIV6 proteins work together to facilitate trichome development in banana.

**Fig. 7.**
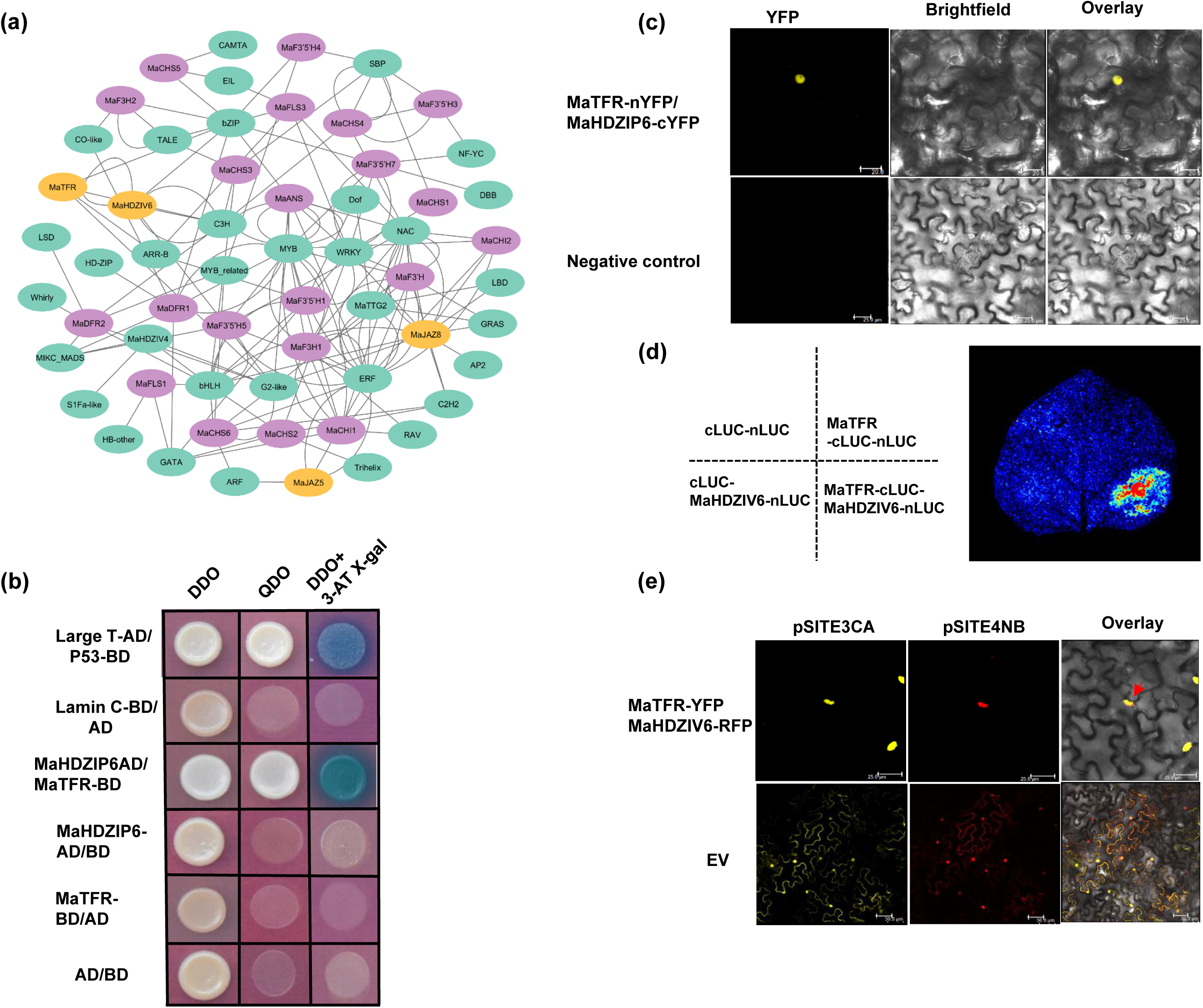
Interaction between MaTFR and MaHDZIV6 proteins. **(a)** Co-expression network of selected genes involved in trichome development and flavonoids biosynthesis. **(b)** The interaction between MaTFR and MaHDZIV6 was tested using a yeast two-hybrid assay. MaTFR was fused to the GAL4 activation domain (AD), while MaHDZIV6 was fused to the GAL4 DNA-binding domain (BD). Yeast cells were grown on double dropout (DDO) medium (-Leu - Trp) for selection and quadruple dropout (QDO) medium (-Leu -Trp -Ade -His) to confirm the interaction. When plated on QDO medium supplemented with X-α-Gal and aureobasidin A (QDO/X/A), blue colonies formed, indicating a strong interaction through the cleavage of X-α-Gal by α-glucosidase. **(c)** BiFC assay was performed to confirm the interaction of MaTFR and MaHDZIV6 in living plant cells (*in planta*). The fluorescence signal was detected under a microscope, demonstrating protein interaction. YFPN + YFPC constructs were used as a negative control. Scale bar: 20 μm. **(d)** Luciferase complementation imaging (LCI) assay showing the interaction between The MaTFR and MaHDZIV6. Three negative control combinations were tested: MaTFR-nLUC + cLUC, nLUC + MaHDZIV6-cLUC, and nLUC + cLUC. Luminescence signals confirmed that MaTFR and MaHDZIV6 interact in plant cells**. (e)** The co-localization assay reveals that MaTFR-pSITE3CA-YFP and MaHDZIV6-pSITE4NB-RFP are localized in the nucleus, where they overlap, indicating their interaction.

### Transient overexpression and RNAi silencing of *MaHDZIV6* modulate structural genes of flavonol biosynthesis in banana ECS

Given that MaHDZIV6 is predominantly expressed in TR (GN and RB) tissues with high flavonoid accumulation, we aimed to uncover its role in regulating flavonoid biosynthesis. To explore this, *MaHDZIV6* was transiently overexpressed and silenced in banana ECS. Successful transient expression was confirmed through GUS staining of the ECS (Fig. 8a). qRT-PCR analysis confirmed that *MaHDZIV6* expression was significantly upregulated (19.7-fold, P ≤ 0.0001) in *proZmUBI::MaHDZIV6* transformed ECS, while its expression was notably reduced (0.28-fold, P ≤ 0.0005 in *MaHDZIV6* silenced ECS compared to their respective EV controls. Flavonol biosynthesis genes exhibited differential expression patterns in response to the genetic manipulation of *MaHDZIV6* regulation. *MaFLS1* was upregulated (2.5-fold) in *proZmUBI::MaHDZIV6* OE ECS but downregulated (0.77-fold) in *MaHDZIV6* silenced ECS. Similarly, *MaFLS2* was upregulated (2.28-fold) in OE ECS and downregulated (0.79-fold) in silenced ECS. The flavonoid biosynthesis-related CYP450 genes also showed varying expression patterns. In *MaHDZIV6* OE ECS, *MaF3’5’H1* (5.3-fold), *MaF3’5’H2* (3.59-fold), *MaF3’5’H3,* and *MaF3’5’H5* were up regulated, while *MaF3’5’H6* showed no significant change. In contrast, in *MaHDZIV6* silenced ECS, *MaF3’5’H1* (0.31-fold) and *MaF3’5’H2* (0.338-fold) were down regulated, whereas *MaF3’5’H3, MaF3’5’H4, MaF3’5’H5, MaF3’5’H6,* and *MaF3’5’H7* remained unchanged (Fig. 8b). Flavonol quantification in *MaHDZIV6* OEECS showed increased levels of kaempferol, myricetin, and dihydro-flavonols (DHK and DHM), while naringenin levels remained unchanged. Conversely, in *MaHDZIV6*-silenced ECS, naringenin, kaempferol, myricetin, and dihydro-flavonols (DHK and DHQ) were significantly reduced as compared to EV control. Notably, rutin levels decreased in *MaHDZIV6* OE ECS but increased in MaHDZIV6 silenced ECS (Fig. 8c). These findings highlight MaHDZIV6 as a crucial regulator of flavonol biosynthesis, shaping the expression of *MaFLS, MaF3’H,* and *MaF3’5’H* paralogs and playing a significant role in the metabolic pathways of banana.

**Fig. 8.**
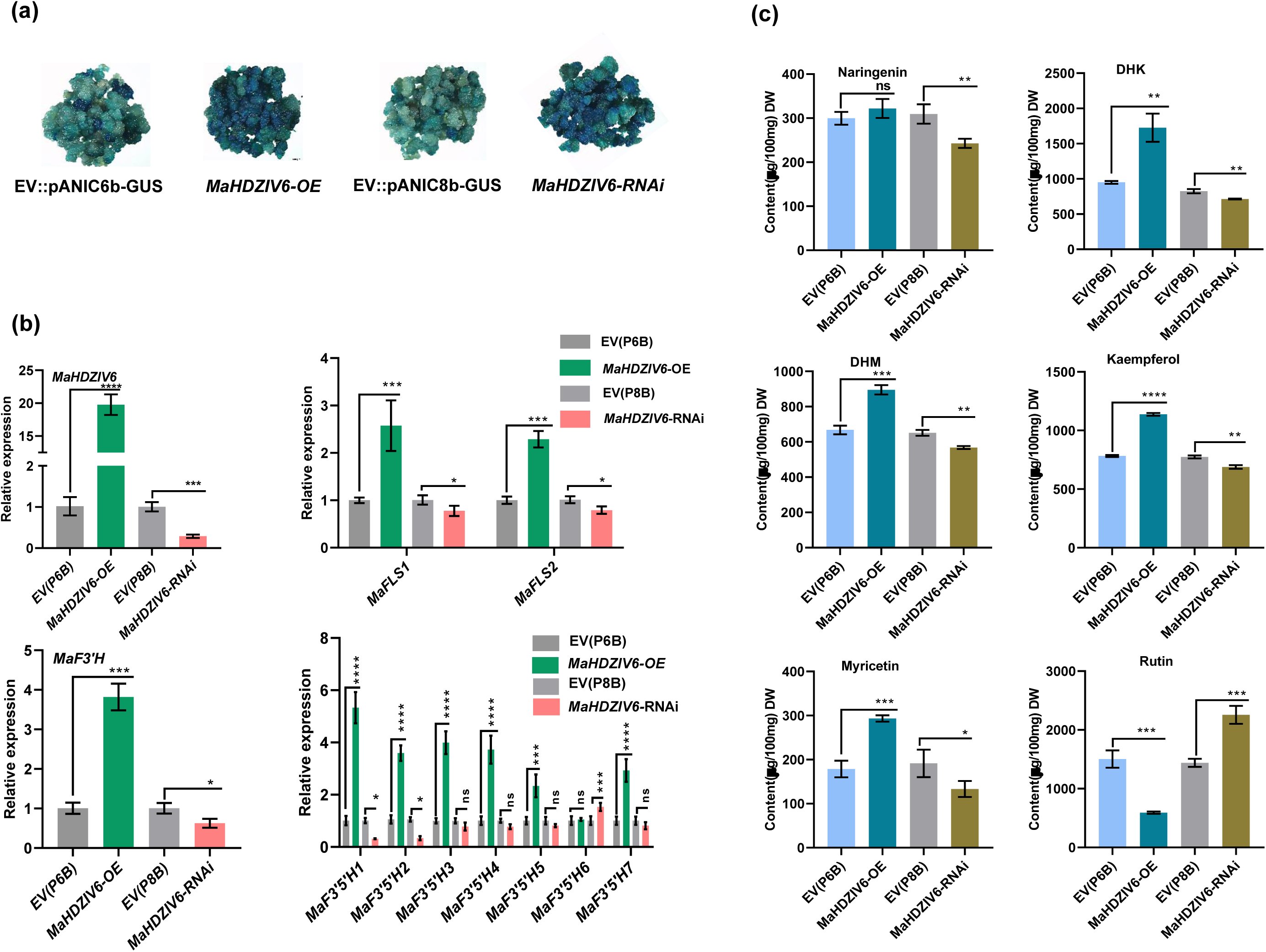
Transient overexpression (OE) and silencing of *MaHDZIV6* and its target gene expression in banana ECS. **(a)** Representative images of GUS-stained banana ECS confirm successful transient transformation with *MaHDZIV6* OE and silencing constructs. Images were taken four days post-transformation. **(b)** The relative expression of *MaHDZIV6*, *MaFLS1/2*, *MaF3’H*, and *MaF3’5’H* paralogs was analysed in banana ECS subjected to transient *MaHDZIV6* OE and silencing using RT-qPCR. Expression levels were compared to EV controls-pANIC6b for OE and pANIC8B for silencing. Data are presented as three biological and three technical replicates. **(c)** The accumulation of kaempferol, quercetin, and myricetin aglycones was quantified in *MaHDZIV6* OE and silenced banana ECS. Flavonol contents are expressed in µg per 100 g of dry weight (DW) based on three replicates. Statistical analysis was conducted using one-way ANOVA followed by Tukey’s multiple comparison test. Asterisks *, **, ***, and *** represents level of significance at p-value ≤0.05, ≤0.01, ≤ *0.001*, and ≤0.0001, respectively, ns = not significant.

### MaHDZIV6 is a transcriptional activator of the structural genes of flavonol biosynthesis

The expression of structural genes involved in flavonol biosynthesis was found to be up regulated in *MaHDZIV6* OE banana ECS and downregulated in *MaHDZIV6* silenced ECS, compared to the EV control. These data suggest that MaHDZIV6, apart from regulating trichome development, may act as a transcriptional regulator of flavonoid biosynthesis. Structural analysis of *proMaFLS1–2556*, and *proMaFLS2–1500*, *MaF3’5’H1-1335*, and *MaF3’5’H2-1000,* fragments revealed single HDZIP binding sites L1-box (TAAATG) present in all four promoter fragments (Fig. 9a). In order to study the potential transcriptional regulation of flavonoid biosynthesis offered by MaHDZIV6, dual luciferase-based transactivation assay was carried in *N. benthamiana* leaves. Results demonstrated that *MaFLS1, MaFLS2, MaF3’5’H1,* and *MaF3’5’H2* promoter fragments showed positive transactivation activity, with the highest activation seen for the *MaF3’5’H1* and *MaF3’5’H2* promoter (Fig. 9b-c). These results suggest that MaHDZIV6 may bind to the *MaFLS,* and *MaF3*′*5*′*H* promoter and modulates the flavonol content. Further we performed Y1H assay to validate the interaction between MaHDZIV6 with the *MaFLS2 and MaF3’5’H1* promoters. To this end, we utilized promoter fragments of same size containing the L1-box to assess their interaction with the MaHDZIV6-AD fusion protein. The Y1H growth assay demonstrated strong binding affinity between MaHDZIV6 and the *MaFLS2,* and *MaF3’5’H1* promoter fragments, suggesting a direct regulatory role of MaHDZIV6 in modulating *MaFLS2* and *MaF3’5’H1* transcription (Fig. 9d). To further test the direct binding of MaHDZIV6 via the L1-box, we overexpressed pro*ZmUBI::MaHDZIV6* construct into banana ECS and performed a ChIP assay using an anti-GFP antibody. The ChIP-qPCR assay revealed enrichment of the L1-box containing promoter fragments of pro*MaFLS2* (140 bp) and pro*MaF3’5’H1* (189 bp). However, no fold enrichment occurs in the case of pro*MaFLS1*, and pro*MaF3’5’H2* fragments (Fig. 9e).

**Fig. 9.**
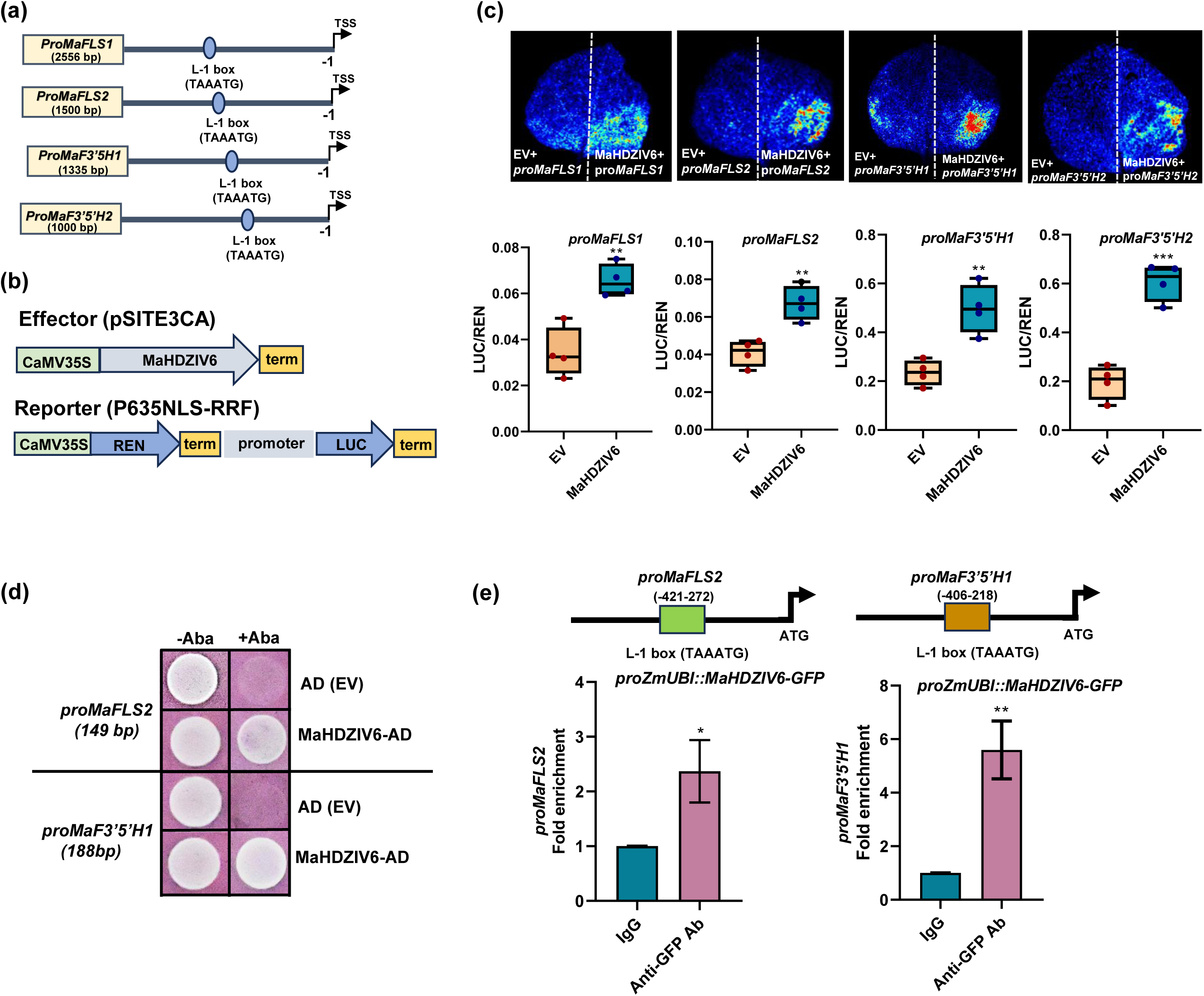
Dual luciferase assay and binding analysis of MaHDZIV6 to the promoters of *MaFLS1, MaFLS2, MaF3’5’H1,* and *MaF3’5’H2*. **(a)** Schematic diagram of the promoters showing the L-1 box. **(b)** Schematic representation of the effector and reporter constructs used in the dual-luciferase assays is shown. **(c)** The transactivation of the *proMaFLS1–2556, proMaFLS2-1500, proMaF3’5’H1-1335 and proMaF3’5’H2-1000* promoter by MaHDZIV6 was analysed using the dual-luciferase assay. The results are shown as the mean ± SD of LUC/REN ratios from four independent biological replicates (*n* = 4). **(d)** The Y1H assay was conducted to confirm the interaction between MaHDZIV6 and the L1-box-containing promoter fragments of *proMaFLS2* and *proMaF3’5’H1*. An empty pGADT7 vector (EV) was used as a negative control. Successful interaction was indicated by yeast growth on synthetic dropout media lacking uracil (Ura–) and supplemented with aureobasidin A (AbA). AD = activation domain. **(e)** ChIP-qPCR was conducted to evaluate the binding of MaHDZIV6 to the promoters of *MaFLS2* and *MaF3’5’H1*, both containing the L1 consensus sequence (L1-box) motif. Fold enrichment values are presented as the mean ± SD from two biological replicates. Ab (antibody).

### Defense hormones are differentially accumulated in the PLT, RB and GN epidermis

Defense hormones play crucial roles in the regulation of trichome development and metabolism (Li *et al*., 2022c). Therefore, in order to study their potential role in the trichome development, we conducted targeted defense hormones profiling using LC-MS to quantitatively analyze defense hormones, including Salicylic acid (SA), Abscisic acid (ABA), and Jasmonic acid (JA) and their precursors (*cis*-OPDA) and derivatives (JA-Ile). The results revealed that SA predominantly accumulated in PLT and GN with the lowest levels detected in RB. Conversely, ABA content was highest in RB and lowest in the PLT cultivars. Interestingly, the precursor of JA, cis-OPDA, was higher in PLT cultivars. However, the active forms of JA were significantly higher in TR cultivars, GN and RB. JA levels were notably elevated in RB, while no significant changes were observed in PLT and GN (Fig. S10a-b). The differential accumulation of defense hormones suggests their critical involvement in developmental processes and metabolic pathway modulation. The higher JA content in RB and GN may play a role in trichome genesis in these cultivars.

### Methyl JA induces the expression of *MaTFR* and *MaHDZIV6* and flavonoids biosynthetic genes in banana seedlings

Transcriptome analysis indicated that among all phytohormones, JA responses were predominately modified in banana cultivars displaying differential development of trichomes. Further, the phytochemical analysis also suggested that active JA pool is comparatively higher in the TR (GN and RB) cultivars. In view of these findings, we hypothesized that JA might regulate the expression of identified regulators (MaTFR and MaHDZIV6) of trichome development and flavonoid biosynthesis in banana. To confirm this, we examined the effects of exogenous methyl JA application on key TFs and structural genes of flavonoid biosynthesis pathway. Methyl JA treatment significantly upregulated *MaTFR* expression at multiple time points (6, 12, 24, and 48 h), with peak expression observed at 12 h. Similarly, *MaHDZIV6* expression was consistently elevated at the same time points but peaked earlier, at 6 h. However, both TFs showed a decline in expression by 72 h (Fig. S11a-b). Flavonoid biosynthesis genes exhibited dynamic expression patterns in response to methyl JA. *MaCHS* and its paralogs expression peaked at 6 h, with sustained upregulation at 12, 24, and 48 h, but showed negligible changes at 72 h. *MaCHI2* expression was notably elevated at 6 h, with no significant changes at later time points. Similarly, *MaFLS1* and *MaFLS2* peaked at 6 h, but while *MaFLS1* expression remained stable, *MaFLS2* showed additional increases at 12 and 24 h. Additionally, M*aF3’5’H* and its paralogs were upregulated in early time points but declined at later stages (Fig. S12a). Flavonol quantification further supported these findings. Kaempferol levels increased following Methyl JA treatment, reaching their highest levels at 72 h, while myricetin peaked at 6 h, aligning with the expression *of MaFLS1, MaFLS2*, and other biosynthetic genes. Quercetin and rutin levels were also increased significantly post-treatment (Fig. S12b). Overall, these results strongly suggest that methyl JA treatment activates *MaTFR* and *MaHDZIV6*, leading to the induction of flavonoid biosynthesis genes. This regulation contributes to trichome development and enhanced stress tolerance, highlighting JA’s critical role in modulating plant defense and metabolic pathways.

### MaJAZ5 and MaJAZ8 interacts with MaTFR and attenuating its transcriptional activity

The phylogenetic analysis of identified MaJAZ proteins in banana showed clustering into three major clades along with previously known JAZ proteins from other plant species (Fig. S13a). We also performed the expression profiling of all *MaJAZ* genes, where certain MaJAZs were highly expressed in TR cultivars. We selected five *MaJAZ5* (Macma4_03_g06070), *MaJAZ8* (Macma4_05_g05150), *MaJAZ9* (Macma4_05_g05960), *MaJAZ20* (Macma4_09_g14550), and *MaJAZ21* (Macma4_09_g15840) based on their distinct expression patterns and their close clustering with previously identified JAZ having role in trichome development (Fig. S13b-c). These MaJAZ proteins contained TIFY domain (ZIM domain) and Jas motif (C-terminal domain). In Y2H screening, all five MaJAZ proteins were tested for interaction with the MaTFR proteins. These connections are mediated by the Jas domain of the JAZ proteins and the C-terminal domain of MaTFR protein. The results showed that only MaJAZ5 and MaJAZ8 interacted with MaTFR protein (Fig. 10a). However, none of the MaJAZs interact with MaHDZIV6 (Fig. S14a). Further, both MaJAZ5 and MaJAZ8 showed nuclear localization as observed by overlapping YFP and nuclear RFP marker signals in the nucleus (Fig. S14b). The interactions of MaJAZ5 and MaJAZ8 with MaTFR were validated *in planta* by BiFC and LCI assays were conducted in *N. benthamiana* leaves (Fig. 10b-c). To investigate the role of MaJAZ5 and MaJAZ8 after interacting with MaTFR, we analyzed the transcriptional activity of MaTFR and found significant reduction in transactivation of *proMaFLS1*, *proMaF3’5’H1, proMaF3’5’H2* and *proMaHDZIV6*, thereby reducing trichome development and flavonol biosynthesis (Fig. 10d).

**Fig. 10.**
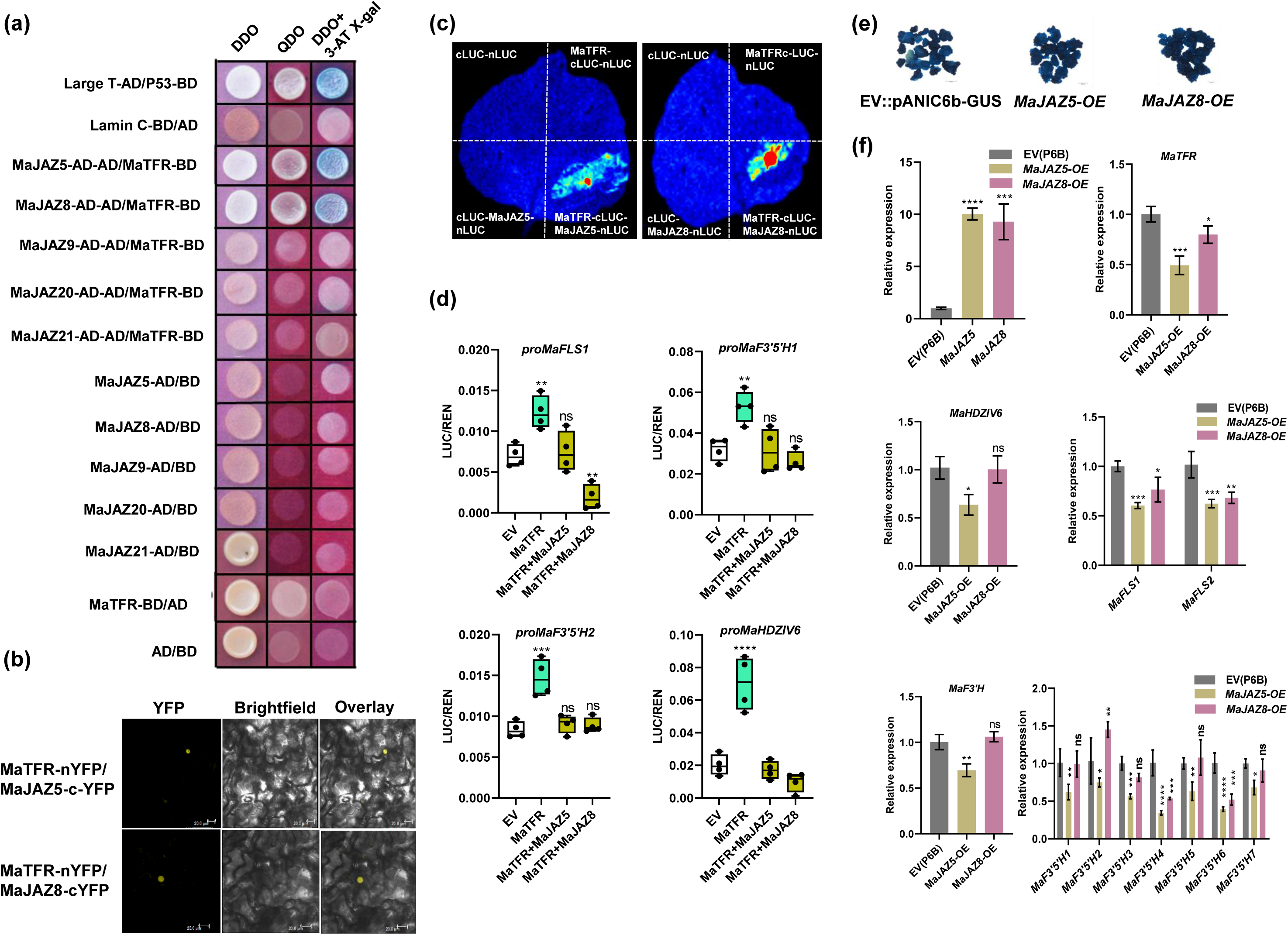
MaJAZ physically interacts with MaTFR and diminishes its transcriptional activity. **(a)** Yeast-two-hybrid assays showing the interaction of MaTFR with MaJAZ5 and MaJAZ8. AD, GAL4 activation domain; BD, GAL4 DNA-binding domain; DDO, double dropout synthetic defined (SD)−Leu−Trp medium; QDO, quadruple dropout SD–Leu–Trp–Ade–His medium; QDO/X/A, QDO + X-α-Gal) medium. Blue colonies show cleavage of X-α-Gal by α-glucosidase to generate blue colour products, suggesting a strong interaction. **(b)** Bimolecular fluorescence complementation (BiFC) assay indicating the *in-planta* interaction of MaTFR with MaJAZ5 and MaJAZ8. *YFP^N^* + *YFP^C^* constructs were used as a negative control. Bar, 20 μm. **(c)** Luciferase complementation imaging (LCI) assay showing the interaction between MaTFR and MaJAZ5 and MaJAZ8. The following three pairs of constructs were used as negative controls: *MaTFR-nLUC + cLUC*, *nLUC + MaJAZ-cLUC* and *nLUC + cLUC*. **(d)** MaJAZ5 and MaJAZ8 interacts with MaTFR and represses the *MaHDZIV6*, *MaFLS1*, *MaF3’5’H1* and *MaF3’5’H2* promoters. Results from transient luciferase assays in *N. benthamiana* leaves. Constructs harbouring the firefly luciferase reporter gene (*LUC*) are driven by 1994 bp *proMaHDZIV6,* 1500 bp *proMaFLS2*, 1335 bp *proMaF3’5’H1* and 1000 *proMaF3’5’H2* promoter fragments and were transiently co-infiltrated with *MaTFR*, *MaJAZ5* and *MaJAZ8* effector constructs, either alone or in the indicated combinations. *, *P* ≤ 0.05; **, *P* ≤ 0.01; ***, *P* ≤ 0.001, as determined by one-way analysis of variance. Data are shown as the means ± SD of four biological replicates**. (e)** Representative images of GUS-stained banana ECS confirm the successful transient transformation with MaJAZ5 and MaJAZ8, captured four days post-transformation. **(f)** The relative expression levels of MaJAZ5/8, *MaTFR*, *MaHDZIV6, MaFLS1/2*, *MaF3’H MaF3’5’H1-7* were analysed in banana ECS subjected to transient *MaJAZ5* and *MaJAZ8* overexpression (OE) by RT-qPCR. Expression data was compared to EV controls-pANIC6b for OE. Values are shown as fold changes from three biological replicates and three technical replicates.

### Transient overexpression of *MaJAZ5 and MaJAZ8* in banana ECS reduces the expression of *MaTFR, MaFLS* and *MaF3’5’H*

To further analyse the regulatory role of MaJAZ5 and MaJAZ8, we transiently overexpressed *proZmUBI::MaJAZ5/8* in banana ECS and confirmed by GUS staining (Fig. 10e). The qRT-PCR analysis confirmed that the expression of *MaJAZ5* and *MaJAZ8* was enhanced up to 9-fold and 10-fold, respectively, in *proZmUBI::MaJAZ5/8* banana ECS than the EV control. Furthermore, the expression of the target gene *MaTFR* was significantly downregulated in both *MaJAZ5* and *MaJAZ8* OE tissues. This data strongly suggests that MaJAZ5 and MaJAZ8 show strong repressing activity over *MaTFR*. We earlier found that MaTFR is an upstream regulator of *MaHDZIV6*. As expected, the expression of *MaHDZIV6* was also reduced in the case of *MaJAZ5* OE, however, its expression was not significantly modulated in *MaJAZ8* OE ECS. The expression of *MaFLS1* was reduced in *MaJAZ5* OE ECS, while *MaJAZ8* OE showed no significant changes. In contrast, *MaFLS2* expression was downregulated in both *MaJAZ5* and *MaJAZ8* OE ECS. Additionally, the expression of B-ring decorating enzyme paralogs (*MaF3’5’H1-7*) showed a marked decrease in *MaJAZ5* and *MaJAZ8* OE banana ECS, with *MaJAZ5* exhibiting stronger repressive activity than *MaJAZ8* (Fig. 10f). Further, we quantified the levels of different flavonol compounds in *MaJAZ5* and *MaJAZ8* OE ECS. Metabolite quantification showed decreased levels of naringenin, kaempferol, quercetin, and rutin. Both MaJAZ5 and MaJAZ8 exhibit repressive activity on *MaTFR,* while only MaJAZ5 represses the *MaHDZIV6* expression. The overexpression of *MaJAZ5* and *MaJAZ8* leads to decreased expression of flavonol biosynthesis genes and metabolite content in banana ECS, highlighting their role in balancing trichome development and flavonoid content accumulation (Fig. S15c).

## Discussion

### Diversity of trichomes in *Musa* species

Trichome development in monocots has been relatively underexplored, with only a few studies reported in rice and maize. In the present work, we reported the presence of trichomes in banana (*Musa spp.*), particularly on the inflorescence stalk. However, their occurrence in banana appears to be restricted to specific cultivars, indicating potential cultivar-specific evolutionary advantages. In tomato, multicellular trichomes are classified into seven types (Types I to VII), each exhibiting distinct morphologies and functions (Simmons & Gurr, 2005). Trichomes are typically categorized based on their structure and function into GTs and NGTs, which can be either unicellular or multicellular. For instance, *A. thaliana* possesses unicellular NGTs, whereas *S. lycopersicum*, *C. sativus*, feature multicellular GTs, and *N. tabacum* has multicellular NGTs (Chalvin *et al*., 2020). Similarly, banana cultivars GN and RB having triploid (AAA) genome display structurally distinct types of trichomes, which are predominantly unicellular, with a few trichomes being multicellular types on the fruit stalk. This pattern suggests a potential evolutionary transition from unicellular to multicellular trichomes, likely as an adaptation to enhance environmental resilience. DPBA staining indicated a significant enrichment of kaempferol derivatives in the TR cultivars, suggesting a higher accumulation of flavonols within their trichomes. This observation was further supported by the elevated expression levels of *MaFLS1* and *MaFLS2* genes in the TR (GN) cultivars, which are key enzymes involved in flavonol biosynthesis. The enhanced presence of these compounds in banana trichomes may contribute to increased tolerance against both biotic and abiotic stresses, potentially providing an adaptive advantage under challenging environmental conditions.

In order to decipher molecular mechanism of trichome development as well as the impact of presence of trichomes on specialized metabolism, we conducted transcriptome analysis of TF and TR banana cultivars and identified key candidates through phylogenetic analysis, transcriptome expression profiles, and validation via qRT-PCR. Our investigation focuses on the roles of *MaTFR* (a R2R3-MYB) and Ma*HDZIV6* in trichome development, utilizing heterologous *GUS* expression and complementation in mutant backgrounds. Additionally, we demonstrate their involvement in flavonoid pathway regulation through transient overexpression and silencing in banana ECS. By targeting of flavonoid promoters (*MaFLS1*, *MaFLS2, MaF3’5’H1* and *MaF3’5’H2* and the protein-protein interaction between *MaTFR* and *MaHDZIV6*, providing comprehensive insights into their functional roles. Further, the variation in JA content between TF and TR cultivars suggest the involvement of JA signaling in trichome development. MaJAZ5 and MaJAZ8 interact with MaTFR, thereby reducing its transcriptional potential in banana. Taken together, we identified key transcriptional regulators which are not only involved in the trichome development but also directly regulate structural genes of flavonoid biosynthesis.

### MaTFR is involved in trichome development and flavonol modulation

Numerous studies have established that specific *MYB* genes play a pivotal role in regulating GT development (Qin *et al*., 2021). In Arabidopsis, several *MYB* genes, including *GL1*, *TRY*, *CPC*, *ETC1*, and *ETC2*, have been implicated in trichome development (Kirik *et al*., 2004; Wang *et al*., 2008; Xu *et al*., 2023). Among these, *GL1* acts as a positive regulator of trichome formation, while *TRY*, *CPC*, and *ETCs* function as negative regulators (Kirik *et al*., 2004). Additionally, *AtMYB1* and *AtMYB61* have been identified as regulators of GT initiation and development in *A. annua* (Matías Hernández *et al*., 2017). AtMYB82 also functions in trichome development, while AtMYB106 and AtMYB16 are involved in trichome initiation (Liang *et al*., 2014). *AtMYB5* is specifically expressed in trichome cells, where it influences trichome size and length (Li *et al*., 2009). These findings underscore the conserved and diverse roles of *MYB* genes in regulating trichome formation across different plant species. Previously, we have shown that MaTFR belongs to general flavonoids and trichome development subclade (Pucker *et al*., 2020). *MaTFR* is a close homolog of *AtMYB5* and shows higher expression in TR (GN and RB) banana cultivars compared to TF cultivars. The *GUS* expression driven by *MaTFR* promoter in Arabidopsis trichomes further supports its potential role in trichome development. Similarly, in cotton, *GhMYB3* has been shown to regulate trichome development in *Arabidopsis*. Wang *et al*., reported that *35S::GL1* could not sustain normal trichome production in T2 and subsequent generations compared to *GL1::GL1*, indicating that the 5′ and 3′ noncoding sequences of *GL1* are essential for proper trichome patterning (Wang *et al*., 2008). Consistent with this, the expression of *MaTFR* under the *35S* promoter partially complemented the trichome phenotype in the trichome-less *gl1* mutant of *Arabidopsis*. Higher expression of *AtF3’H* and *AtFLS1* and increased levels of myricetin and quercetin derivatives were observed in *35S::MaTFR-* expressing lines within the *gl1* mutant background in *A. thaliana*. Similar to MaTFR, in the *Camellia sinensis*, an R2R3-MYB gene, *CsMYB1*, regulates both trichome development and the biosynthesis of galloylated *cis*-catechins (Li *et al*., 2022b). In the homologous system, using banana transient expression in banana ECS, we demonstrated that transient overexpression and silencing of *MaTFR* in banana ECS led to increased and decreased expression of *MaFLS1, MaF3’5’H1,* and *MaF3’5’H2,* respectively, confirming its regulatory role in flavonoid biosynthesis. Transactivation and binding assays confirmed that MaTFR directly regulates the structural genes involved in the flavonol biosynthesis in banana. In Arabidopsis, AtGL1 acts as an upstream regulator of AtGL2 in trichome development (Chalvin *et al*., 2020) suggesting that MaTFR may have a similar regulatory role in banana trichome formation. Thus, based on multiple experimental evidences involving expression pattern of *MaTFR* in different cultivars differing in trichome development, complementation assay, transient expression and promoter transactivation assay, we established that MaTFR is a regulator of trichome development and flavonoid biosynthesis in banana.

### MaHDZIV6 involved in trichome development and flavonol modulation

HDZIP IV TFs play a vital role in regulating plant growth, development, metabolism, and stress responses (Schrick *et al*., 2023). The HDZIP IV preferentially displays expression in the outer cell layers such as epidermal cells of plant organs (Chew *et al*., 2013). In *A. annua*, *AaHD8* and *AaHD1* can directly regulate to trichome initiation and cuticle development. Previously identified regulators of tomato trichomes belong to the HDZIP family, including AtGL2, AaHD8, and Woolly (Yang *et al*., 2011; Khosla *et al*., 2014; Yan *et al*., 2018). Beyond tomato, the HDZIP TF GhHOX3 in cotton has also been shown to regulate fiber length(Shan *et al*., 2014). *GLABROUS* (*CmGL*) encodes a HDZIP IV TF playing roles in multicellular trichome initiation in melon (Zhu *et al*., 2018). GhHOX4 an HDZIP-IV TF influences fibre elongation by interacting with phosphatidic acid in cotton plants (Wang *et al*., 2024). *NtHD9* and *NtHD12* members synergistically participate in the development of long GTs in response to JA signaling in *N. tabacum* (Zhang *et al*., 2023). Based on these findings, we hypothesized that HDZIP-IV family transcription factor might be involved in the trichome development in banana. To this end, two HDZIP-IV family TFs, namely two MaHDZIV6 and MaHDZIV8 were found to be putative orthologs of rice OsGL2 and Artemisia AaHD1. Further, our study showed that the expression of *MaHDZIV6* is upregulated in the trichome bearing cultivars as compared to trichome-less banana cultivars, strengthening the assumption that MaHDZIV6 is a regulator of trichome development in banana. The role of MaHDZIV6 in the trichome development was confirmed by its complementation study in *gl2* mutant background. *MaHDZIV6* partially rescued the trichome phenotype in the trichome-less *gl2* mutant of *Arabidopsis*, indicating its functional role in trichome development.

Trichome development is a complex process governed by multiple genes, and the partial complementation observed could be attributed to the use of the *35S* promoter instead of the native *MaHDZIV6* promoter, which may have influenced gene regulation. Another plausible reason is the evolutionary distance between banana (a monocot) and Arabidopsis (a dicot), which could limit the functional compatibility of trichome regulatory networks. GLABRA2, an HDZIP IV family TF acts as an activator for proanthocyanidin biosynthesis in *Medicago truncatula* seed coat (Gu *et al*., 2024). Similarly, MaHDZIV6 increased the expression of *AtF3’H* and *AtFLS1* in Arabidopsis, indicating its involvement in the regulation of flavonol biosynthesis. Likewise, in the present study, we demonstrated that MaHDZIV6 is a regulator of flavonol biosynthesis. Our findings are based on firm experimental supports, including transient expression in banana ECS and promoter-transactivation assay, Y1H and ChIP experiments. These results establish MaHDZIV6 as a regulator of trichome development and flavonoid biosynthesis in banana. Further, we also reported that MaHDZIV6 comes under the regulon of MaTFR, which was also demonstrated to be the regulator of flavonoid biosynthesis and trichome development in banana. Thus, our study reveals that a module of MaTFR and MaHDZIV6 fine-tunes trichome development and flavonoid biosynthesis in banana.

### JA coordinates the trichome development and flavonol biosynthesis in banana

Jasmonates are signalling molecules, which act as positive regulators of trichome development and specialized metabolism in diverse plant species (Li *et al*., 2022c). For example, Methyl JA treatment enhances trichome density in *A. thaliana*, *S. lycopersicum*, and *A. annua*. Which possess GTs and NGTs (Han *et al*., 2022; Hua *et al*., 2022). Methyl JA enhances the flavonoid accumulation of phenolic compounds in pear and *Castilleja tenuiflora respectively* (Rubio Rodríguez *et al*., 2019; Premathilake *et al*., 2020). In our study, transcriptome data initially revealed that JA responses were one of the key hormonal responses in TR cultivars compared to TF ones. Further, naturally high JA-Ile and JA content in TR (GN and RB) cultivars hint the involvement of JA in trichome development in banana.

A key limitation in our banana system is that trichomes appear only on the fruit stalk at mature stage, posing challenge to measure trichome density after JA treatments. Therefore, we studied the effect of JA on banana plantlet in the context of its effect over the trichome-related genes. To this end, we observed that *MaTFR* expression increased significantly following methyl JA treatment in seedlings, with peak expression at 12 and 24 h. Similarly, *MaHDZIV6* expression was found to be up regulated in response to methyl JA treatment. In our study, we also observed that methyl JA treatment enhanced the expression of *MaCHS* paralogs, *MaCHI2, MaFLS1/2,* and *MaF3’5’H* paralogs at different time points. As a result, the levels of key flavonoids such as kaempferol, myricetin, quercetin, and their glycosylated forms, including rutin, increased. Recently, it has been reported that methyl JA induces the accumulation of phenylphenalenone, contributing to resistance against black leaf streak disease in banana.

The overexpression of *SlJAZ2* in tomato was reported to reduce trichome numbers (Yu *et al*., 2018). Similarly, in cotton, overexpression of *GhJAZ2*, a repressor of the JA signaling pathway, was found to inhibit fiber elongation (Hu *et al*., 2016). The allocation of metabolic resources toward producing plant defense compounds is often linked to reduced growth and biomass accumulation. GA and JA have both antagonistic and synergistic effects on plant growth in *Arabidopsis*. JAZ proteins and DELLA, a repressor in the GA signaling pathway, interact with the WD-repeat/bHLH/MYB complex to regulate trichome initiation (Qi *et al*., 2014). Additionally, exogenous JA treatment has been shown to increase both trichome density and the number of leaves in *Arabidopsis* (Traw & Bergelson, 2003). In Arabidopsis, JAZ proteins interact with the R2R3-MYB TFs MYB21 and MYB24 to regulate jasmonate-mediated stamen development (Song *et al*., 2011). Our transcriptome analysis revealed five differentially regulated MaJAZ repressors. However, only MaJAZ5 and MaJAZ8 showed interactions with MaTFR, while none of the MaJAZ proteins interacted with MaHDZIV6. Interestingly, in transactivation assays, the addition of MaJAZ5 and MaJAZ8 suppressed the transcriptional activity of *MaFLS1, MaF3’5’H1, MaF3’5’H2,* and the M*aHDZIV6* promoters. Furthermore, in banana ECS transiently OE *MaJAZ5* and *MaJAZ8,* the expression levels of *MaTFR, MaHDZIV6,* and flavonol biosynthetic genes, along with flavonol content, were reduced compared to the empty vector (EV) control. This highlights a striking functional conservation of the JAZ–MYB regulatory module from dicots to monocots in plant systems. Flavonoids such as kaempferol and quercetin have been implicated in stress responses and photoprotection (Naik *et al*., 2022).

Based on our findings, we propose a regulatory model illustrating how transcriptional regulators and JA signaling are integrated to control trichome development and flavonoid biosynthesis in banana (*Musa spp.)* (Fig. 11). Our study identifies MaTFR and MaHDZIV6 as key regulators of trichome development and flavonol biosynthesis in banana. MaTFR enhances the expression of MaHDZIV6 and flavonol biosynthetic genes, reinforcing its role in trichome formation.

**Fig. 11.**
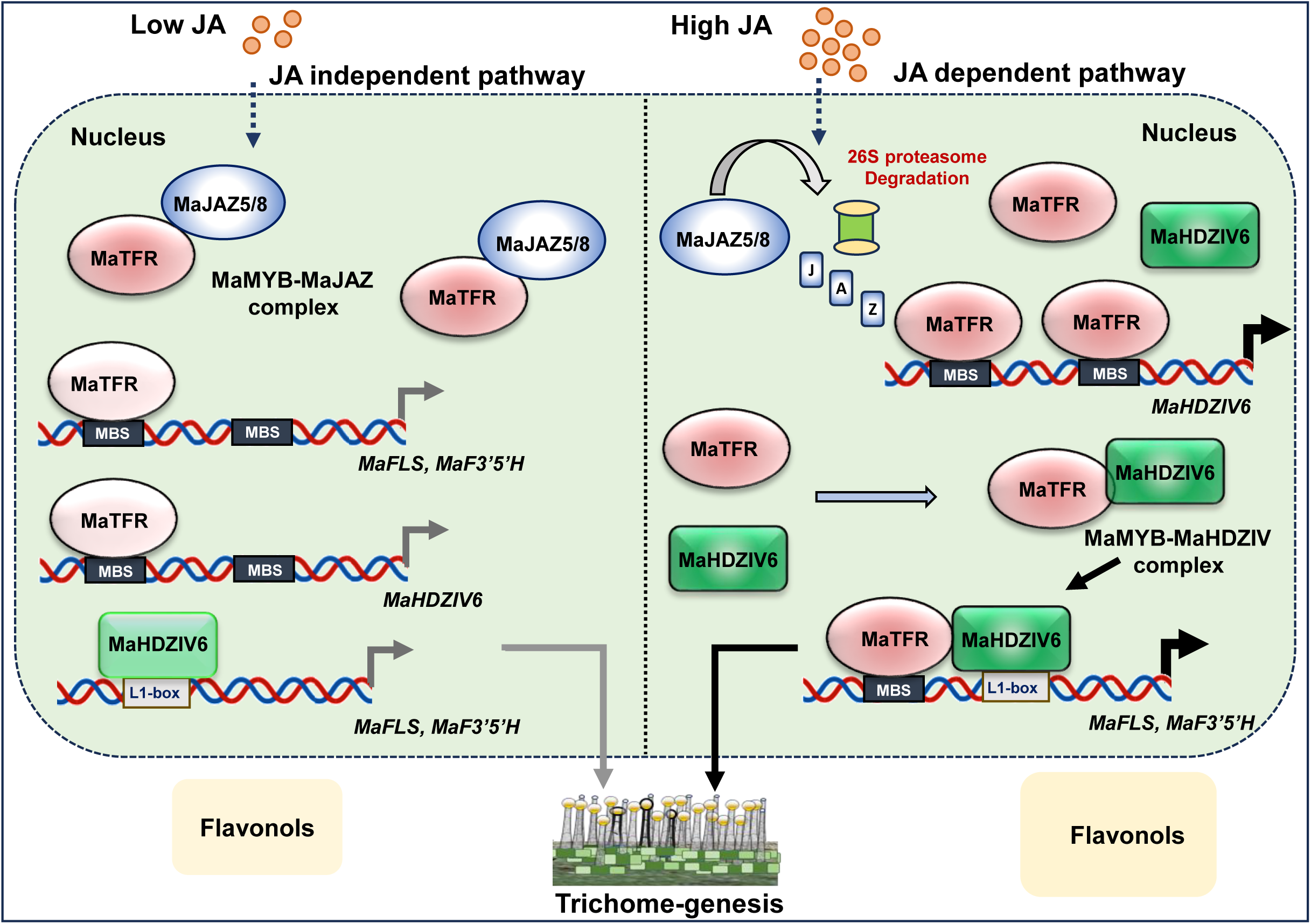
The proposed model of JA-induced regulation of trichome development and flavonoid biosynthesis in *Musa* species. Working model illustrating the regulatory network involving MaTFR, MaHDZIV6, and MaJAZ repressors in jasmonic acid (JA)-mediated trichome development and flavonoid biosynthesis in banana. In TF cultivars, endogenous active levels of JA are low, allowing enhanced level of MaJAZ repressor, which interact with MaTFR to form MYB-JAZ complex. It reduces the binding of MaTFR to *proMaFLS*, *proMaF3 5 H* and *proMaHDZIV6* resulting in reduced flavonol content and limiting trichome development. When JA levels are high, MaJAZ undergoes degradation, releasing MaTFR from the MYB-JAZ complex. This strengthens MaTFR’s transactivation activity toward MaHDZIV6 and flavonol promoters, positively regulating trichome development and flavonoid pathway. Overall, this study provides a detailed model of the JA-mediated transcriptional regulatory module controlling trichome development and flavonol pathway in banana, highlighting the complex interplay between transcriptional activators and repressors in response to JA signaling.

Similarly, MaHDZIV6 also promotes the expression of flavonol biosynthetic genes. Interestingly, MaTFR and MaHDZIV6 physically interact at the protein level, strengthening their functional activity in regulating both trichome development and flavonol biosynthesis. Additionally, JA quantification and methyl JA treatment suggest that JA is involved in trichome development. We also report that two MaJAZ5 and MaJAZ8 interact with MaTFR, thereby reducing its transcriptional potential and modulating its regulatory activity over flavonoid biosynthetic genes. This suggests that the regulation of MaTFR–MaHDZIV6 module is fine-tuned by JA signalling, where JAZ proteins act as negative regulators under stress or developmental cues. Overall, the MaTFR–MaHDZIV6–JAZ regulatory module plays a pivotal role in coordinating trichome development and flavonoid biosynthesis in banana. These findings will provide deeper insights into the evolutionary and functional significance of trichomes in monocots, expanding our understanding of plant defence and specialized metabolism. A better understanding of the molecular mechanisms that control the biosynthesis of special metabolites in trichomes will be important for future stress adaptations and commercial biopharming applications.

## Authors’ contributions

AP conceived the idea and designed the research. SS and ST conducted experiments. SS, ST, and AP interpreted the data. SS and ST wrote the manuscript. AP and PM provided insightful suggestions during discussion. AP and PM finalized the manuscript. AP coordinated the research project. All authors read and approved the final manuscript.

## Acknowledgements

This work was supported by the core grant from the National Institute of Plant Genome Research and the research grants by Department of Biotechnology, Government of India, to AP (BT/PR36694/NNT/28/1722/2020 and BT/PR38402/GET/119/308/2020). SS acknowledges the University Grants Commission (UGC), Government of India, for the Senior Research Fellowship. ST acknowledges the ANRF-National Post Doctoral Fellowship (N-PDF). The authors are thankful to the DBT-eLibrary Consortium (DeLCON) for providing access to e-resources. We acknowledge the Metabolome facility at NIPGR for phytochemical analysis.

## Conflict of interest

The authors declare no conflict of interest.

## Data availability statement

All data supporting the findings of this study are available within the paper and within the supplemental data published online. The raw sequencing data generated in this study have been deposited in the NCBI SRA under the Bio-Project accession number PRJNA1274196.

**Fig. S1 Differential gene expression and MAPMAN analysis in TF and TR cultivars. (a)** Volcano plots showing the log2(fold change) differentially up- and down regulated genes in PLT vs GN, PLT vs RB and GN vs RB. **(b)** MapMan visualization of pathway enrichment for polyketide/acetate pathway of secondary metabolism in three banana cultivars: PLT, GN, and RB. Data are based on differentially expressed genes mapped to corresponding pathways, providing insights into cultivar-specific modulation of secondary metabolism. Red coloured boxes represent the level of enrichment or expression change.

**Fig. S2 Defence hormone-related gene expression in TF and TR cultivars.** Heatmaps showing expression of defence hormone biosynthetic and signalling related genes **(a)** abscisic acid (ABA), **(b)** ethylene (ET), **(c)** salicylic acid (SA) and **(d)** jasmonic acid (JA). Colour bar scale represents log2 normalized gene expression and aqua blue to magenta corresponds to low to high expression.

**Fig. S3 Gene expression profiling of different transcription factor families.** Heatmaps representing the log2 normalized expression of **(a)** *MabHLH***, (b)** *MaWRKY,* **(c)** *MaHDZIP-IV*, **(d)** *MaC2H2* transcription factor genes, **(e)** selected *MaMYBs* putatively known for their role in trichome formation and development, and **(f)** other putative trichome specific genes in banana sharing homology with known trichome specific regulators in other plant species. Colour bar scale represents normalized expression values, with aqua blue indicating low expression, and magenta indicating high expression.

**Fig. S4 Expression profiling of flavonoids biosynthesis pathway genes. (a)** Pictorial representation of flavonoids biosynthesis pathway along with the heatmaps showing the log2 normalized expression of biosynthetic genes involved at different steps. Colour bar scale (aqua blue to magenta) indicate low expression to high expression. **(b)** Relative expression analysis of different flavonoids biosynthesis pathway genes by RT-qPCR. Standard deviation and significance level are represented by error bars and asterisks, respectively.

**Fig. S5 Flavonoids content in TF and TR cultivars. (a)** Bar graphs showing the differential content of various phenylpropanoids, such as *p*-coumaric acid, cinnamic acid, and syringic acid; and flavonoids including flavonols and dihydro-flavonols such as kaempferol, myricetin, dihydromyricetin, rutin hydrate, and kaempferol derivatives in the TR (GN and RB) and the TF (PLT) cultivars. **(b)** Proanthocyanidin content (catechin, epicatechin gallate and epigallocatechin) in TR (GN and RB) and TF (PLT) cultivars. **(c)** Bar graph showing the total anthocyanin content enriched significantly in TR (RB) cultivar.

**Fig. S6 Gene expression analysis of selected *MaMYB* transcription factors in PLT, GN and RB.** Expression analysis of *MaMYB* genes associated with general flavonoid and trichome regulation revealed that gene IDs *Macma4_06_g16640.1*, *Macma4_02_g22860.1*, *Macma4_09_g11600.1*, *Macma4_05_g06590.1*, and *Macma4_09_g27080.1* were highly expressed in TR (GN and RB) tissues. In contrast, *Macma4_06_g40680.1* and *Macma4_07_g20460.1* showed higher expression levels in TF tissues compared to TR tissues.

**Fig. S7 Expression and metabolite analysis in *WT, gl1,* and *MaTFR* overexpressing *gl1* complemented lines. (a)** RT-qPCR analysis showed increased relative expression of flavonoid biosynthetic genes (*AtF3’H* and *AtFLS1*) in *MaTFR*-complemented lines of *gl1* mutant of Arabidopsis. **(b)** UHPLC-based flavonol quantification of quercetin, myricetin and kaempferol showing enhanced levels of myricetin and quercetin in the complemented lines.

**Fig. S8 Identification of *MaHDZIV* genes and expression analysis in *MaTFR*-overexpressing and silencing ECS. (a)** The phylogenetic tree illustrating the relationships among all MaHDZIV transcription factors from banana and trichome-related HDZIV from other plant species and formed distinct clades. The MaHDZIV6 showed high evolutionary relatedness to OsGL2-type from rice, ZmOCL4 from *Zea mays*, GaHOX2 from *Gossypium arboreum* and AaHD1 from *Artemisia annua* and highlighted in green. Branch lengths are marked on each branch, and bootstrap support is represented by pink bubble sizes at the nodes. **(b)** Bar graphs showing the qRT-PCR based expression analysis of *MaHDZIV6*, *MaHDZIV8* and *MaHDZIV16*. **(c)** Transcript levels of *MaHDZIV6*, *MaHDZIV8* and *MaHDZIV16*, where *MaHDZIV6* and *MaHDZIV8* were significantly upregulated in *MaTFR*-OE ECS. Significance level is indicated by number of asterisks.

**Fig. S9 Expression and metabolite analysis in *WT, gl1,* and *MaHDZIV6* overexpressing *gl2* complemented lines. (a)** Bar graph displaying themodulation in expression of flavonoid biosynthetic genes (*AtF3’H* and *AtFLS1*) in *MaHDZIV6*-complemented lines in the *gl2* mutant of Arabidopsis by RT-qPCR analysis. **(b)** Increased flavonol (quercetin, myricetin and kaempferol) content in complemented lines (L2 and L8) compared to *gl2* mutant of Arabidopsis.

**Fig. S10 Estimation of defense hormones in PLT, GN and RB cultivars by LC-MS. (a)** Bar graphs showing differential accumulation of ABA and SA across TF (PLT) and TR (GN and RB) cultivars. **(b)** The levels of JA-Ile and jasmonic acid levels were higher in TR cultivars, while cis-OPDA was more abundant in the TF cultivar.

**Fig. S11 Relative expression analysis of regulatory genes in methyl JA treated banana seedlings**. **(a)** Representative image of banana seedlings used as mock and treated with 100µM methyl jasmonate (JA) for 0, 6, 12, 24, 48, and 72 h. **(b)** Relative expression analysis of *MaTFR* and *MaHDZIV6* in mock and methyl JA treated banana seedlings at 0, 6, 12, 24, 48, and 72 h by RT-qPCR. Values are shown as means ± SD of three biological replicates.

**Fig. S12 Relative expression analysis of flavonol biosynthesis genes in methyl JA treated banana seedlings**. **(a)** RT-qPCR based relative expression level of *MaCHS1*, *MaCHS2*, *MaCHS3, MaCHS4, MaCHS6*, *MaCHI2*, *MaFLS1-2,* and *MaF3’5’H1, MaF3’5’H2, MaF3’5’H4, MaF3’5’H5,* and *MaF3’5’H6,* and **(b)** flavonols quantification (kaempferol, myricetin, quercetin and rutin) at different time points (0, 6, 12, 24, 48, 72 h) of control (mock) and JA treated. Data represents means ± SD of three biological replicates.

**Fig. S13 Phylogeny and expression analysis of *MaJAZ* in banana. (a)** The phylogenetic tree was constructed using the maximum likelihood method in MEGA-X software, with 1,000 bootstrap values. The resulting tree was visualized using iTOL v6.4 (http://itol.embl.de/) software. **(b)** Heatmap showing the log2 normalized expression of *MaJAZ*s in TF (PLT) and TR (GN and RB) cultivars. The hierarchical clustering was done on the basis of expression, and aqua blue to magenta corresponds to low to high expression. **(c)** Relative expression analysis of selected *MaJAZ* repressors in banana revealed differential expression patterns between TF and TR cultivars using RT-qPCR.

**Fig. S14 Protein-protein interaction between MaHZIV6 with MaJAZs repressors and subcellular localization. (a)** Pictorial representation depicting protein–protein interactions between MaHDZIV6 and five selected MaJAZ proteins (MaJAZ5, MaJAZ8, MaJAZ9, MaJAZ20, and MaJAZ21) using yeast two-hybrid (Y2H) screening. No interaction was detected between MaHDZIV6 and any of the tested MaJAZ proteins. **(b)** The subcellular localization of MaJAZ5 and MaJAZ8 was analyzed in *N. benthamiana* leaves. The YFP-tagged MaJAZ fusion proteins were observed via confocal microscopy and were found to localize specifically to the nucleus, co-localizing with the nuclear marker NLS-RFP. An EV (pSITE-3CA) expressing free YFP was used as a negative control. Scale bar: 50µm.

**Fig. S15 Flavonol content in *MaJAZ5* and *MaJAZ8* transiently overexpressing banana ECS.** Bar graphs showing the flavonol content in EV, *MaJAZ5* and *MaJAZ8* transiently overexpressing tissues. The aglycone forms of flavonols, kaempferol and quercetin, were significantly reduced in *MaJAZ5* and *MaJAZ8* OE banana ECS compared to the EV control. Similarly, glycan forms (naringenin and rutin) levels were also lower in *MaJAZ5* and *MaJAZ8* OE banana ECS compared to the EV vector control.

**Table S1.** List of primers used in the study.

**Dataset S1. Differential gene expression profiles across PLT, GN and RB epidermis.** The results of differential gene expression analysis conducted across three pairwise epidermal tissue comparisons: PLT vs GN, PLT vs RB and GN vs RB. RNA-seq data were processed and analyzed using the DESeq2 to estimate expression fold changes and statistical significance. Genes exhibiting a false discovery rate (FDR)-adjusted *p*-value < 0.05 and an absolute log2 fold change ≥ 1 were considered differentially expressed. For each tissue comparison, differentially expressed genes (DEGs) are classified as upregulated or downregulated based on expression changes relative to the first condition in the pairwise analysis.

**Dataset S2. Gene-ontology (GO) enrichment analysis of DEGs.** The GO enrichment analysis of downregulated and upregulated DEGs genes across three pairwise comparisons: (1) PLT vs GN, (2) PLT vs RB, and (3) GN vs RB.

## Notes

### Competing Interest Statement

The authors have declared no competing interest.

