## Supplementary Figures for "Coordinated regulation of trichome morphogenesis and flavonoid pathway by a MYB–HDZIP–JAZ module in banana (*Musa* sp.)"

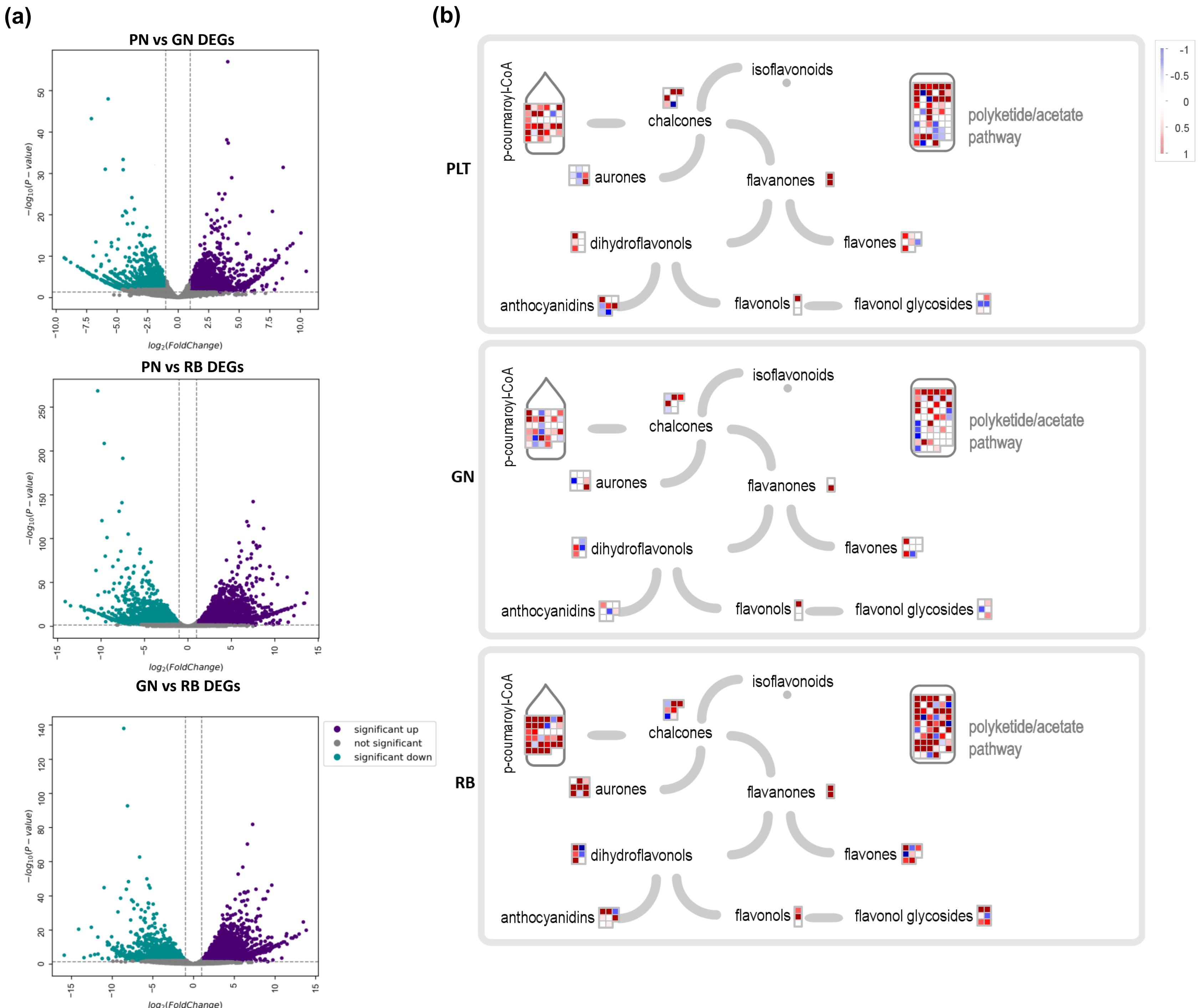
(a)
PN vs GN DEGs
PN vs RB DEGs
GN vs RB DEGs
−log10(P − value)
log2(FoldChange)
significant up
not significant
significant down
(b)
PLT
GN
RB
p-coumaroyl-CoA
chalcones
isoflavonoids
aurones
flavanones
dihydroflavonols
flavones
anthocyanidins
flavonols
flavonol glycosides
polyketide/acetate pathway
-1
-0.5
0
0.5
1

Fig. S1

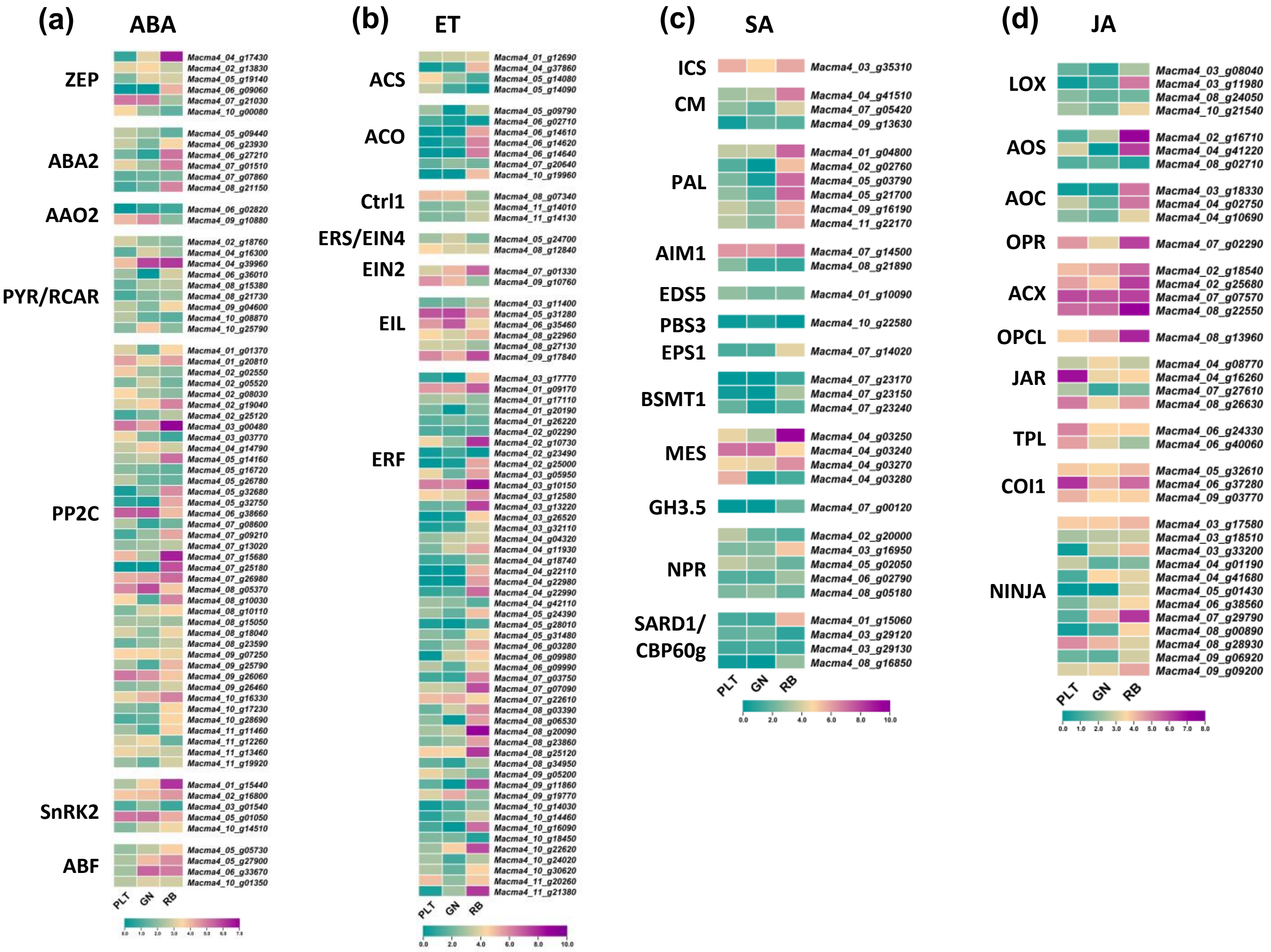

(a)
ABA
ZEP
ABA2
AAO2
PYR/RCAR
PP2C
SnRK2
ABF
PLT
GN
RB
(b)
ET
ACS
ACO
Ctrl1
ERS/EIN4
EIN2
EIL
ERF
PLT
GN
RB
(c)
SA
ICS
CM
PAL
AIM1
EDS5
PBS3
EPS1
BSMT1
MES
GH3.5
NPR
SARD1/
CBP60g
PLT
GN
RB
(d)
JA
LOX
AOS
AOC
OPR
ACX
OPCL
JAR
TPL
COI1
NINJA
PLT
GN
RB


Fig. S2

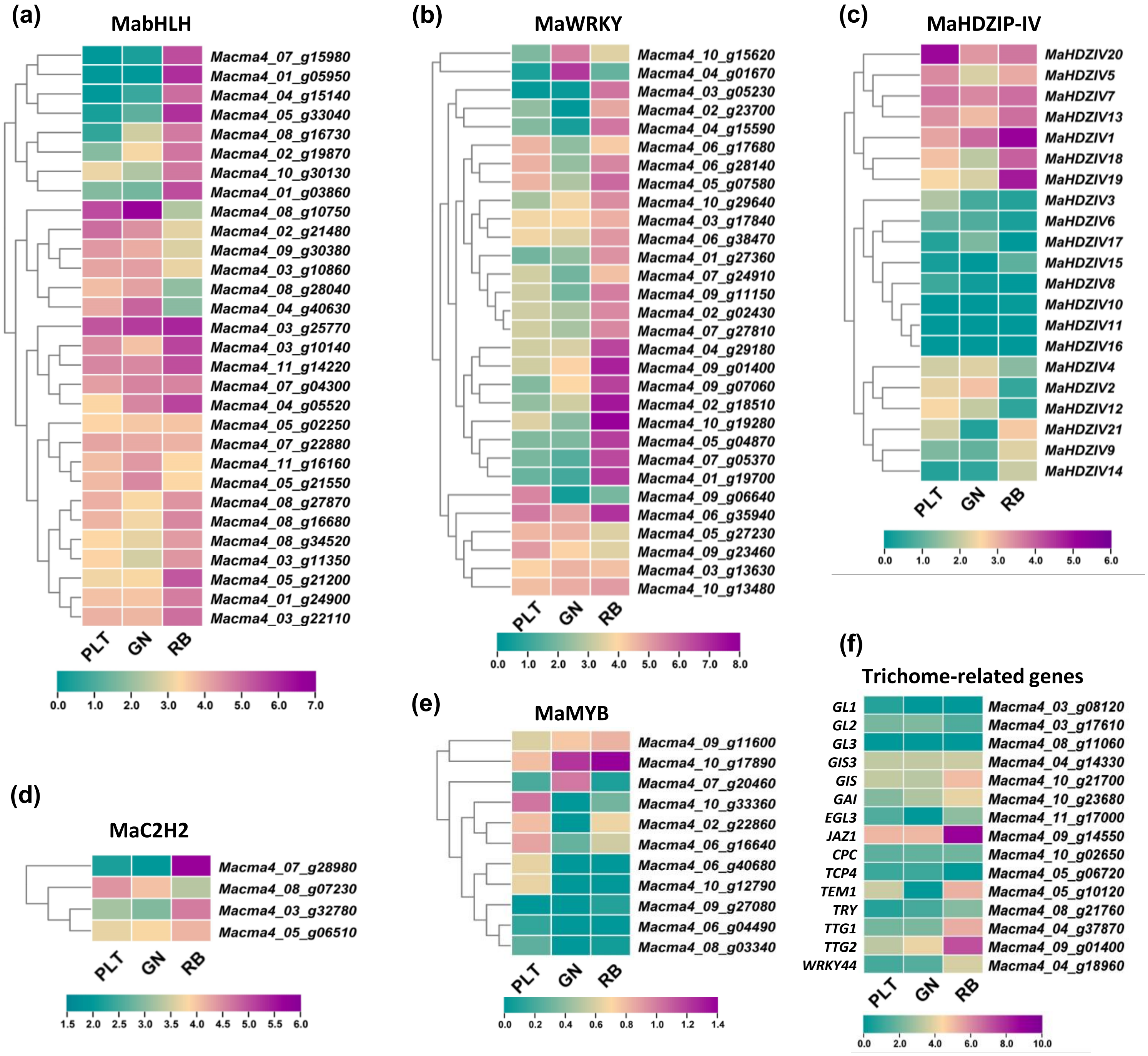
(a)
MabHLH
Macma4_07_g15980
Macma4_01_g05950
Macma4_04_g15140
Macma4_05_g33040
Macma4_08_g16730
Macma4_02_g19870
Macma4_10_g30130
Macma4_01_g03860
Macma4_08_g10750
Macma4_02_g21480
Macma4_09_g30380
Macma4_03_g10860
Macma4_08_g28040
Macma4_04_g40630
Macma4_03_g25770
Macma4_03_g10140
Macma4_11_g14220
Macma4_07_g04300
Macma4_04_g05520
Macma4_05_g02250
Macma4_07_g22880
Macma4_11_g16160
Macma4_05_g21550
Macma4_08_g27870
Macma4_08_g16680
Macma4_08_g34520
Macma4_03_g11350
Macma4_05_g21200
Macma4_01_g24900
Macma4_03_g22110
PLT
GN
RB
0.0 1.0 2.0 3.0 4.0 5.0 6.0 7.0
(b)
MaWRKY
Macma4_10_g15620
Macma4_04_g01670
Macma4_03_g05230
Macma4_02_g23700
Macma4_04_g15590
Macma4_06_g17680
Macma4_06_g28140
Macma4_05_g07580
Macma4_10_g29640
Macma4_03_g17840
Macma4_06_g38470
Macma4_01_g27360
Macma4_07_g24910
Macma4_09_g11150
Macma4_02_g02430
Macma4_07_g27810
Macma4_04_g29180
Macma4_09_g01400
Macma4_09_g07060
Macma4_02_g18510
Macma4_10_g19280
Macma4_05_g04870
Macma4_07_g05370
Macma4_01_g19700
Macma4_09_g06640
Macma4_06_g35940
Macma4_05_g27230
Macma4_09_g23460
Macma4_03_g13630
Macma4_10_g13480
PLT
GN
RB
0.0 1.0 2.0 3.0 4.0 5.0 6.0 7.0 8.0
(c)
MaHDZIP-IV
MaHDZIV20
MaHDZIV5
MaHDZIV7
MaHDZIV13
MaHDZIV1
MaHDZIV18
MaHDZIV19
MaHDZIV3
MaHDZIV6
MaHDZIV17
MaHDZIV15
MaHDZIV8
MaHDZIV10
MaHDZIV11
MaHDZIV16
MaHDZIV4
MaHDZIV2
MaHDZIV12
MaHDZIV21
MaHDZIV9
MaHDZIV14
PLT
GN
RB
0.0 1.0 2.0 3.0 4.0 5.0 6.0
(d)
MaC2H2
Macma4_07_g28980
Macma4_08_g07230
Macma4_03_g32780
Macma4_05_g06510
PLT
GN
RB
1.5 2.0 2.5 3.0 3.5 4.0 4.5 5.0 5.5 6.0
(e)
MaMYB
Macma4_09_g11600
Macma4_10_g17890
Macma4_07_g20460
Macma4_10_g33360
Macma4_02_g22860
Macma4_06_g16640
Macma4_06_g40680
Macma4_10_g12790
Macma4_09_g27080
Macma4_06_g04490
Macma4_08_g03340
PLT
GN
RB
0.0 0.2 0.4 0.6 0.8 1.0 1.2 1.4
(f)
Trichome-related genes
GL1 Macma4_03_g08120
GL2 Macma4_03_g17610
GL3 Macma4_08_g11060
GIS3 Macma4_04_g14330
GIS Macma4_10_g21700
GAI Macma4_10_g23680
EGL3 Macma4_11_g17000
JAZ1 Macma4_09_g14550
CPC Macma4_10_g02650
TCP4 Macma4_05_g06720
TEM1 Macma4_05_g10120
TRY Macma4_08_g21760
TTG1 Macma4_04_g37870
TTG2 Macma4_09_g01400
WRKY44 Macma4_04_g18960
PLT
GN
RB
0.0 2.0 4.0 6.0 8.0 10.0

Fig. S3

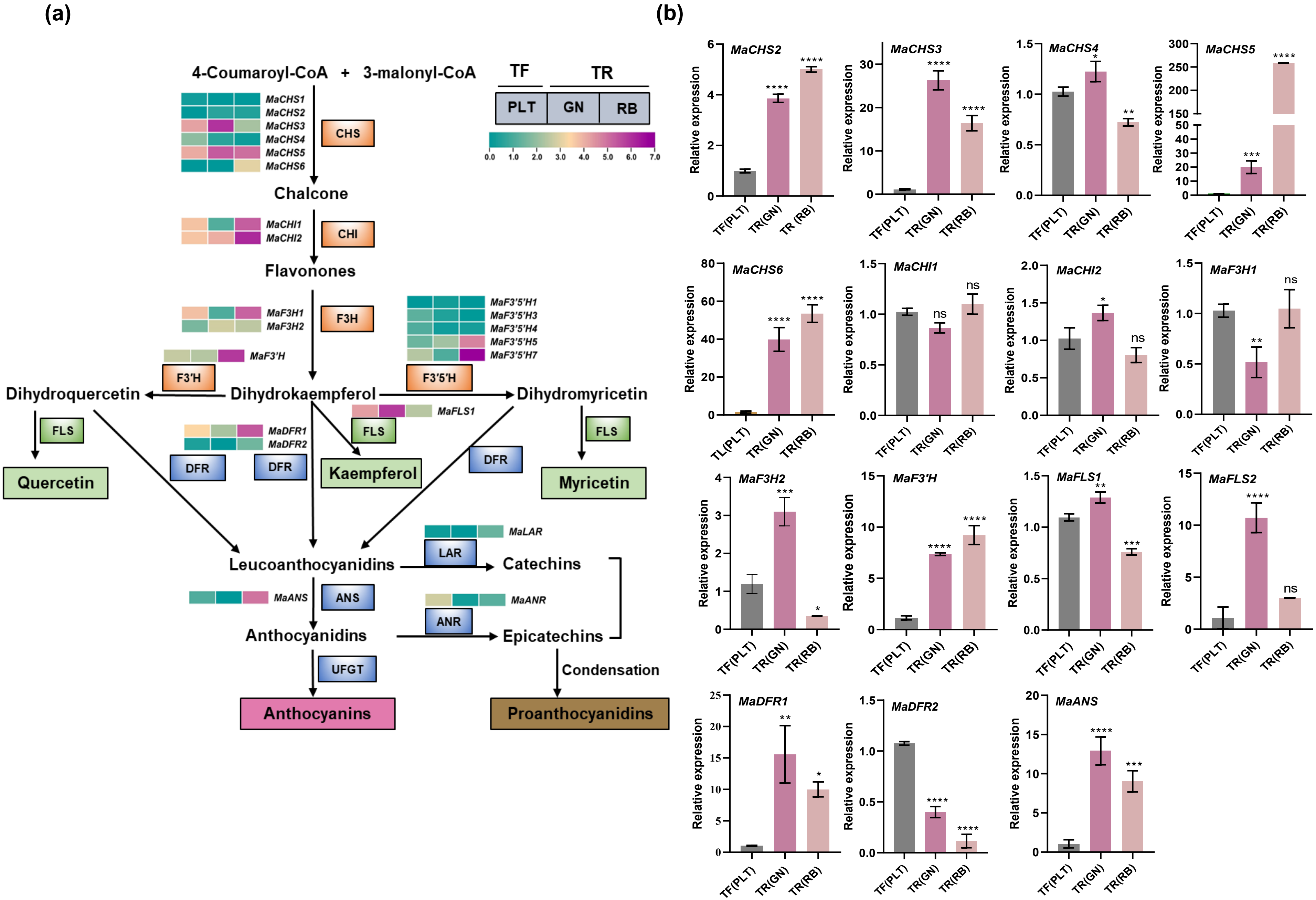
(a)
4-Coumaroyl-CoA + 3-malonyl-CoA
TF
TR
PLT
GN
RB
0.0 1.0 2.0 3.0 4.0 5.0 6.0 7.0
MaCHS1
MaCHS2
MaCHS3
MaCHS4
MaCHS5
MaCHS6
CHS
Chalcone
MaCHI1
MaCHI2
CHI
Flavonones
MaF3H1
MaF3H2
F3H
MaF3'5'H1
MaF3'5'H3
MaF3'5'H4
MaF3'5'H5
MaF3'5'H7
MaF3'H
F3'H
F3'5'H
Dihydroquercetin
Dihydrokaempferol
Dihydromyricetin
MaFLS1
FLS
FLS
FLS
MaDFR1
MaDFR2
DFR
DFR
DFR
Quercetin
Kaempferol
Myricetin
MaLAR
LAR
Leucoanthocyanidins
Catechins
MaANS
ANS
MaANR
ANR
Anthocyanidins
Epicatechins
UFGT
Condensation
Anthocyanins
Proanthocyanidins
(b)
MaCHS2
MaCHS3
MaCHS4
MaCHS5
MaCHS6
MaCHI1
MaCHI2
MaF3H1
MaF3H2
MaF3'H
MaFLS1
MaFLS2
MaDFR1
MaDFR2
MaANS
Relative expression
TF(PLT)
TR(GN)
TR(RB)
TR (RB)
TL(PLT)
ns
Fig. S4

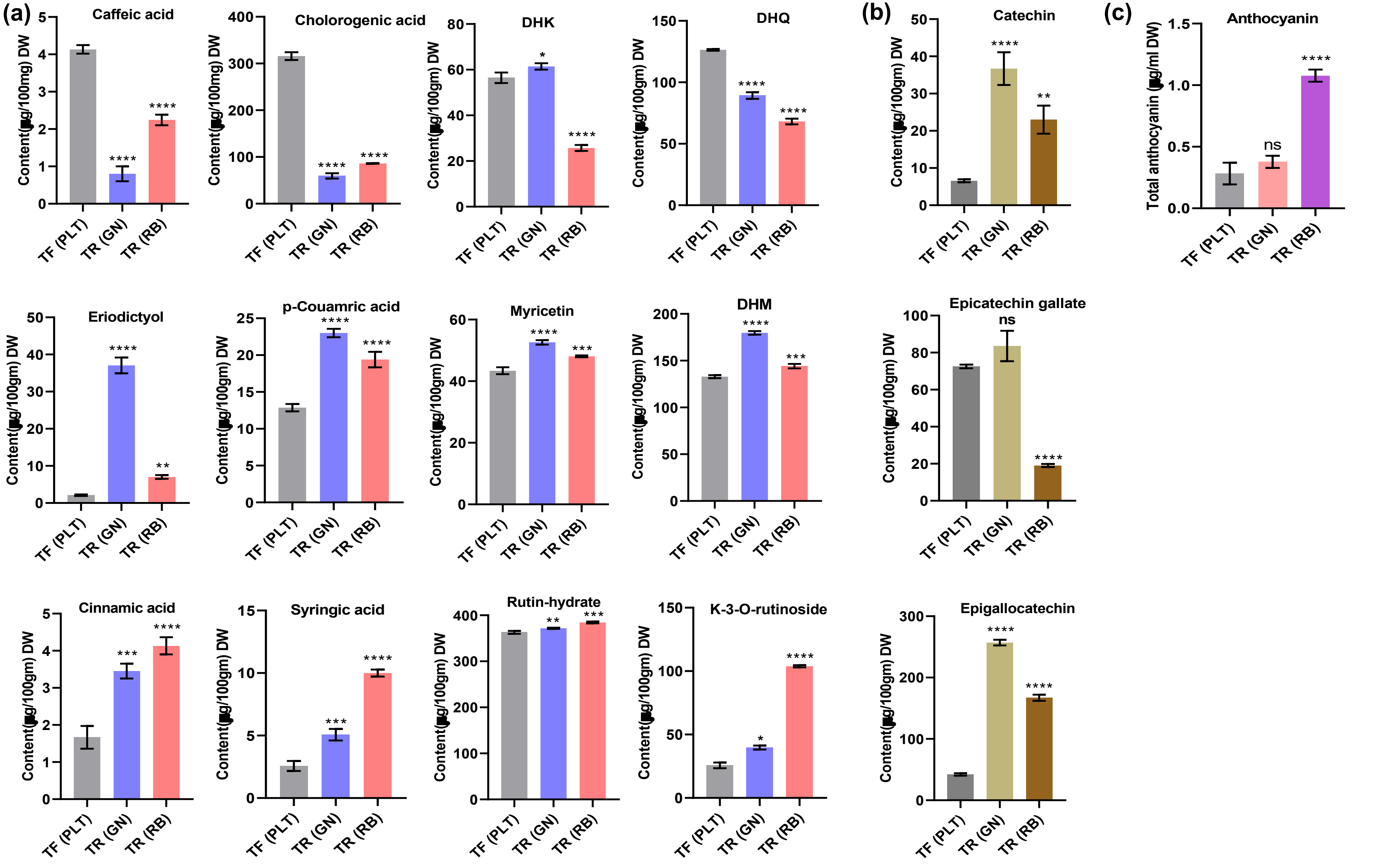
(a)
Caffeic acid
Content(μg/100mg) DW
Cholorogenic acid
Content(μg/100mg) DW
DHK
Content(μg/100gm) DW
DHQ
Content(μg/100gm) DW
(b)
Catechin
Content(μg/100gm) DW
(c)
Anthocyanin
Total anthocyanin (μg/ml DW)
ns
Eriodictyol
Content(μg/100gm) DW
p-Couamric acid
Content(μg/100gm) DW
Myricetin
Content(μg/100gm) DW
DHM
Content(μg/100gm) DW
Epicatechin gallate
ns
Content(μg/100gm) DW
Cinnamic acid
Content(μg/100gm) DW
Syringic acid
Content(μg/100gm) DW
Rutin-hydrate
Content(μg/100gm) DW
K-3-O-rutinoside
Content(μg/100gm) DW
Epigallocatechin
Content(μg/100gm) DW
TF (PLT)
TR (GN)
TR (RB)

Fig. S5

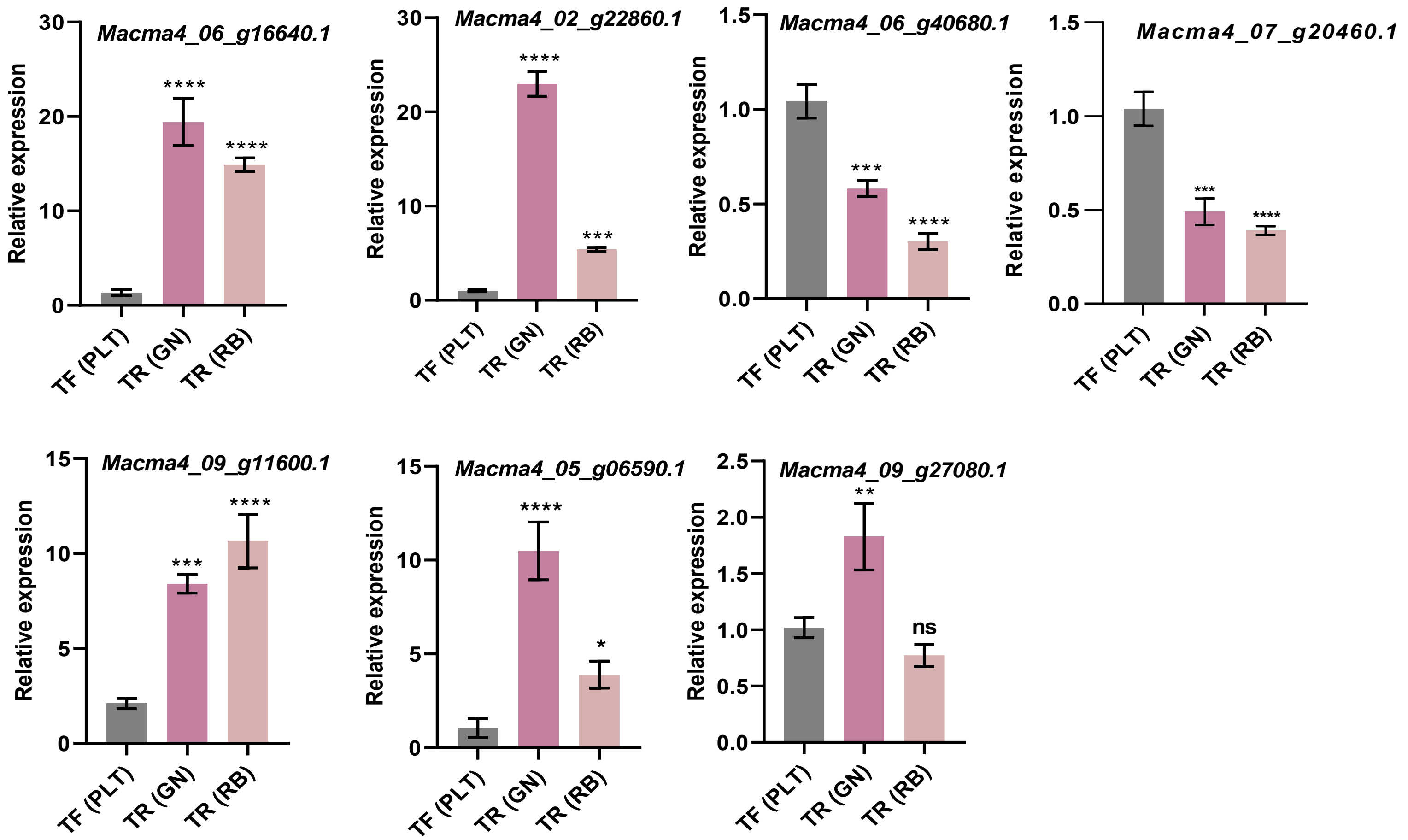
Macma4_06_g16640.1
Macma4_02_g22860.1
Macma4_06_g40680.1
Macma4_07_g20460.1
Macma4_09_g11600.1
Macma4_05_g06590.1
Macma4_09_g27080.1
Relative expression
TF (PLT)
TR (GN)
TR (RB)
****
***
**
*
ns
0
5
10
15
20
30
0.0
0.5
1.0
1.5
2.0
2.5

Fig. S6

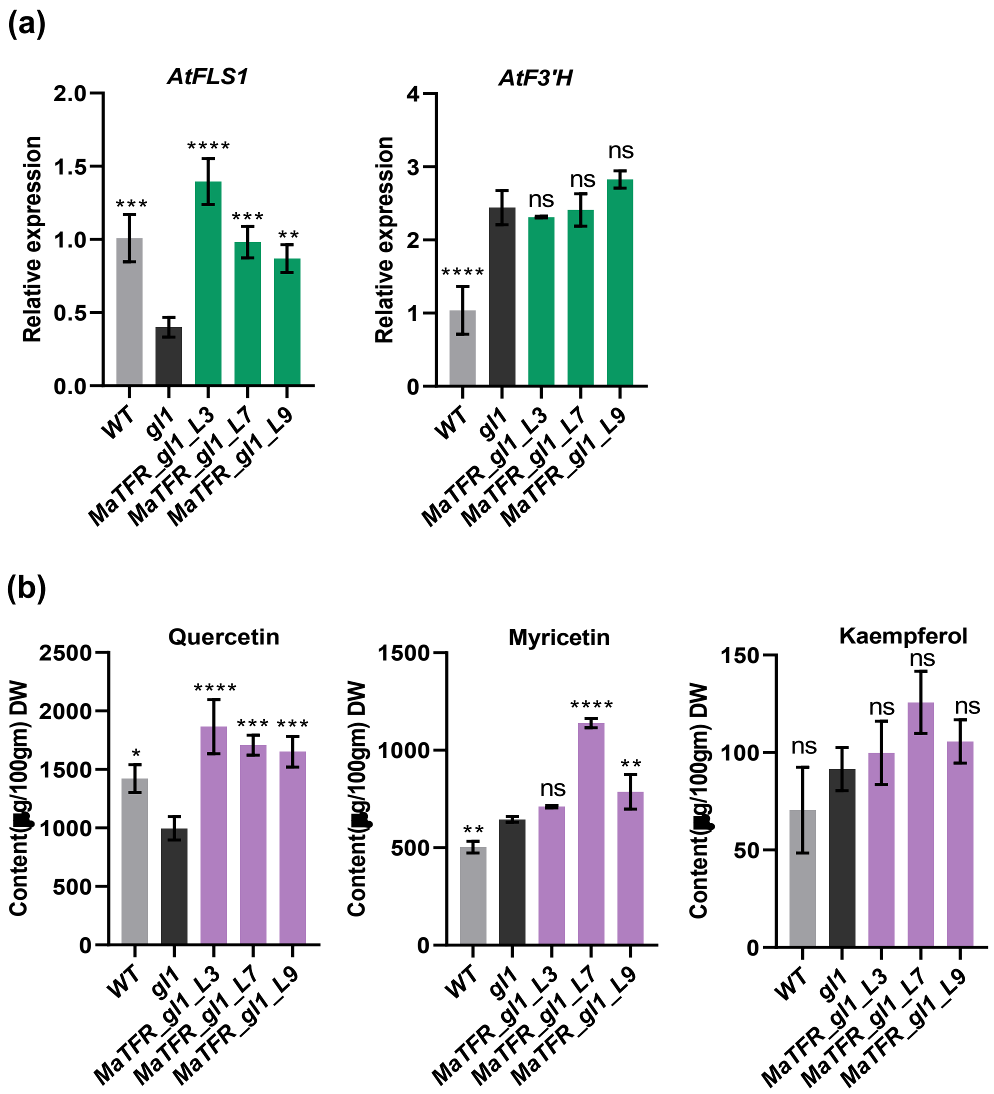
(a)
AtFLS1
Relative expression
2.0
1.5
1.0
0.5
0.0
***
****
***
**
WT
gl1
MaTFR_gl1_L3
MaTFR_gl1_L7
MaTFR_gl1_L9
AtF3'H
Relative expression
4
3
2
1
0
****
ns
ns
ns
(b)
Quercetin
Content(µg/100gm) DW
2500
2000
1500
1000
500
0
*
****
***
***
Myricetin
1500
1000
500
0
**
ns
****
**
Kaempferol
150
100
50
0
ns
ns
ns
ns

Fig. S7

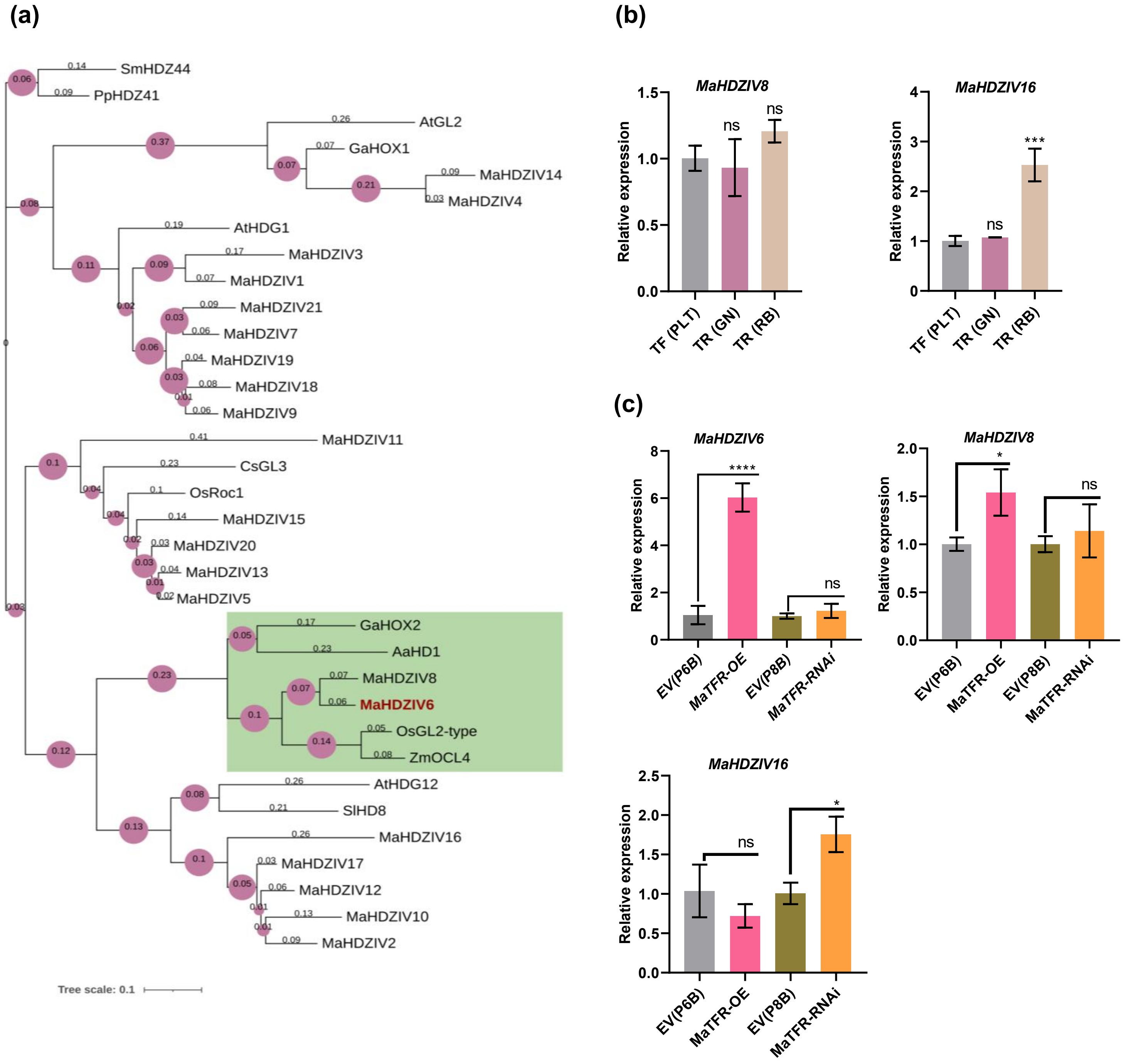
(a)
SmHDZ44
PpHDZ41
AtGL2
GaHOX1
MaHDZIV14
MaHDZIV4
AtHDG1
MaHDZIV3
MaHDZIV1
MaHDZIV21
MaHDZIV7
MaHDZIV19
MaHDZIV18
MaHDZIV9
MaHDZIV11
CsGL3
OsRoc1
MaHDZIV15
MaHDZIV20
MaHDZIV13
MaHDZIV5
GaHOX2
AaHD1
MaHDZIV8
MaHDZIV6
OsGL2-type
ZmOCL4
AtHDG12
SlHD8
MaHDZIV16
MaHDZIV17
MaHDZIV12
MaHDZIV10
MaHDZIV2
Tree scale: 0.1
(b)
MaHDZIV8
MaHDZIV16
Relative expression
TF (PLT)
TR (GN)
TR (RB)
ns
***
(c)
MaHDZIV6
MaHDZIV8
MaHDZIV16
Relative expression
EV(P6B)
MaTFR-OE
EV(P8B)
MaTFR-RNAi
****
*
ns

Fig. S8

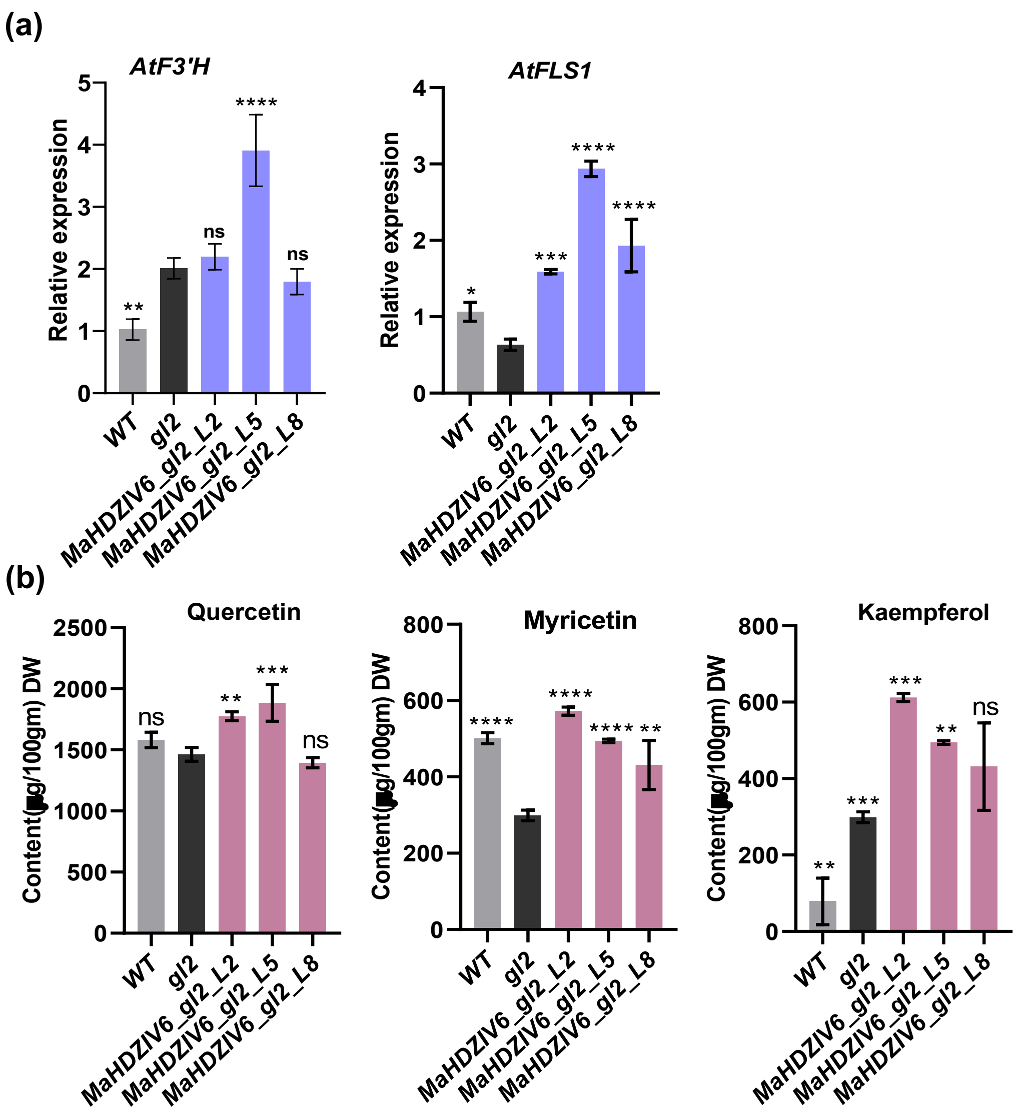
(a)
AtF3'H
AtFLS1
Relative expression
WT
gl2
MaHDZIV6_gl2_L2
MaHDZIV6_gl2_L5
MaHDZIV6_gl2_L8
ns
(b)
Quercetin
Myricetin
Kaempferol
Content(µg/100gm) DW

Fig. S9

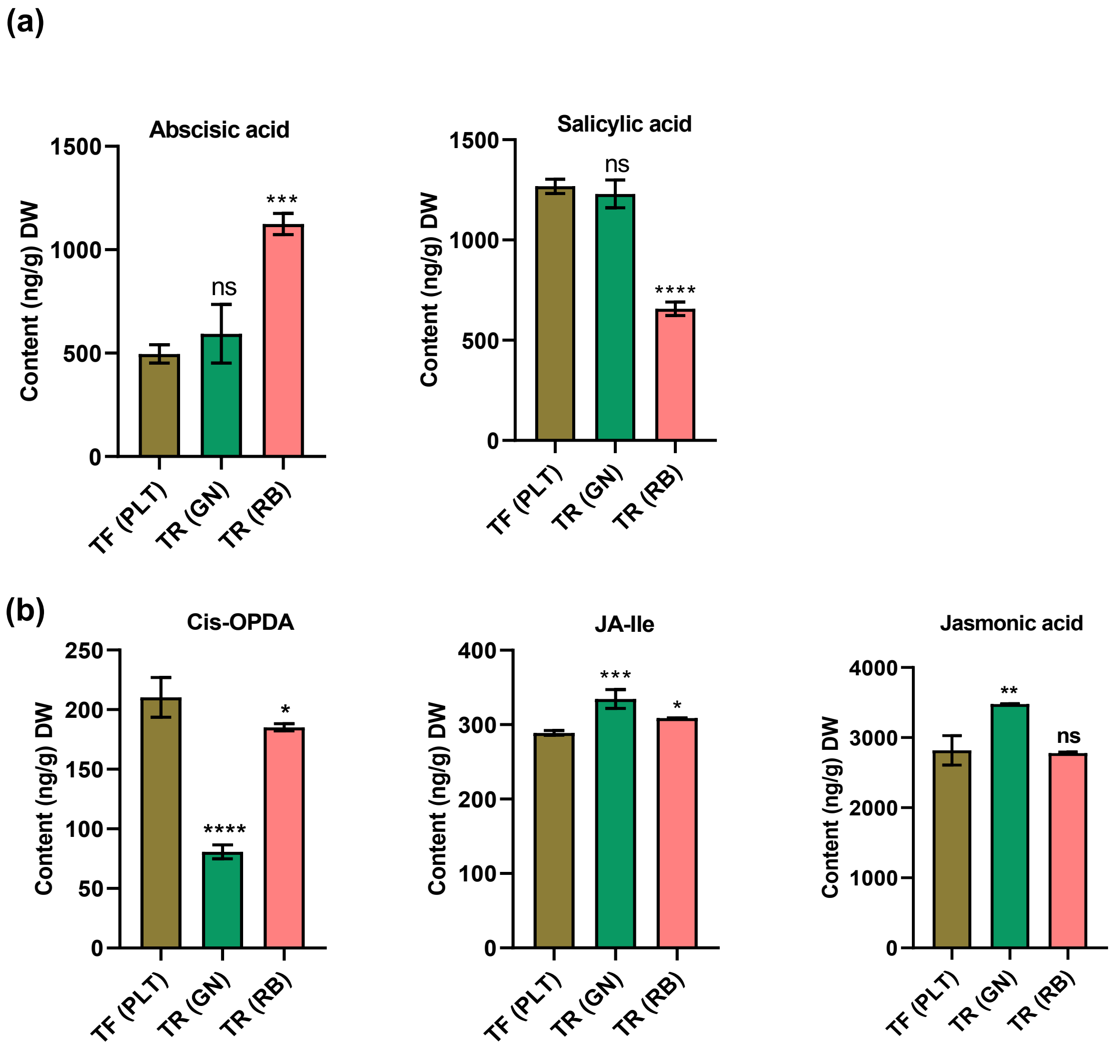
(a)
Abscisic acid
Content (ng/g) DW
1500
1000
500
0
ns
***
TF (PLT)
TR (GN)
TR (RB)
Salicylic acid
Content (ng/g) DW
1500
1000
500
0
ns
****
TF (PLT)
TR (GN)
TR (RB)
(b)
Cis-OPDA
Content (ng/g) DW
250
200
150
100
50
0
****
*
TF (PLT)
TR (GN)
TR (RB)
JA-Ile
Content (ng/g) DW
400
300
200
100
0
***
*
TF (PLT)
TR (GN)
TR (RB)
Jasmonic acid
Content (ng/g) DW
4000
3000
2000
1000
0
**
ns
TF (PLT)
TR (GN)
TR (RB)

Fig. S10

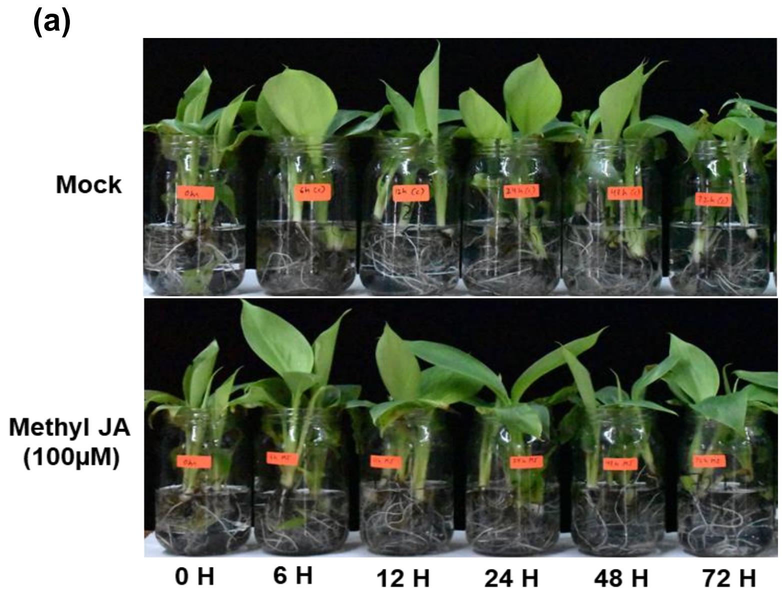

(a)
Mock
Methyl JA
(100µM)
0 H
6 H
12 H
24 H
48 H
72 H


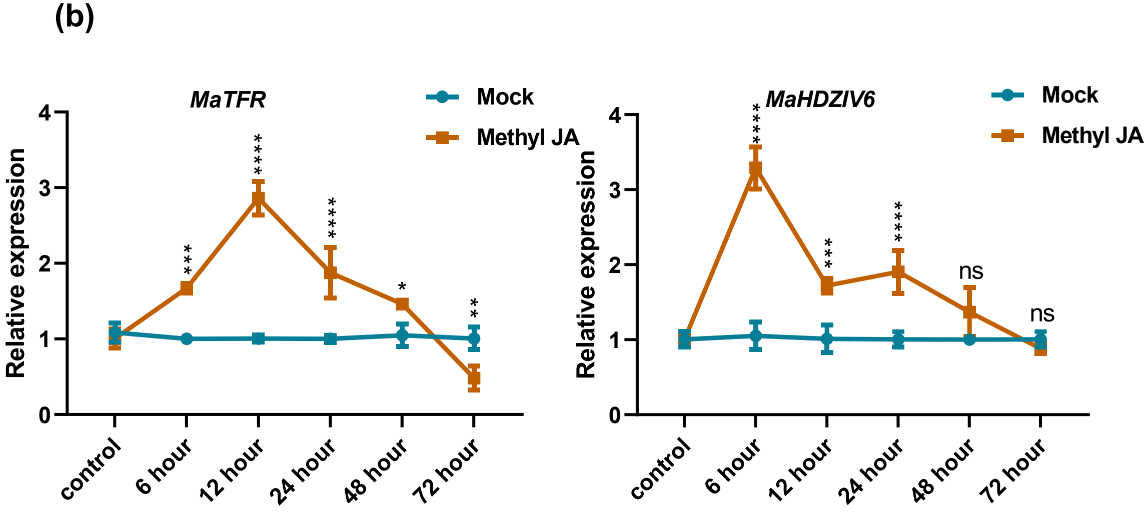

(b)
MaTFR
Mock
Methyl JA
Relative expression
0
1
2
3
4
control
6 hour
12 hour
24 hour
48 hour
72 hour
***
****
****
*
**
MaHDZIV6
Mock
Methyl JA
Relative expression
0
1
2
3
4
control
6 hour
12 hour
24 hour
48 hour
72 hour
****
***
****
ns
ns


Fig. S11

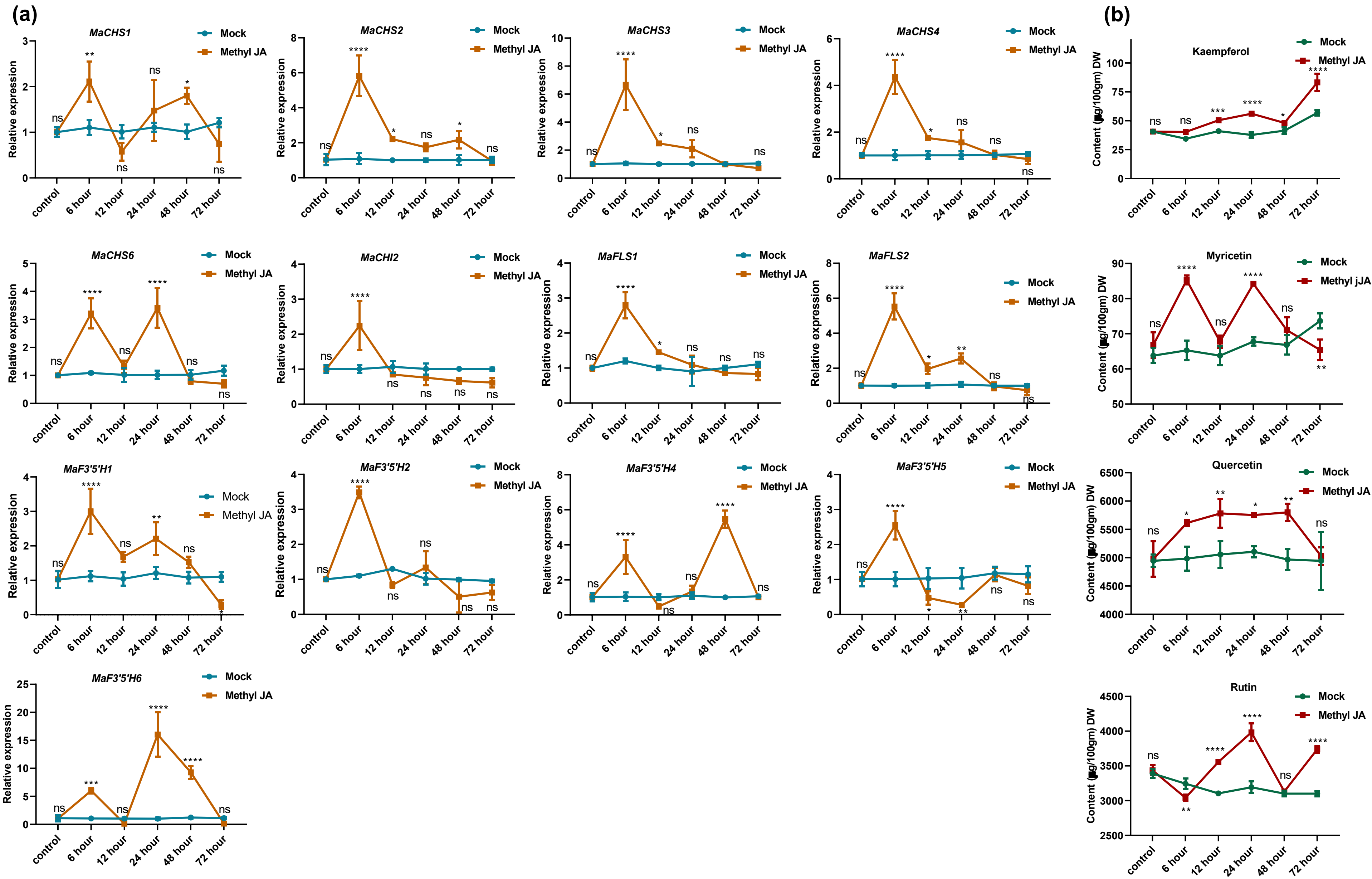
(a)
(b)
MaCHS1
MaCHS2
MaCHS3
MaCHS4
MaCHS6
MaCHI2
MaFLS1
MaFLS2
MaF3'5'H1
MaF3'5'H2
MaF3'5'H4
MaF3'5'H5
MaF3'5'H6
Kaempferol
Myricetin
Quercetin
Rutin
Mock
Methyl JA
Methyl jJA
Relative expression
Content (µg/100gm) DW
control
6 hour
12 hour
24 hour
48 hour
72 hour
ns
*
**
***
****

Fig. S12

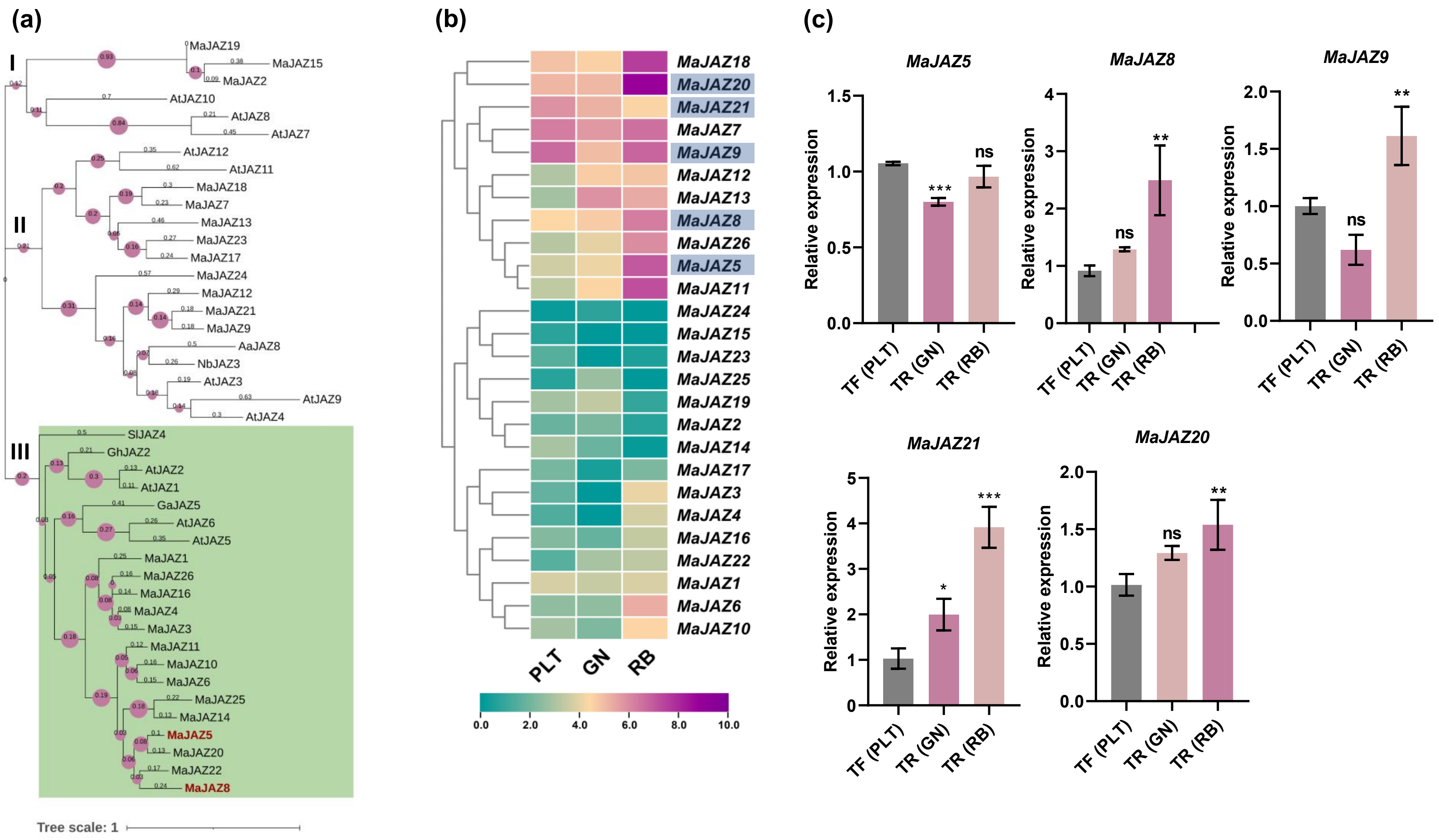
(a)
I
II
III
MaJAZ19
MaJAZ15
MaJAZ2
AtJAZ10
AtJAZ8
AtJAZ7
AtJAZ12
AtJAZ11
MaJAZ18
MaJAZ7
MaJAZ13
MaJAZ23
MaJAZ17
MaJAZ24
MaJAZ12
MaJAZ21
MaJAZ9
AaJAZ8
NbJAZ3
AtJAZ3
AtJAZ9
AtJAZ4
SlJAZ4
GhJAZ2
AtJAZ2
AtJAZ1
GaJAZ5
AtJAZ6
AtJAZ5
MaJAZ1
MaJAZ26
MaJAZ16
MaJAZ4
MaJAZ3
MaJAZ11
MaJAZ10
MaJAZ6
MaJAZ25
MaJAZ14
MaJAZ5
MaJAZ20
MaJAZ22
MaJAZ8
Tree scale: 1
(b)
MaJAZ18
MaJAZ20
MaJAZ21
MaJAZ7
MaJAZ9
MaJAZ12
MaJAZ13
MaJAZ8
MaJAZ26
MaJAZ5
MaJAZ11
MaJAZ24
MaJAZ15
MaJAZ23
MaJAZ25
MaJAZ19
MaJAZ2
MaJAZ14
MaJAZ17
MaJAZ3
MaJAZ4
MaJAZ16
MaJAZ22
MaJAZ1
MaJAZ6
MaJAZ10
PLT
GN
RB
0.0
2.0
4.0
6.0
8.0
10.0
(c)
MaJAZ5
MaJAZ8
MaJAZ9
MaJAZ21
MaJAZ20
Relative expression
TF (PLT)
TR (GN)
TR (RB)
ns
*
**
***

Fig. S13

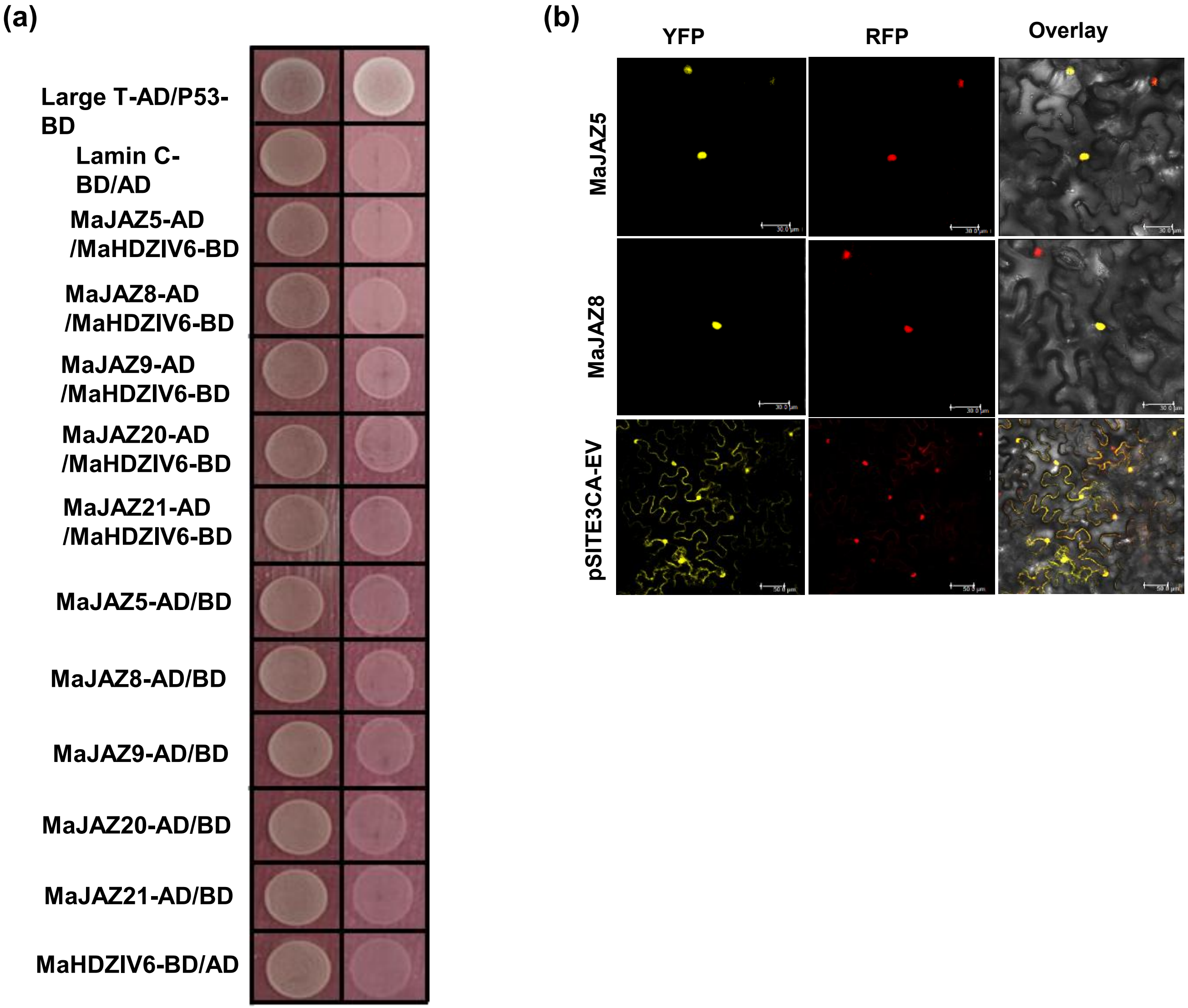
(a)
Large T-AD/P53-BD
Lamin C-BD/AD
MaJAZ5-AD /MaHDZIV6-BD
MaJAZ8-AD /MaHDZIV6-BD
MaJAZ9-AD /MaHDZIV6-BD
MaJAZ20-AD /MaHDZIV6-BD
MaJAZ21-AD /MaHDZIV6-BD
MaJAZ5-AD/BD
MaJAZ8-AD/BD
MaJAZ9-AD/BD
MaJAZ20-AD/BD
MaJAZ21-AD/BD
MaHDZIV6-BD/AD
(b)
YFP
RFP
Overlay
MaJAZ5
MaJAZ8
pSITE3CA-EV

Fig. S14

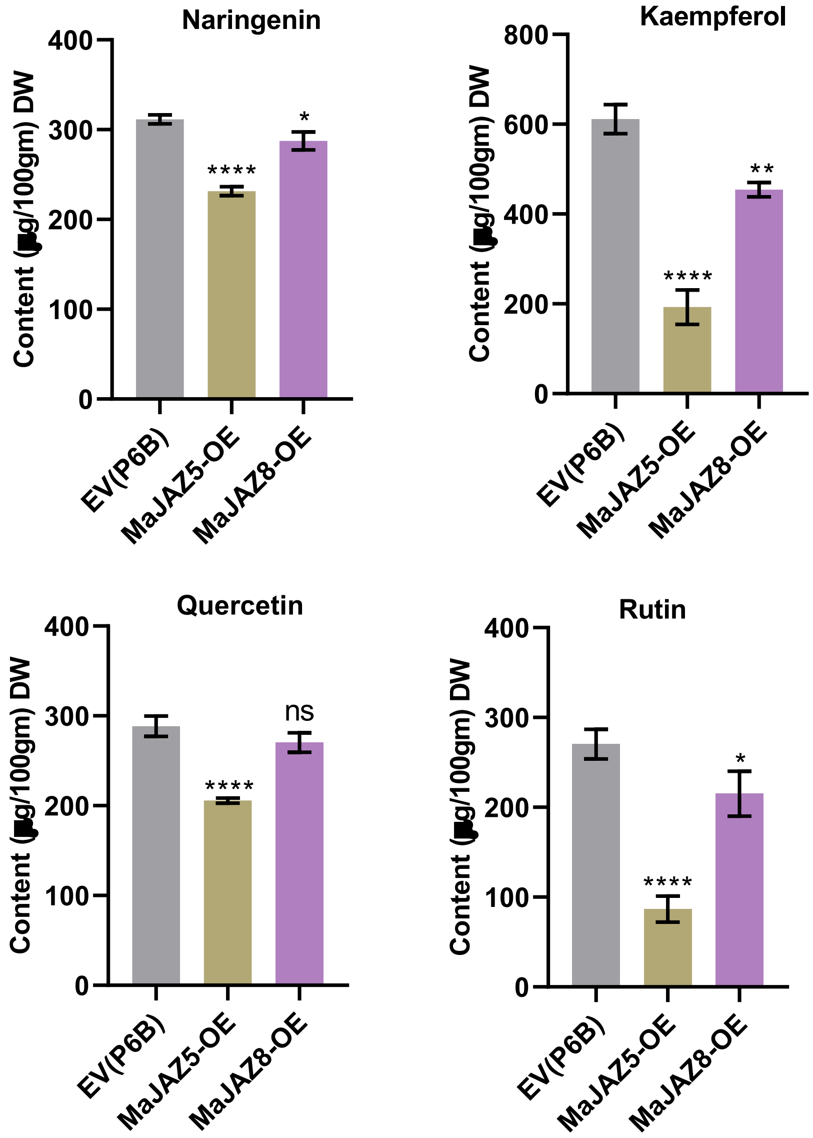

Naringenin
Kaempferol
Quercetin
Rutin
Content (■g/100gm) DW
0
100
200
300
400
600
800
EV(P6B)
MaJAZ5-OE
MaJAZ8-OE
****
**
*
ns


Fig. S15
